# Ketogenesis supports hepatic polyunsaturated fatty acid homeostasis via fatty acid elongation

**DOI:** 10.1101/2024.07.09.602593

**Authors:** Eric D. Queathem, Zahra Moazzami, David B Stagg, Alisa B. Nelson, Kyle Fulghum, Abdirahman Hayir, Alisha Seay, Jacob R. Gillingham, D. Andre d’Avignon, Xianlin Han, Hai-Bin Ruan, Peter A. Crawford, Patrycja Puchalska

## Abstract

Therapeutic interventions targeting hepatic lipid metabolism in metabolic dysfunction-associated steatotic liver disease (MASLD) and steatohepatitis (MASH) remain elusive. Using mass spectrometry-based stable isotope tracing and shotgun lipidomics, we established a novel link between ketogenesis and MASLD pathophysiology. Our findings show that mouse liver and primary hepatocytes consume ketone bodies to support fatty acid (FA) biosynthesis via both de novo lipogenesis (DNL) and FA elongation. Analysis of ^13^C-labeled FAs in hepatocytes lacking mitochondrial D-β-hydroxybutyrate dehydrogenase (BDH1) revealed a partial reliance on mitochondrial conversion of D-βOHB to acetoacetate (AcAc) for cytoplasmic DNL contribution, whereas FA elongation from ketone bodies was fully dependent on cytosolic acetoacetyl-CoA synthetase (AACS). Ketone bodies were essential for polyunsaturated FA (PUFA) homeostasis in hepatocytes, as loss of AACS diminished both free and esterified PUFAs. Ketogenic insufficiency depleted liver PUFAs and increased triacylglycerols, mimicking human MASLD, suggesting that ketogenesis supports PUFA homeostasis, and may mitigate MASLD-MASH progression in humans.

## Introduction

The prevalence of metabolic dysfunction-associated steatotic liver disease (MASLD) and steatohepatitis (MASH) are witnessing a global surge, presently ranking as the most common cause of liver disease in Western countries^1,2^. MASLD is strongly associated with metabolic syndrome, obesity, type 2 diabetes, and dyslipidemia, reflecting its strong link to overall metabolic health. However, the natural history of MASLD progression is complex, and is driven by both systemic and liver-autonomous metabolic dysfunction, leading to the accumulation of excess fat in hepatocytes. Augmentation of hepatic lipid metabolism is also associated with inflammation and cellular stress, which contribute to liver damage and fibrosis, thereby stimulating the progression of MASLD into more severe liver diseases, such as MASH, cirrhosis and hepatocellular carcinoma^2–6^. With most agents failing due to poor efficacy or toxicity, only one agent has been FDA-approved for MASLD treatment^7^, underscoring the need to understand how triacylglycerol (TAG) accumulation in the liver is linked to the heightened risk of developing more severe liver disease^8^.

Steatosis in MASLD is linked to increased hepatic de novo lipogenesis (DNL)^9–14^; however, in early stages of the disease, hepatic lipid disposal pathways, such as very low-density lipoprotein (VLDL) secretion^15,16^ and fat oxidation^17–19^, are also augmented. Therefore, a clearer description of the relationship between hepatic fat synthesis and fat disposal over the natural history of MASLD is needed. Hepatic fat oxidation principally yields mitochondrial acetyl-CoA, which is disposed of either through terminal oxidation in the tricarboxylic acid (TCA) cycle, or through the synthesis of the ketone bodies acetoacetate (AcAc) and D-beta-hydroxybutyrate (D-βOHB). Recent studies have linked diminished fasting hepatic ketogenesis to increased liver steatosis in human MASLD, with concurrent acceleration in fluxes through both the TCA cycle and DNL^19,20^. This is consistent with prior reports comparing human liver biopsies, which showed that hepatic 3-hydroxymethylglutaryl-CoA synthase 2 (HMGCS2) and D-βOHB dehydrogenase 1 (BDH1) protein, two major enzymes in the ketogenesis pathway, decrease in progressive human MASH, with commensurate decreases in fasting circulating ketone body concentrations^21^. Moreover, genetic disruption of hepatic ketone metabolism in rodents has been linked to increased liver injury and fibrosis, through both hepatocyte-autonomous and hepatocyte-hepatic sinusoidal macrophage exchange (i.e., an intrahepatic ketone shuttle), in which liver macrophages oxidize ketone bodies synthesized by neighboring hepatocytes^22–25^.

Ketogenesis-derived AcAc and D-βOHB are effluxed from hepatocytes and serve as alternative fuels in extrahepatic cells that express the enzyme required for ketone body oxidation, succinyl-CoA:3-oxoacid-CoA transferase (SCOT), which is not expressed in hepatocytes^26^ **(Supplementary Figure 1)**. While ketone body oxidation plays a key role in energy metabolism in many high-energy demanding tissues such as the brain, heart, and muscle^26–28^, the anabolic utilization of ketones to support lipid synthesis through acetoacetyl-CoA synthetase (AACS) occurs in adipose and the developing brain^28,29^. In these tissues, ketone bodies may be preferentially sourced for cholesteroneogenesis, but also contribute significantly to fatty acid (FA) biosynthesis^29,30^. Though quantifiable, the physiological relevance of AcAc-dependent lipogenesis in the liver has not yet been elucidated, and thus far, no physiological role for D-βOHB-sourced lipogenesis has been reported in any tissue^29,31–33^.

In this study, we leveraged high resolution and high mass accuracy mass spectrometry (MS)-based ^13^C-stable isotope tracing untargeted metabolomics (ITUM), multidimensional MS-based shotgun lipidomics, and transgenic mouse models lacking BDH1, AACS, or HMGCS2, critical enzymatic mediators of ketone body metabolism, to establish a previously unrecognized role for ketone bodies in FA elongation, polyunsaturated FA (PUFA) homeostasis and the glycerophospholipidome of the liver.

## Results

### The liver metabolizes ketone bodies

While the liver is a net exporter of ketone bodies, we first aimed to determine whether circulating ketone bodies are incorporated into hepatic pathways *in vivo*. To this end, we injected sodium D-[^12^C]βOHB or D-[U-^13^C_4_]βOHB intraperitoneally (IP) (10 μmol/g body weight; n=4 mice) into random-fed wild-type mice, and collected blood and multiple organs for analysis via ^13^C-ITUM (**Supplementary Figure 2A-B**). To survey the metabolic fates of D-[U-^13^C_4_]βOHB-sourced carbon *in vivo*, metabolites were extracted from lyophilized tissues, and analyzed using an optimized ultra-high performance liquid chromatography (UHPLC) coupled to high resolution mass spectrometry (HRMS)-based ^13^C-ITUM pipeline^25,34^. Several dozen chemical features were found to be ^13^C-enriched in all analyzed tissues, and surprisingly, the liver had the second highest number of ^13^C-labeled features of all tissues analyzed **(Supplementary Figure 2C and Supplementary Table 1),** suggesting that a large degree of circulating ketone bodies returned to the liver *in vivo.* To address the role of prospective *in vivo* secondary tracers derived from extrahepatic metabolism of [U-^13^C_4_]ketones, we perfused mouse livers *ex vivo* via the portal vein using oxygenated Krebs-Henseleit buffer supplemented with lactate, pyruvate, and either 1 mM sodium D-[U-^13^C_4_]βOHB or [U-^13^C_4_]AcAc **(Figure 1A).** Both [U-^13^C_4_]ketones were avidly extracted by the liver **(Figure 1B)**, even when accounting for the interconversion through mitochondrial BDH1 **(Figure 1C)**. To study the metabolic fates of ketone bodies in the liver **(Figure 1D),** ^13^C-ITUM was employed, which revealed D-[U-^13^C_4_]βOHB and [U-^13^C_4_]AcAc ^13^C-enriched acetyl-CoA in M+2 at 22.5±9.1% and 17.8±4.2% of the total pool, respectively **(Figure 1E)**. While most amino acids were not found to be ^13^C-enriched **(Supplementary Figure 3A**), several acetylated amino acids **(Supplementary Figure 3B)**, as well as the amino acids glutamate and aspartate were ^13^C-enriched, as were TCA cycle intermediates **(Figure 1F-G and Supplementary Figure 3C)**. The most intense isotopologue of each TCA cycle intermediate was M+2, suggesting that ketone body-derived carbon entered as acetyl-CoA, but did not proceed through multiple turns of the TCA cycle **(Figure 1F and Supplementary Figure 3C)**. Moreover, metabolites linked to the first-span of the TCA cycle (*e.g.,* citrate and glutamate), were ^13^C-enriched nearly 3-fold higher than those from the second-span (*e.g.,* succinate, malate, and aspartate) **(Figure 1G).** [U-^13^C_4_]AcAc ^13^C-enriched the TCA cycle to a greater extent than D-[U-^13^C_4_]βOHB **(Figure 1G).** In both [U-^13^C_4_]ketone perfusions neither pyruvate nor lactate, was significantly ^13^C-enriched **(Figure 1H).** Collectively, these data demonstrate that ketone bodies are rapidly metabolized by the perfused liver and penetrate numerous metabolic pathways, presumably proceeding through the cytosolic acetyl-CoA pool.

**Figure 1.**
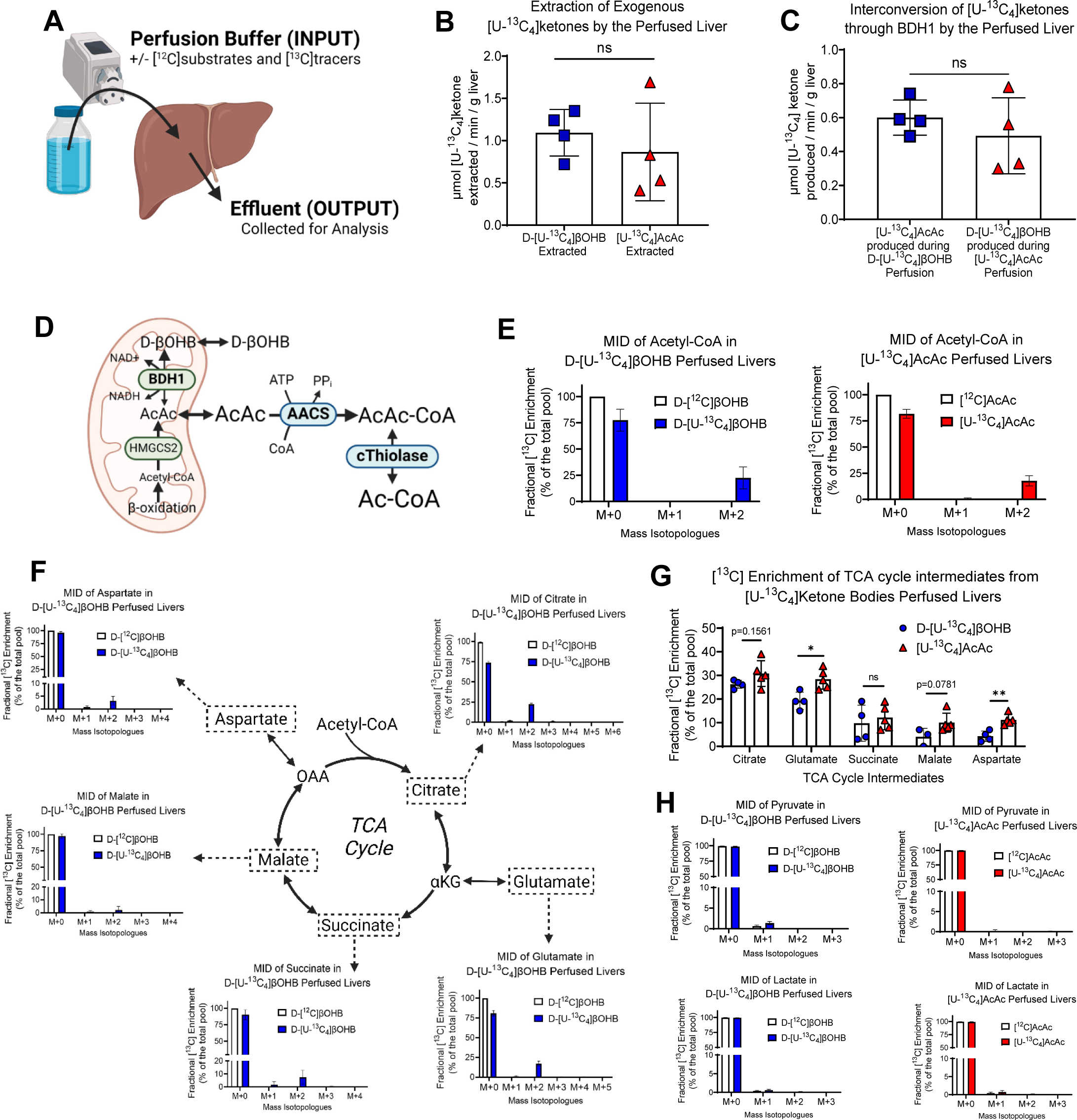
Exogenous ketone bodies are metabolized by the *ex vivo* perfused liver. **(A)** *Ex vivo* portal vein liver perfusions of random-fed mice were performed using oxygenated Krebs-Henseleit buffer supplemented with either 1mM sodium D-[U-^13^C4]beta-hydroxybutyrate (D-[U-^13^C4]βOHB) or 1mM sodium [U-^13^C4]acetoacetate ([U-^13^C4]AcAc) for 30 min. **(B)** Extraction and **(C)** Interconversion of D-[U-^13^C4]βOHB or [U-^13^C4]AcAc by perfused wild-type livers (n=4/group). **(D)** Diagram outlining the known non-oxidative pathway facilitating the entry of ketone-body derived carbon into the metabolome. The cytosolic enzyme acetoacetyl-CoA synthetase (AACS) activates and converts AcAc to AcAc-CoA, which equilibrates with the cytosolic acetyl-CoA (Ac-CoA) pool via cytosolic thiolase. Entry of D-βOHB into metabolism via AACS requires mitochondrial interconversion to AcAc via BDH1. **(E)** Mass isotopologue distribution (MID) of acetyl-CoA and **(F)** tricarboxylic acid (TCA) cycle intermediates or **(G)** fractional [^13^C] enrichment of TCA cycle intermediates from 1 mM sodium D-[U-^13^C4]βOHB or 1 mM sodium [U-^13^C4]AcAc in freeze-clamped and lyophilized livers post 30 minute perfusion analyzed via [^13^C] isotope tracing untargeted metabolomics (ITUM) (n=4-5/group). **(H)** MID of pyruvate (top) and lactate (bottom) from 1 mM sodium D-[U-^13^C4]βOHB (blue) or [U-^13^C4]AcAc (red) in freeze-clamped and lyophilized livers post 30 minute perfusion analyzed via [^13^C] ITUM (n≤5/group). Data are expressed a mean ± SD. Statistically significant differences were determined by Student’s t test, and accepted as p<0.05. *p < 0.05, **p < 0.01, ***p < 0.001, ****p < 0.0001 as indicated. ns = not statistically different.

### Primary hepatocytes metabolize ketone bodies

The liver is a heterogenous organ, containing multiple cell-types^35^, and though hepatocytes do not express SCOT, it is abundantly present in non-parenchymal cells (*e.g.,* macrophages)^25^. Therefore, to test whether hepatocytes metabolize ketone bodies, we isolated primary hepatocytes from random-fed wild-type mice and employed ^13^C-ITUM to trace [U-^13^C_4_]ketone body-derived carbon *in vitro* **(Figure 2A).** We first validated the isolated primary hepatocyte model by confirming that cells were ketogenic and maintained a physiologically normal energy charge and redox state **(Supplementary Figure 4A-C).** RT-qPCR was used to survey parenchymal (*i.e.,* hepatocyte) and non-parenchymal (*i.e.,* non-hepatocyte) gene expression markers, which validated the purity of isolated hepatocyte populations **(Supplementary Figure 4D-E).** Immunoblotting confirmed the presence of HMGCS2 and BDH1 protein, and the absence of SCOT, in isolated primary hepatocytes **(Supplementary Figure 4F).**

**Figure 2.**
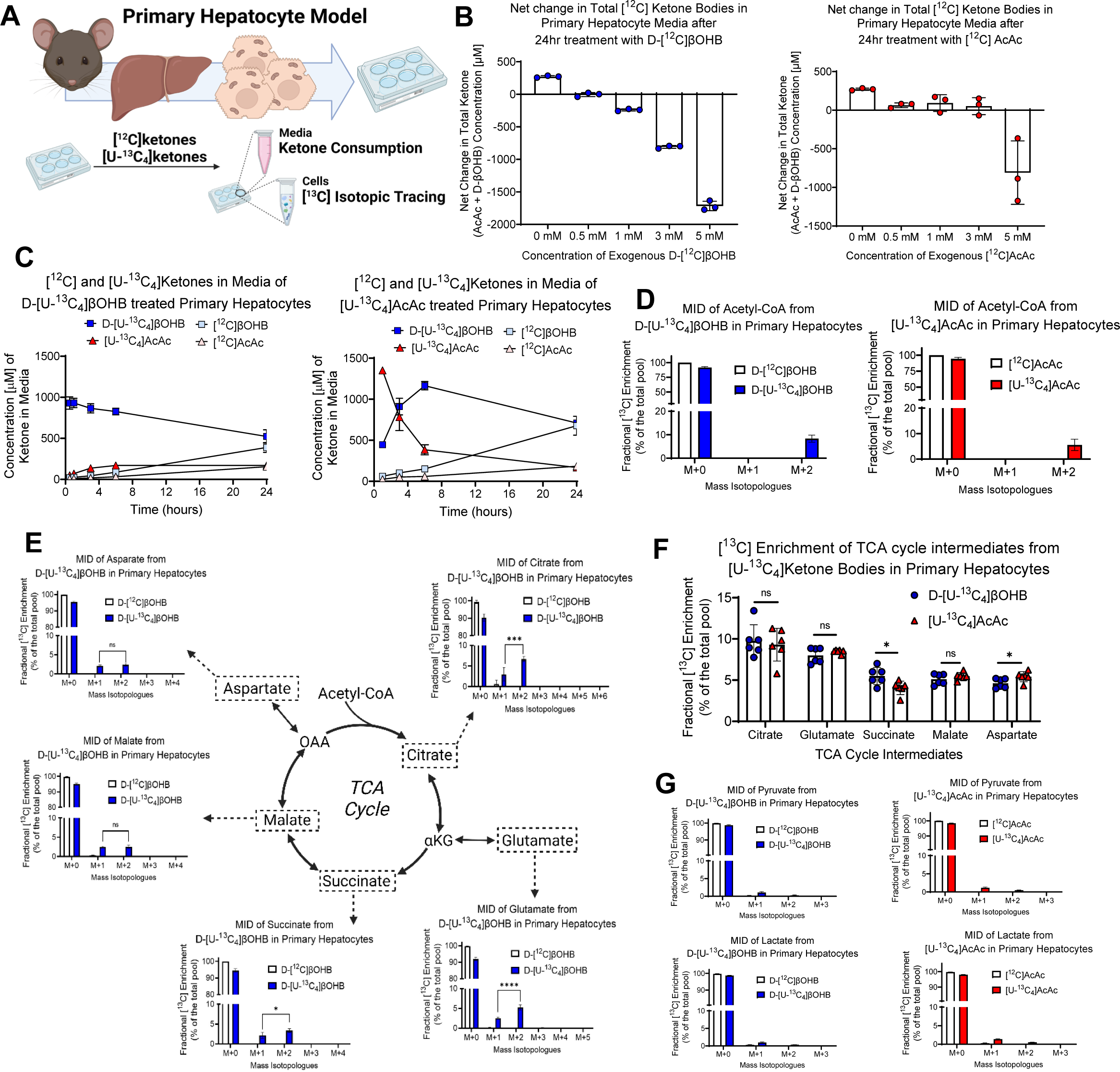
Exogenous ketone bodies are metabolized by primary mouse hepatocytes. **(A)** Primary mouse hepatocytes were treated with media supplemented with 1 mM sodium [U-^13^C4]ketones, incubated for 24 hours, and then media and cells were harvested. **(B)** Net change in total ketone bodies in conditioned media from hepatocytes after 24 hour incubation with increasing concentrations of exogenous naturally-occurring unlabeled ‘[^12^C]’ D-βOHB (blue) or AcAc (red) (n=3/group). **(C)** Concentration (µM) of exogenous [U-^13^C4]ketones, and endogenous produced [^12^C]ketones, in conditioned media over the course of 24 hours from hepatocytes treated with 1 mM sodium D-[U-^13^C4]βOHB (left) or 1 mM sodium [U-^13^C4]AcAc (right) (n=3/group). MID of **(D)** acetyl-CoA, or **(E)** tricarboxylic acid (TCA) cycle intermediates from 1 mM sodium D-[U-^13^C4]βOHB (blue) or 1 mM sodium [U-^13^C4]AcAc (red) in primary hepatocytes after 24 hours analyzed via ^13^C-ITUM (n=6/group). **(F)** Fractional [^13^C] enrichment of TCA cycle intermediates and **(G)** MID of pyruvate (top) and lactate (bottom) from 1 mM sodium D-[U-^13^C4]βOHB (blue) or 1 mM sodium [U-^13^C4]AcAc (red) in primary hepatocytes after 24 hours analyzed via [^13^C] ITUM (n=6/group). Data are expressed a mean ± SD. Statistical differences were determined by Student’s t-test, and accepted as significant if p<0.05. *p < 0.05, **p < 0.01, ***p < 0.001, ****p < 0.0001, as indicated. ns = not statistically different.

Next, we incubated isolated hepatocytes with increasing concentrations of naturally-occurring D-βOHB or AcAc, and found that hepatocytes transitioned from being net producers to net consumers of ketone bodies, in a ketone concentration-dependent manner **(Figure 2B).** Treatment of hepatocytes with 1 mM sodium D-[U-^13^C_4_]βOHB or sodium [U-^13^C_4_]AcAc, revealed a decrease in the media concentration of total [U-^13^C_4_]ketone bodies by 29±7% and 51±2%, respectively, over 24 hours **(Figure 2C).** The M+2 isotopologue of the acetyl-CoA pool was enriched by 8.3±1.4% and 5.6±2.0% from D-[U-^13^C_4_]βOHB and [U-^13^C_4_]AcAc, respectively **(Figure 2D)**, as was the acetyl group of acetylated amino acids **(Supplementary Figure 5A-B).** TCA cycle intermediates were also ^13^C-enriched from both [U-^13^C_4_]ketone bodies **(Figure 2E and Supplementary Figure 5C).** Similar to the perfused liver, the most intense mass isotopologue in each intermediate was M+2 **(Figure 2E and Supplementary Figure 5C)**, and the intermediates linked to the first-span of the TCA cycle (*e.g.,* citrate and glutamate) were ^13^C-enriched to a greater extent than the second-span intermediates (*e.g.,* succinate, malate and aspartate) **(Figure 2F)**. However, unlike the perfused liver, in primary hepatocytes, both [U-^13^C_4_]AcAc and D-[U-^13^C_4_]βOHB ^13^C-labeled TCA cycle intermediates to the same extent **(Figure 2F).** A time course analysis revealed that the TCA cycle became ^13^C-enriched within 30 minutes of [U-^13^C_4_]ketone delivery to cells **(Supplementary Figure 5D),** while pyruvate and lactate enrichment were only ∼1% enriched in M+1 isotopologues after 24 hours **(Figure 2G).** Taken together these data demonstrate an unexpected ability of primary hepatocytes to metabolize ketone bodies, in a SCOT-independent manner, and to selectively partition ketone body-derived carbon into specific metabolic pathways, some of which ultimately proceed through the TCA cycle.

### Ketone bodies contribute to DNL and FA elongation in hepatocytes

Due to the ^13^C-enrichment of the acetyl-CoA pool from [U-^13^C_4_]ketones, in both the perfused liver and primary hepatocytes, we next examined the contribution of ketone body-sourced carbon to FA synthesis in isolated primary mouse hepatocytes **(Figure 3A)**. Both D-[U-^13^C_4_]βOHB and [U-^13^C_4_]AcAc ^13^C-enriched the M+2 isotopologue of malonyl-CoA at 13.0±4.2% and 11.3±1.2% of the total pool, respectively **(Figure 3B).** ^13^C-ITUM was then used to interrogate the three most abundant free FAs (FFAs), as proxies for DNL. Palmitate (total pool within primary hepatocytes was 363.7±42.6 nmol/mg protein), stearate (173.4±38.0 nmol/mg protein) and oleate (336.0±41.7 nmol/mg protein), constituted 24.4±0.7%, 22.6±1.2% and 11.6±2.0% of the total FFA pool, respectively, and were all found to be ^13^C-enriched from both [U-^13^C_4_]ketones, ranging from 1.2±0.3% up to 9.5±1.1% of the total pool **(Figure 3C and Supplementary Figure 6A)**. Because a portion of these FFAs are also derived from exogenously provided fetal bovine serum, the percentage of *de novo* synthesized FFAs from [U-^13^C_4_]ketones is likely higher than that reported here. Furthermore, these FAs were ^13^C-enriched in even numbered mass isotopologues (*e.g*., M+2, M+4, M+6, etc.), consistent with the sequential incorporation of M+2 labeled malonyl-CoA into newly synthesized FAs, via fatty acid synthase (FAS) through the DNL pathway^36^ **(Figure 3A and 3D**). Interestingly, the ^13^C atom percent enrichment (APE) of palmitate, which quantifies the total ^13^C-enrichment of all available carbons in the palmitate pool, from D-[U-^13^C_4_]βOHB was equal to that from [U-^13^C_4_]AcAc, at 1.03±0.15% and 1.25±0.24%, respectively, suggesting that both ketone bodies contributed to DNL equally **(Figure 3E).**

**Figure 3.**
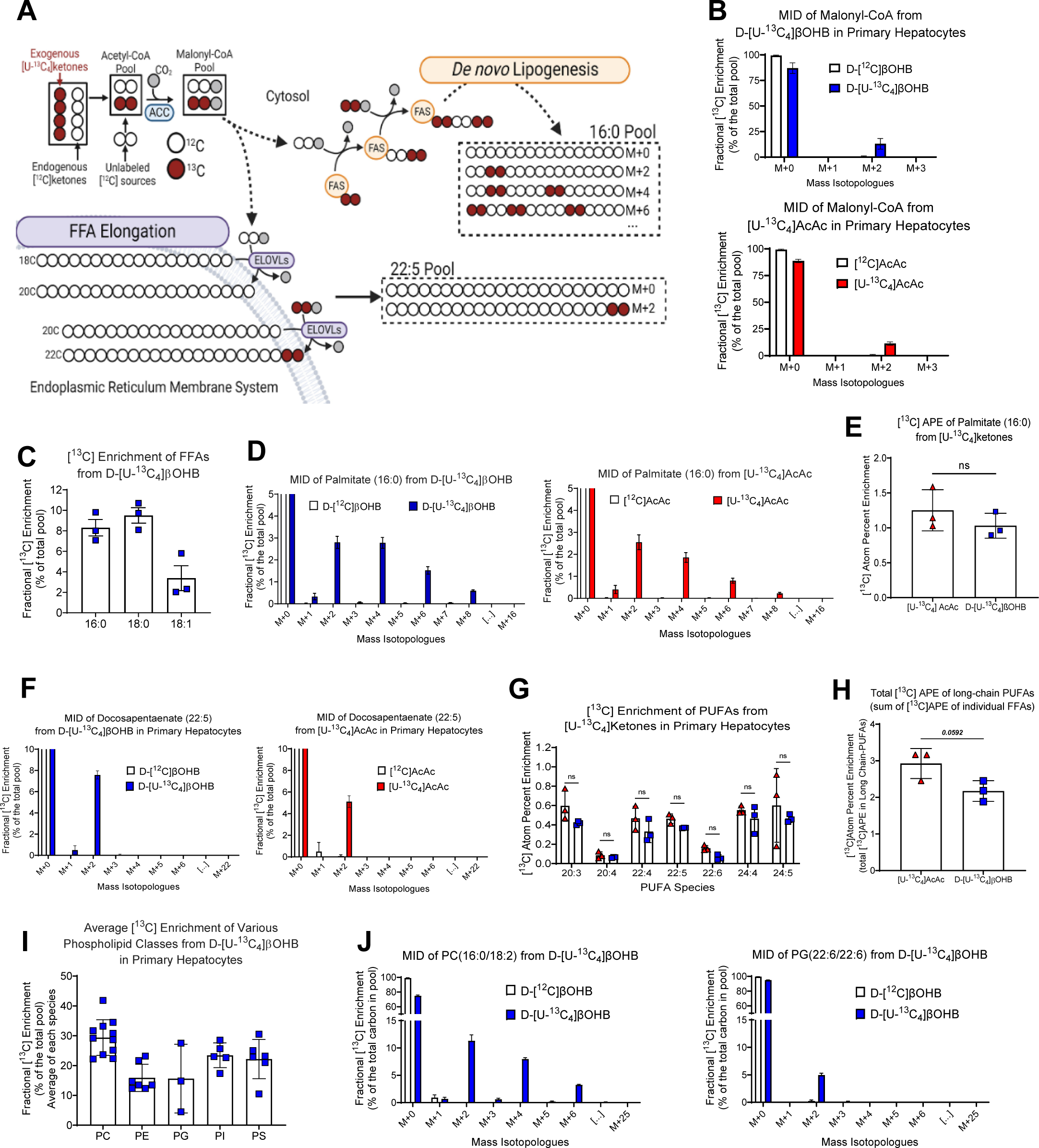
Ketone bodies contribute to lipid biosynthesis in primary hepatocytes via both DNL and FA elongation pathways. **(A)** Scheme outlining [^13^C] ITUM to distinguish DNL and FA elongation. Entry of ketone bodies into lipogenesis proceeds through the cytosolic acetyl-CoA and malonyl-CoA pools, [^13^C]-enriching both pools in M+2 isotopologues. DNL builds palmitate, in a successive manner in the cytosol leading to a characteristic M+2, M+4, M+6… pattern. During FA elongation, malonyl-CoA derived acetyl-units are incorporated by elongases (ELOVL 1-7) localized to the endoplasmic reticulum membrane system, leading to very long chain PUFAs, [^13^C]-enriching PUFA pools solely in M+2. **(B)** Fractional [^13^C] enrichment of malonyl-CoA, and **(C)** FFAs from 1mM sodium D-[U-^13^C4]βOHB (blue) or 1mM sodium [U-^13^C4]AcAc (red) in primary hepatocytes after 24 hours (n=3-6/group). **(D)** MID and **(E)** [^13^C] atom percent enrichment (APE) of palmitate in primary hepatocytes treated with 1mM sodium D-[U-^13^C4]βOHB (blue) or 1mM sodium [U-^13^C4]AcAc (red) (n=3/group). **(F)** MID of docosapentaenate (22:5) and **(G)** [^13^C] APE of various longer chain polyunsaturated fatty acids (PUFAs) from hepatocytes treated with 2 mM sodium D-[U-^13^C4]βOHB (blue) or 1 mM sodium [U-^13^C4]AcAc (red) (n=3/group). **(H)** Summed total of [^13^C] APE of longer chain PUFAs (20-24 carbons) from [U-^13^C4]ketones, normalized to initial ketone concentration (n=3/group). P-value is shown above bar. **(I)** Average fractional [^13^C] enrichment of various phospholipid (PL) species from 1mM D-[U-^13^C4]βOHB, including phosphatidylcholine (PC), phosphatidylethanolamine (PE), phosphatidylglycerol (PG), phosphatidylinositol (PI) and phosphatidylserine (PS) lipid classes. Each point represents enrichment of a unique lipid species (n=6/group). Enrichment of individual PL species found in Supplementary Figure 6. **(J)** MID of the PC (16:0/18:2) (left) and PG (22:6/22:6) (right) from 1mM D-[U-^13^C4]βOHB. Data are expressed a mean ± SD. Statistical differences between AcAc and D-βOHB groups were determined when appropriate student’s t test, and accepted as significant if p<0.05. *p < 0.05; **p < 0.01; ***p < 0.001; ****p < 0.0001; as indicated. ns = not statistically different.

To determine whether ketone bodies also contributed to FA elongation, we scrutinized the ^13^C-enrichment patterns of very long chain PUFAs (20-24 carbons), which do not arise from the DNL pathway, but rather from the elongation of exogenous fats^37,38^. Similar to DNL, malonyl-CoA is the substrate for FA elongation, however, unlike FAS-dependent DNL, only a single malonyl-CoA-derived two-carbon acetyl unit is added to a pre-existing FA, via the actions of one of seven elongases (ELOVL 1-7) localized within the endoplasmic reticulum (ER) membrane system^37,38^. The PUFAs eicosatrienoic acid (20:3), adrenic acid (22:4), docosapentaenoic acid (22:5), docosahexaenoic acid (22:6), tetracosatetraenoic acid (24:4), and tetracosapentaenoic acid (24:5) were all ^13^C-enriched from both [U-^13^C_4_]ketone bodies, ranging between 3.8±0.1% to 11.2±0.8% of their total pools **(Figure 3F-G and Supplementary Figure 6B).** As expected, the ^13^C mass isotopologue distribution (MID) pattern of PUFAs was only ^13^C-enriched in a single M+2 isotopologue, as opposed to the M+2, M+4, M+6… pattern arising from DNL **(Figure 3A and 3F and Supplementary Figure 6B)**. While the ^13^C-APEs of individual PUFAs from D-[U-^13^C_4_]βOHB and [U-^13^C_4_]AcAc were comparable (Student’s t-test, p>0.05, *all)* **(Figure 3G),** within the total pool of measured PUFAs, there was a trend towards a preference for [U-^13^C_4_]AcAc (2.9±0.3% ^13^C-APE) as a substrate for FA elongation over D-[U-^13^C_4_]βOHB (2.2±0.2% ^13^C-APE) (Student’s *t*-test, p=0.0592) **(Figure 3H).** To determine if ketone body-sourced FAs were incorporated into the glycerophospholipidome, we interrogated our ^13^C-ITUM datasets, and found that [U-^13^C_4_]ketone bodies were incorporated extensively into hepatocyte phospholipid (PL) pools, culminating in the ^13^C-enrichment of multiple phosphatidylcholine (PC, 29.4±5.7% of the totoal pool), phosphatidylethanolamine (PE, 15.9±4.2%), phosphatidylglycerol (PG, 15.6±9.4%), phosphatidylinositol (PI, 23.4±3.7%) and phosphatidylserine (PS, 22.2±6%) lipid species **(Figure 3I and Supplementary Figure 6C)**. Furthermore, the ^13^C MIDs of these PLs indicated that ketone bodies contributed to the esterified acyl chains of these lipids via both the DNL and FA elongation pathways **(Figure 3J and Supplementary Figure 6D).** Thus, ketone body-sourced carbon supports both the DNL and FA elongation pathways in hepatocytes, which contribute to PUFA and PL biosynthesis.

### Metabolism of ketone bodies by hepatocytes is partially BDH1-dependent

To determine whether D-βOHB required mitochondrial conversion to AcAc in order to contribute to DNL and FA elongation, we utilized primary mouse hepatocytes isolated from hepatocyte-specific BDH1 KO mice **(Figure 4A)**^39^. We validated the BDH1 KO primary hepatocyte cell culture model by demonstrating the absence of endogenously produced D-βOHB in conditioned media **(Figure 4B),** the absence of BDH1 protein, assessed via immunoblot **(Supplementary Figure 7A)**, and the absence of interconversion of exogenously provided [U-^13^C_4_]ketones **(Supplementary Figure 7B)**. Compared to hepatocytes isolated from wild-type (WT) littermate controls, hepatocytes isolated from BDH1 KO mice showed a 52.0±14.8% (Student’s t-test, p<0.001) diminution in the ^13^C-enrichment of M+2 acetyl-CoA from D-[U-^13^C_4_]βOHB, whereas the reduction in ^13^C-enrichment of malonyl-CoA, was not statistically significant **(Figure 4C)**. As expected, ^13^C-enrichment of acetyl-CoA and malonyl-CoA from [U-^13^C_4_]AcAc was not diminished in BDH1 KO hepatocytes **(Figure 4C)**. Consistent with the labeling of acetyl-CoA, D-[U-^13^C_4_]βOHB-derived fractional ^13^C-enrichment of palmitate was diminished by 62.1±11.6% in the absence of BDH1 (Student’s t-test, p<0.001), but was maintained from [U-^13^C_4_]AcAc (Student’s t-test, p=0.253) (**Figure 4D**-**E**). Additionally, the ^13^C-APE of palmitate derived from D-[U-^13^C_4_]βOHB was decreased by 72.6±9.8% (Student’s t-test, *p* < 0.001) in hepatocytes lacking BDH1, whereas from [U-^13^C_4_]AcAc, the ^13^C-APE of palmitate was only decreased by 25.7±11.0% (Student’s t-test, p=0.086) (**Figure 4F).** Similar results were observed in the stearate and oleate pools, although a 17.6±1.7% diminution in the ^13^C-enrichment of oleate from [U-^13^C_4_]AcAc was observed (Student’s t-test, p<0.001) **(Figure 4D).** The approximate 30% residual ^13^C-APE of palmitate from D-[U-^13^C_4_]βOHB, combined with preservation of D-[U-^13^C_4_]βOHB consumption by BDH1 KO hepatocytes **(Supplementary Figure 7C)**, suggested that BDH1-independent pathways may support D-βOHB contribution to FA anabolism. However, unlike the partial diminution in the ^13^C-enrichment of palmitate from D-[U-^13^C_4_]βOHB in the absence of BDH1, the contribution of D-[U-^13^C_4_]βOHB to very long chain PUFAs was completely eliminated in the BDH1 KO, but was only marginally impaired or unchanged from [U-^13^C_4_]AcAc **(Figure 4G-H and Supplementary Figure 7D).** Taken together, these findings suggest that BDH1 is required for the procession of exogenous D-[U-^13^C_4_]βOHB into FA elongation, but is only partially required for D-[U-^13^C_4_]βOHB-sourced DNL.

**Figure 4.**
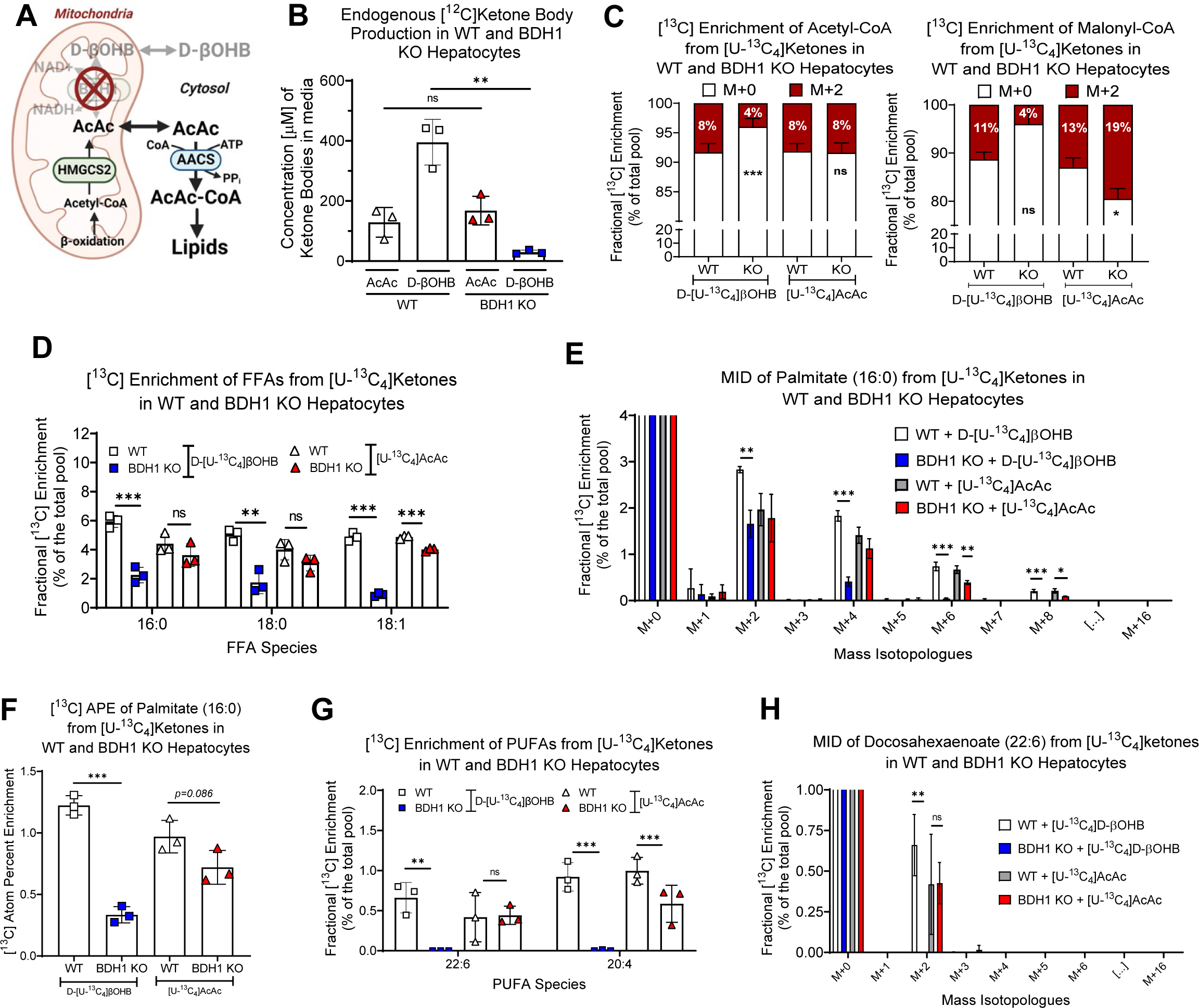
Mitochondrial BDH1 is partially required for exogenous D-βOHB to contribute to lipogenesis. **(A)** Mitochondrial βOHB dehydrogenase 1 (BDH1) interconverts AcAc and D-βOHB in hepatocytes. **(B)** Concentration (µM) of endogenously produced ketones in conditioned media after 24 hours from non-treated WT and BDH1 KO primary mouse hepatocytes (n=3/group). Fractional [^13^C] enrichment of **(C)** acetyl-CoA (left) and malonyl-CoA (right), and **(D)** FFAs in WT and BDH1 KO hepatocytes from 1 mM sodium D-[U-^13^C4]βOHB or 1 mM sodium [U-^13^C4]AcAc after 24 hours (n=3-6/group). **(E)** MID and **(F)** [^13^C] APE of palmitate from WT and BDH1 KO hepatocytes from 1 mM sodium D-[U-^13^C4]βOHB or 1 mM sodium [U-^13^C4]AcAc after 24 hours (n=3/group). **(G)** Fractional [^13^C] enrichment of very long chain PUFAs and **(H)** MID of Docosahexaenoate (22:6) in WT and BDH1 KO hepatocytes from 1 mM sodium D-[U-^13^C4]βOHB or 1 mM sodium [U-^13^C4]AcAc after 24 hours (n=3/group). Data are expressed a mean ± SD. Statistical differences between genotypes were determined by Student’s t test, and accepted as significant if p<0.05. *p < 0.05; **p < 0.01; ***p < 0.001; ****p < 0.0001; as indicated. ns = not statistically different.

### Ketone bodies contribute to FA elongation pathways via an AACS-dependent pathway

AACS is a cytosolic enzyme responsible for the ATP-dependent activation and conversation of AcAc into AcAc-CoA, a step thought to be crucial for the lipogenic fate of ketone bodies^29^. To determine the role of AACS in ketone body-sourced DNL and FA elongation, we first isolated primary hepatocytes from mice with a whole-body genetically engineered loss-of-function in the *Aacs* gene locus **(Figure 5A)**, then confirmed the absence of AACS protein via immunoblot **(Supplementary Figure 8A).** Remarkably, conditioned media from cultured AACS KO hepatocytes accumulated approximately 1.85±0.13 μmoles of total ketone bodies per mg protein, compared to 0.70±0.05 μmoles from hepatocytes isolated from littermate WT controls (Student’s t-test, p<0.001) **(Figure 5B).** While the expression of HMGCS2 and BDH1, two major ketogenic enzymes, were unaltered in AACS KO hepatocytes **(Supplementary Figure 8B),** ^13^C-tracer-dilution modeling of hepatocytes treated with 2 mM sodium D-[U-^13^C_4_]βOHB or 1 mM sodium [U-^13^C_4_]AcAc, confirmed that AACS KO hepatocytes exhibited higher rates of endogenous ketogenesis, as the magnitude of tracer dilution rose by 71.3±7.2% (Student’s t-test, p<0.001) and 55.0±9.3% (Student’s t-test, p=0.003), respectively, compared to WT control cells **(Figure 5C)**.

**Figure 5.**
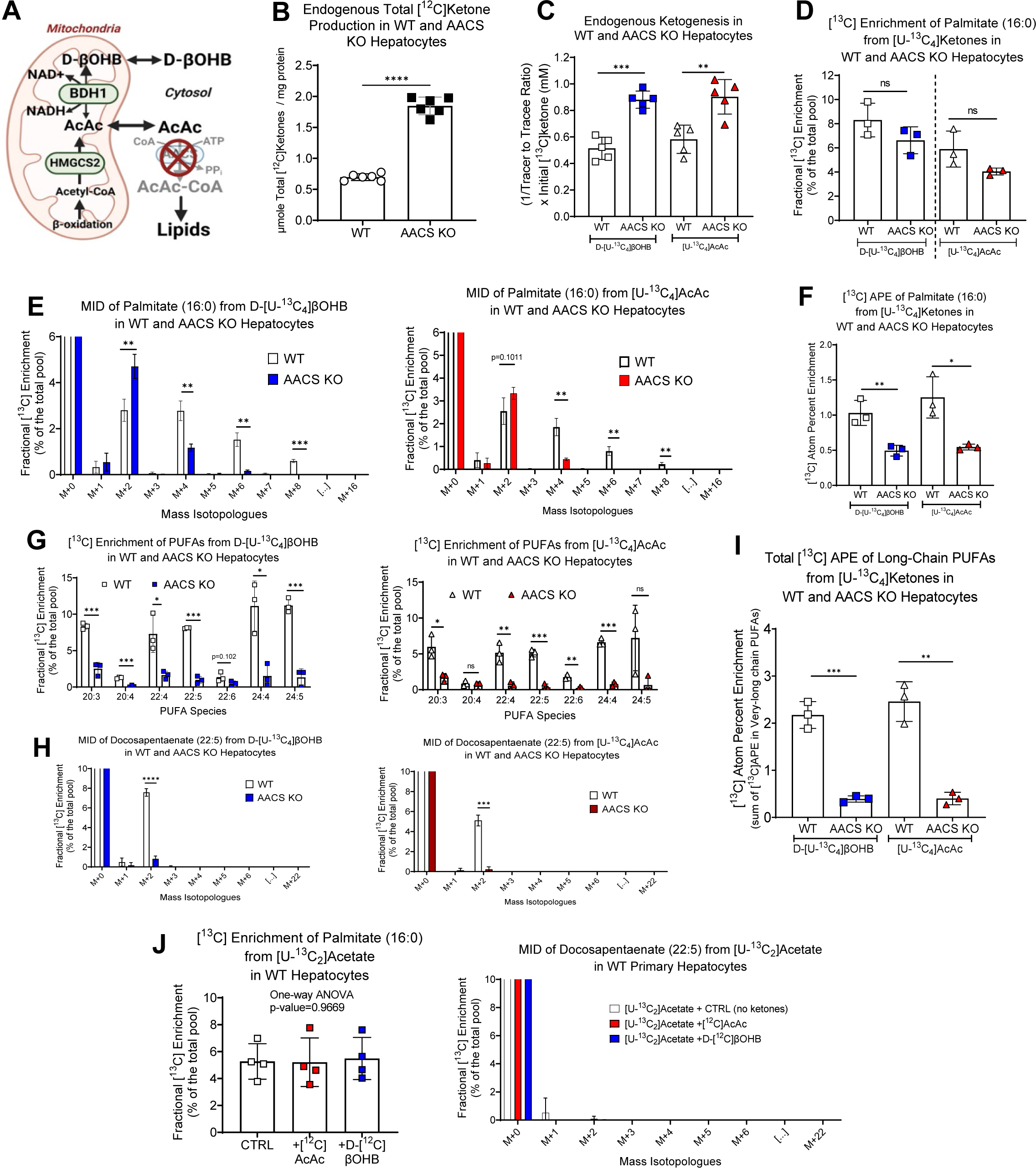
AACS is essential for incorporation of ketone bodies into FA elongation, but is partially dispensable for their entry into DNL. **(A)** AACS activates and converts cytosolic AcAc into AcAc-CoA, and thus facilitates ketone body-derived carbon entry into lipid synthesis. **(B)** Endogenous naturally-occurring unlabeled (‘[^12^C]’) ketones produced in conditioned media after 24 hours in WT and AACS KO primary mouse hepatocytes. Ketone levels were normalized to total protein amount in each sample (n=6/group). **(C)** Endogenous ketogenesis assayed via tracer dilution modeling using 2 mM D-[U-^13^C4]βOHB or 1 mM [U-^13^C4]AcAc after 24 hours in WT and AACS KO primary hepatocytes. The inverse of the Tracer:Tracee ratio (TTR) multiplied by initial [^13^C]ketone concentration was used as surrogate for formal flux quantification. TTR = total [U-^13^C4]ketones / total [^12^C]ketones (n=6/group). **(D)** Fractional [^13^C] enrichment **(E)** MID and **(F)** [^13^C] APE (normalized to initial ketone concentration) of palmitate from 2 mM D-[U-^13^C4]βOHB (blue) or 1 mM [U-^13^C4]AcAc (red) in WT and AACS KO primary hepatocytes (n=3/group). **(G)** Fractional [^13^C] enrichment of very long chain PUFAs and **(H)** MID of the FFA docosapentaenate (22:5) from 2 mM D-[U-^13^C4]βOHB (blue) or 1 mM [U-^13^C4]AcAc (red) in WT and AACS KO primary hepatocytes (n=3/group). **(I)** MID of the FFA palmitate (16:0) and **(J)** FFA docosapentaenate (22:5) in WT hepatocytes incubated with 1mM [U-^13^C2]acetate in the absence (control, CTRL) or presence of naturally-occurring [^12^C] 1 mM D-βOHB or AcAc after 24 hours (n=4/group). Data are expressed a mean ± SD. Statistical differences between genotypes were determined by Student’s t test, and accepted as significant if p<0.05. *p < 0.05; **p < 0.01; ***p < 0.001; ****p < 0.0001; as indicated. ns = not statistically different.

Augmentation of hepatocyte ketogenesis in the absence of AACS raised the hypothesis that the AACS-dependent incorporation of endogenously-produced ketones into lipids via AACS serves as a sink for disposing of ketone bodies within hepatocytes. To test this, we used ^13^C-ITUM to quantify the incorporation of [U-^13^C_4_]ketones into FAs in primary hepatocytes harvested from AACS KO and littermate WT control mice. Interestingly, the fractional ^13^C-enrichment of palmitate from both D-[U-^13^C_4_]βOHB and [U-^13^C_4_]AcAc was not statistically different, in the AACS KO hepatocytes (Student’s t-test, p>0.05 for both) **(Figure 5D)**. However, scrutinizing the ^13^C-MID of palmitate revealed a shift towards lower mass isotopologues in the AACS KO **(Figure 5E).** Consistent with this observation, the ^13^C-APE of palmitate from D-[U-^13^C_4_]βOHB and [U-^13^C_4_]AcAc was decreased by 52.1±10.9% (Student’s t-test, p=0.009) and 56.5±11.6% (Student’s t-test, p=0.015), respectively, in the AACS KO **(Figure 5F)**, suggesting that AACS plays a partial, but not obligate role, in facilitating the entry of ketone bodies into DNL. Conversely, fractional ^13^C-enrichment of very long chain PUFAs from both [U-^13^C_4_]ketones was nearly completely abolished in the AACS KO **(Figure 5G-H)**. Consistent with this finding, the total ^13^C-APE of all PUFAs measured in the AACS KO revealed a marked 82.0±10.0% (Student’s t-test, p<0.001) and 83.7±17.5% (Student’s t-test, p=0.001) diminution from D-[U-^13^C_4_]βOHB and [U-^13^C_4_]AcAc, respectively **(Figure 5I).** This suggested that AACS is required for the incorporation of ketone body-sourced carbon into FA elongation.

These collective data raised the notion that ketone bodies may differentially source malonyl-CoA pools that feed DNL and FA elongation. To test this, we incubated WT primary hepatocytes with 1 mM [U-^13^C_2_]acetate, a classic substrate for modeling DNL, in the absence or presence of unlabeled [^12^C]ketone bodies, then quantified the ^13^C-enrichment of FAs using ^13^C-ITUM. As expected [U-^13^C_2_]acetate ^13^C-enriched the palmitate pool 5.3±0.6%, but the addition of either unlabeled [^12^C]ketone body did not dilute the ^13^C-enrichment of palmitate (one-way ANOVA, p=0.967), suggesting that ketone bodies do not compete with acetate for DNL **(Figure 5J and Supplementary Figure 8C)**. Moreover, [U-^13^C_2_]acetate did not label any of the very long chain PUFA species labeled by [U-^13^C_4_]ketones **(Figure 5J and Supplementary Figure 8D)**, which is consistent with the notion that the malonyl-CoA pool, through which acetate proceeds to enter DNL, could be metabolically distinct from the malonyl-CoA pool contributing to FA elongation.

### AACS is essential for maintaining long-chain PUFA pools in hepatocytes

Given that loss of AACS impairs ^13^C-enrichment of FAs from both [U-^13^C_4_]ketones, we next performed multidimensional MS-based shotgun lipidomics to quantify the impact of losing AACS on the total pool size of lipid species in primary mouse hepatocytes **(Figure 6)**. In AACS KO cells, the total FFA pool size was decreased by 20±9% (Student’s t-test, p=0.047), primarily due to a 29±11% diminution in total unsaturated FFAs (Student’s t-test, p=0.010) **(Figure 6A and Supplementary Table 2).** Moreover, the decrease in unsaturated FFAs in the AACS KO was driven by a 28±8% diminution in the percentage of PUFAs, but not monounsaturated FAs (MUFAs), which were increased by 10±2% **(Figure 6B).** Loss of AACS was further associated with a 40±12% (Student’s t-test, p=0.043) and 17±2% (Student’s t-test, p<0.001) rise in the percentage of long chain FFAs, *i.e.,* those with an acyl chain of 14 or 16 carbons, respectively, but a 19±7% (Student’s t-test, p=0.035), 18±10% (Student’s t-test, p=0.116) and 53±20% (Student’s t-test, p=0.014) decrease in very long chain FAs, *i.e.,* those with an acyl chain of 20, 22 and 24 carbons, respectively **(Figure 6C)**. This coincided with a diminution in multiple elongation indices in the AACS KO **(Figure 6D),** collectively demonstrating that loss of AACS induces a lipidomic profile consistent with an impairment in FA elongation **(Figure 6E).**

**Figure 6.**
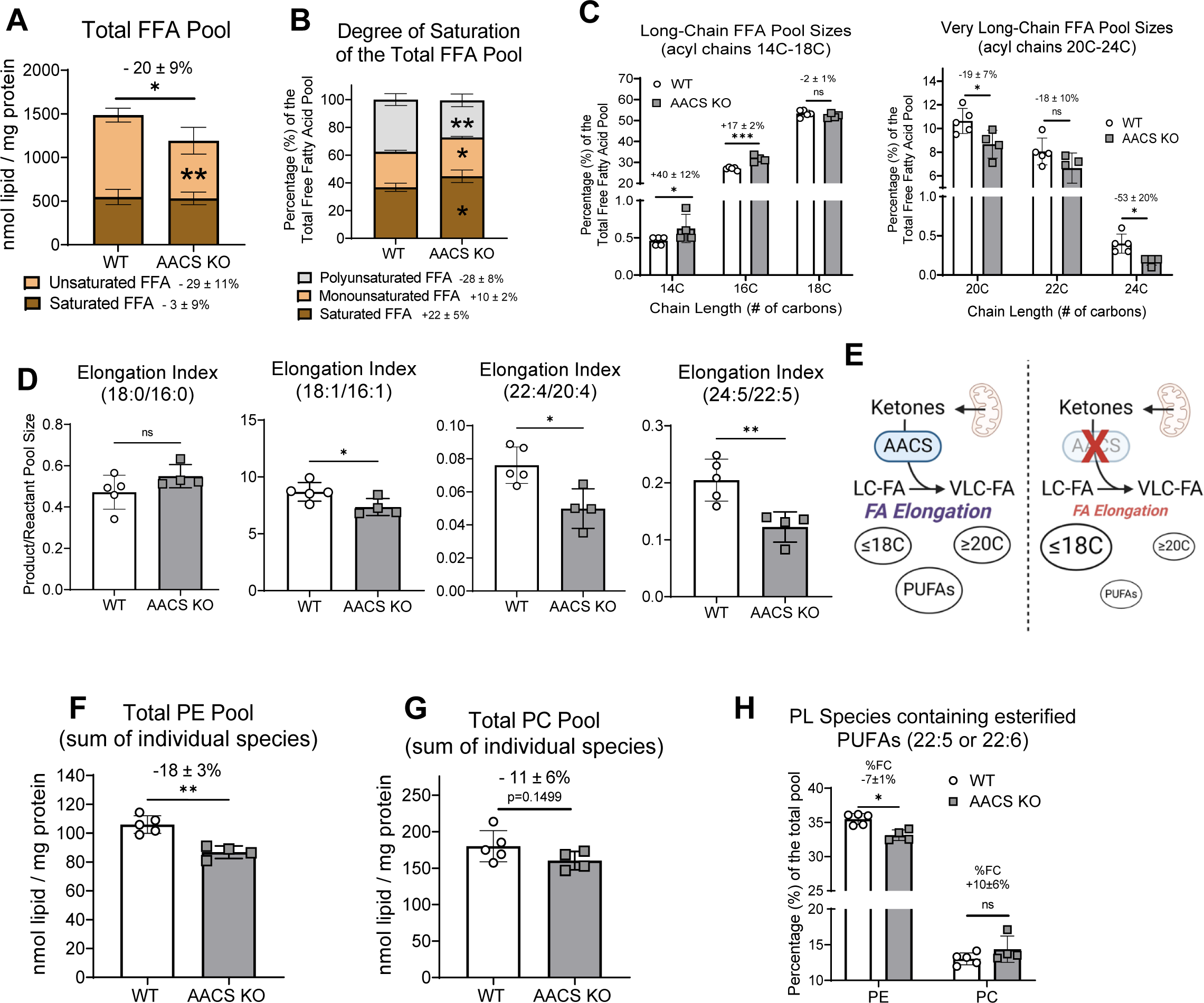
The loss of AACS impairs FA elongation and PUFA biogenesis, disrupting phospholipid metabolism in hepatocytes. **(A)** Total FFA pool size, quantified as the sum of all individual FFAs, broken into saturated (dark brown) and unsaturated (light brown) FFAs in WT and AACS KO primary mouse hepatocytes. Stats bar above the graph shows results of Student’s t-test for total FFAs between WT and AACS KO. Stats symbols inside bars show results of Student’s t-tests for unsaturated and saturated FFAs between WT and AACS KO (n=4-5/group). **(B)** Compositional analysis of the total FFA pool in WT and AACS KO hepatocytes. FFAs were separated into saturated (SFA) (dark brown), monounsaturated (MUFA) (light brown) or polyunsaturated (PUFA) (gray) (n=4-5/group). **(C)** The distribution of acyl chain length within the total FFA pool of WT and AACS KO hepatocytes, broken down into long-chain FAs (18 carbons and below) (left) and very-long-chain FAs (20 carbons and above) (right) (n=4-5/group). **(D)** Elongation indices in WT and AACS KO hepatocytes calculated as the pool size of elongated product divided by precursor (n=4-5/group). **(E)** AACS facilitates the entry of ketone bodies into FA elongation, which supports PUFA biosynthesis. AACS deficient hepatocytes display a FFA lipidomic signature consistent with an impairment in FA elongation encompassing an accumulation of long-chain FAs and a decrease in very-long-chain FAs, coinciding with a depletion in PUFAs. **(F)** Total PE or **(G)** total PC pool sizes calculated as the sum of all individual PE or PC species, respectively, detected (n=4-5/group). **(H)** Percentage of the PE and PC pools enriched in the esterified PUFAs docosahexaenoate (22:6) or docosapentaenoate (22:5), calculated as the sum of all individual species containing esterified 22:6 or 22:5 fatty acids, normalized to the total PL pool (n=4-5/group). Main genotype and PL effects, as well as statistical interactions, were determined via two-way ANOVA. Select results of post-hoc Tukey’s post hoc t-test presented by * on the graph. Data are expressed as mean ± SD. Statistical differences between genotypes were determined by Student’s t-tests, unless otherwise stated and accepted as significant if p<0.05. *p < 0.05; **p < 0.01; ***p < 0.001; ****p < 0.0001; as indicated. ns = not statistically different. Percent change and error propagation are given as X±Y% on graphs when relevant.

To determine the associated impact on esterified PUFAs within PL pools, we performed shotgun lipidomics, which revealed an 18±3% decrease in total hepatocellular PEs (Student’s t-test, p = 0.001) **(Figure 6F, Supplementary Table 3)**, but a non-statistically significant 11±6% decrease in total PCs (Student’s t-test, p=0.150) **(Figure 6G and Supplementary Table 4),** resulting in no net change in the total PC:PE ratio in the AACS KO **(Supplementary Figure 8E).** Loss of AACS slightly increased total TAGs by 14±6% (Student’s t-test, p=0.110) (**Supplementary Figure 8F and Supplementary Table 5).** Investigation of the 22:5- and 22:6-esterified PL pools revealed a phospholipid class-specific role for ketogenesis (two way ANOVA, p=0.004), in which PE pools containing esterified PUFAs decreased by 7±1% in the AACS KO hepatocytes (Tukey’s post hoc test, *p* = 0.032), whereas PC pools containing esterified PUFAs were not different, but tended to increase 10±6% (Tukey’s post hoc test, p=0.312) **(Figure 6H)**. Collectively, these data indicate a role for the AACS-dependent sourcing of ketone bodies to FA elongation within hepatocyte PUFA homeostasis.

### Endogenous ketogenesis remodels PUFAs and the liver glycerophospholipidome via FA elongation

The studies presented above demonstrated in primary hepatocytes that exogenous ketone bodies play a role in remodeling PUFAs and the glycerophospholipidome. To elucidate the role of endogenous ketogenesis in supporting hepatic lipid anabolism *in vivo,* we employed multidimensional MS-based shotgun lipidomics to quantify hepatic lipid species in overnight-fasted mice lacking HMGCS2, the rate limiting step for ketogenesis, selectively in hepatocytes **(Figure 7A)**. We first validated that HMGCS2 was eliminated using immunoblotting and immunohistochemistry **(Supplementary Figure 9A),** and that ketogenesis was disrupted, by demonstrating that hepatocyte-specific HMGCS2 KO mice failed to elevate circulating ketone bodies during fasting **(Supplementary Figure 9B)**. Shotgun lipidomics revealed that livers of HMGCS2 KO mice exhibited a 68±13% increase in total FFAs (Student’s t-test, p=0.014), which was driven by an elevation in both saturated (Student’s t-test, p=0.011) and unsaturated FFAs (Student’s t-test, p=0.016) (**Supplementary Table 6 and Figure 7B)**. Compositional analysis of the bulk FFA pool revealed an 8±3% diminution in the percentage of PUFAs (Student’s t-test, p=0.044), while MUFAs increased by 40±5% (Student’s t-test, p<0.001) (**Figure 7C).** Additionally, long chain FAs of acyl chain length 14, 16 and 18 carbons, were either elevated or unchanged **(Figure 7D)**, while very long chain FAs of acyl chain length 20, 22 and 24 carbons were diminished by 27±8% (Student’s t-test, p=0.004), 36±20% (Student’s t-test, p=0.043), and 31±38% (Student’s t-test, p=0.350), respectively **(Figure 7E).** Lower elongation indices in livers of HMGCS2 KO mice were also observed **(Figure 7F),** collectively supporting an impairment in FA elongation *in vivo* in the absence of hepatic ketogenesis. Quantification of glycerophospholipids in livers of HMGCS2 KO mice revealed no significant differences in total PEs, total PCs, or the total PC:PE ratio **(Supplementary Table 7-8, Figure 7G-I)**. However, analysis of the esterified PUFA fraction within PE and PC pools revealed a PL class-specific role for ketogenesis in the liver (two-way ANOVA, p=0.001) **(Figure 7J)**. Specifically, livers from HMGCS2 KO mice showed a 26±3% diminution in PE species that were specifically enriched in PUFA acyl chains (Tukey’s post hoc t-test, *p* <0.001), with no significant change in the PUFA-esterified PC pool (Tukey’s post hoc t-test, p=0.412) **(Figure 7J).** These observations were all analogous to those in primary hepatocytes harvested from mice lacking AACS. Finally, livers of mice lacking HMGCS2 selectively in hepatocytes exhibited a 2.4±0.2-fold rise in total TAGs *in vivo* **(Supplementary Table 9, Figure 7K)** (Student’s t-test, p <0.001), demonstrating an integral role for hepatic ketogenesis in regulating hepatic steatosis. Taken together, these data suggest that hepatic ketogenesis plays a role in coordinating DNL and FA elongation, to support PUFA homeostasis and the glycerophospholipidome of the liver.

**Figure 7.**
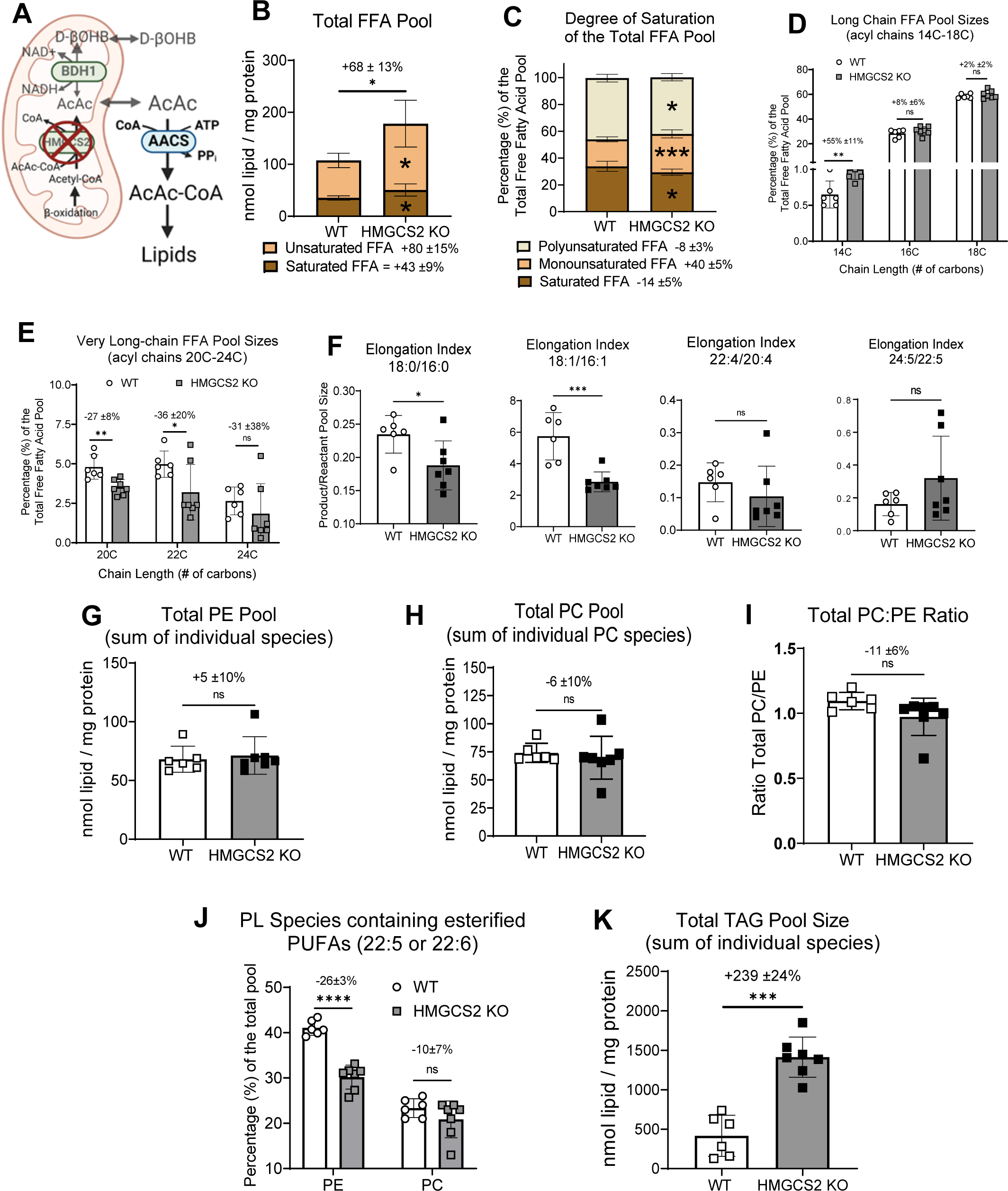
The loss of hepatocyte specific-HMGCS2 *in vivo* impairs FFA elongation, dysregulates PUFA and phospholipid metabolism, and induces hepatic steatosis. **(A)** Mitochondrial HMG-CoA synthase 2 (HMGCS2) is the rate limiting step for ketogenesis in hepatocytes. WT and hepatocyte-specific HMGCS2 null mice were fasted overnight, then multidimensional mass spectrometry-based shotgun lipidomics was used to study the lipidome of the liver. **(B)** Total FFA pool size, quantified as the sum of all individual FFAs, broken into saturated (dark brown) and unsaturated (light brown) FFAs in overnight fasted male WT and HMGCS2 KO livers. Stats bar above graph shows results of student t-test for total FFA between WT and KO. Stats symbol inside bars shows results of student t-test for saturated or unsaturated FFAs between WT and KO (n=6-7/group). **(C)** Compositional analysis of the total FFA pool in WT and HMGCS2 KO livers. FFAs were saturated (SFA) (dark brown), monounsaturated (MUFA) (light brown) or PUFA (gray) (n=6-7/group). **(D)** The distribution of acyl chain length within the total FFA pool of WT and HMGCS2 KO hepatocytes, broken down into long chain FAs (18 carbons and less) (n=6-7/group). **(E)** The distribution of acyl chain length within the total FFA pool of WT and HMGCS2 KO hepatocytes, broken down into very long chain FAs (20 carbons and above) (n=6-7/group). **(F)** Elongation indices in WT and HMGCS2 KO hepatocytes calculated as the pool size of elongated product divided by precursor (n=6-7/group). **(G)** Total PE and **(H)** PC pool sizes calculated as the sum of all individual PE or PC pools, respectively, detected in WT and HMGCS2 KO livers (n=6-7/group). **(I)** The PC:PE ratio calculated by dividing the total PC pool by the total PE pool in WT and HMGCS2 KO livers (n=6-7/group). **(J)** Percentage of the PE and PC pools enriched in the esterified PUFAs docosahexaenoate (22:6) or docosapentaenoate (22:5), calculated as the sum of all individual species containing esterified 22:6 or 22:5 fatty acids, normalized to the total PL pool (n=4-5/group). Main genotype and PL effects, as well as statistical interactions, were determined via 2×2 ANOVA. Main effect of genotype (G) (p<0.0001), main effect of PL (p<0.0001), and GxPL interaction (p=0.0010). Select results of post-hoc Tukey’s t-test presented by * on the graph. **(K)** Total TAG calculated by summing together the pools of each individual TAG species detected (n=6-7/group). Data are expressed as mean ± SD. Statistical differences between genotypes were determined by Student’s t-tests, unless otherwise stated and accepted as significant if p<0.05. *p < 0.05; **p < 0.01; ***p < 0.001; ****p < 0.0001; as indicated. ns = not statistically different. Percent change and error propagation are given as X±Y% on graphs when relevant.

## Discussion

In this study, we leveraged ^13^C-ITUM, multidimensional MS-based shotgun lipidomics, and genetically engineered mouse models to comprehensively map [U-^13^C_4_]ketone body-derived carbon throughout the hepatocyte metabolome and lipidome. Collectively, these novel findings have shifted our view of ketone body metabolism in hepatocytes, by showing that: (1) ketone body-derived carbon can proceed in a SCOT-independent manner into the hepatic metabolome, (2) D-βOHB serves as a substrate for FA synthesis in hepatocytes, (3) ketone bodies contribute to FA synthesis via both the DNL and FA elongation pathways, (4) ketogenesis is linked to PUFA metabolism, in an AACS-dependent manner, and (5) endogenous ketone body-derived carbon supports lipid homeostasis and metabolic health of the liver. Prior studies have linked ketone bodies to human health and metabolic diseases, including MASLD-MASH^19–21,23–27^. However, because hepatocytes robustly express the rate-limiting ketogenic enzyme HMGCS2^40–42^, and lack the fate-committing ketolytic enzyme SCOT^43^, conventionally the liver is viewed solely as a producer of ketone bodies. Prior work in liver macrophages provided nuanced to this view, wherein hepatic macrophage cells were shown to dispose of ketones produced in neighboring hepatocytes^25,44^. No study, however, has yet to reveal a physiological role for ketone body disposal within hepatocytes themselves.

Exogenous D-[U-^13^C_4_]βOHB and [U-^13^C_4_]AcAc each contribute to hepatic metabolism via the acetyl-CoA pool, as evidenced by the ^13^C-enrichment of M+2 acetyl-CoA in both perfused livers and primary hepatocytes. Prior work showed that liver macrophages could oxidize ketones produced by neighboring hepatocytes^25,44^, and thus in the perfused liver, the ^13^C-enrichment of TCA cycle intermediates was expected. While in perfused livers [U-^13^C_4_]AcAc preferentially ^13^C-enriched TCA cycle intermediates, compared to D-[U-^13^C_4_]βOHB, this preferential labeling by [^13^C]AcAc was not observed in primary hepatocytes, suggesting SCOT-independent pathways for allowing the procession of [U-^13^C_4_]ketone body-derived carbon. Metabolic pathways leading to SCOT-independent [U-^13^C_4_]ketone labeling of the TCA cycle and their biological importance are unknown. Pioneering studies previously indicated that ketone body-derived carbon supports biosynthetic reactions, such as DNL and cholesterogenesis in the liver^30–33,45–51^; however these studies only focused on the intact liver, and only traced [^14^C]AcAc-derived carbon. Other cell types, including cells in development and tumor cells, also source ketone bodies to lipogenesis, but the biological significance of these pathways are unknown^29^. Livers of mice lacking BDH1 or AACS provided mechanistic insight into the pathways through which ketone bodies contributed to hepatocyte lipid anabolism. Mitochondrial conversion of exogenous D-βOHB to AcAc via BDH1 was required for ∼70% of the sourcing of D-βOHB to DNL in primary hepatocytes, suggesting that an alternative medium-chain acyl-CoA synthetase exists which can also support the contribution of D-βOHB to DNL. However, sourcing of D-βOHB into FA elongation was fully dependent on BDH1-mediated conversion of exogenous D-βOHB to AcAc. Sourcing of AcAc, on the other hand, to either DNL or FA elongation, did not require BDH1, indicating that AcAc contributes to the malonyl-CoA pool(s) directly, without an intermediate step through D-βOHB, whereas D-βOHB can proceed either through an AcAc intermediate or through an independently-generated intermediate, such as D-βOHB-CoA, which has been proposed in hepatoma cells^52,53^.

The contribution of exogenous D-βOHB to DNL was diminished by ∼50% in the absence of AACS, but its sourcing to PUFAs was nearly completely abrogated in hepatocytes that lacked AACS. Partial dependence on AACS was observed for AcAc tracing to DNL, further underscoring the existence of an alternative medium-chain acyl-CoA synthetase, which serves at least a partially redundant role. There are multiple medium-chain acyl-CoA synthetases for which D-βOHB or AcAc could serve as lower-affinity substrates, thereby facilitating the ATP-dependent synthesis of D-βOHB-CoA or AcAc-CoA, respectively^54^. These collective findings are consistent with an overall model of ketone body-derived FA synthesis (**Figure 8**), in which (a) multiple pools of acetyl-CoA and/or malonyl-CoA are independently sourced from ketone bodies to support DNL and FA elongation, and (b) AcAc is an obligate intermediate for the contribution of ketone bodies to FA elongation, in an AACS-dependent manner. In addition to mitochondrial BDH1, cytosolic BDH2 could also facilitate the interconversion of D-βOHB and AcAc; however, because BDH2 has a K_m_ ∼10-fold higher for D-βOHB and AcAc than BDH1, and because there is virtually no production of D-βOHB by livers or hepatocytes derived from hepatocyte-specific BDH1 KO mice^39,55–57^, this is unlikely. Nonetheless, we cannot rule out the possibility that BDH2-dependent D-βOHB production is directed to DNL and/or FA elongation, rendering no residual D-βOHB available for efflux.

**Figure 8.**
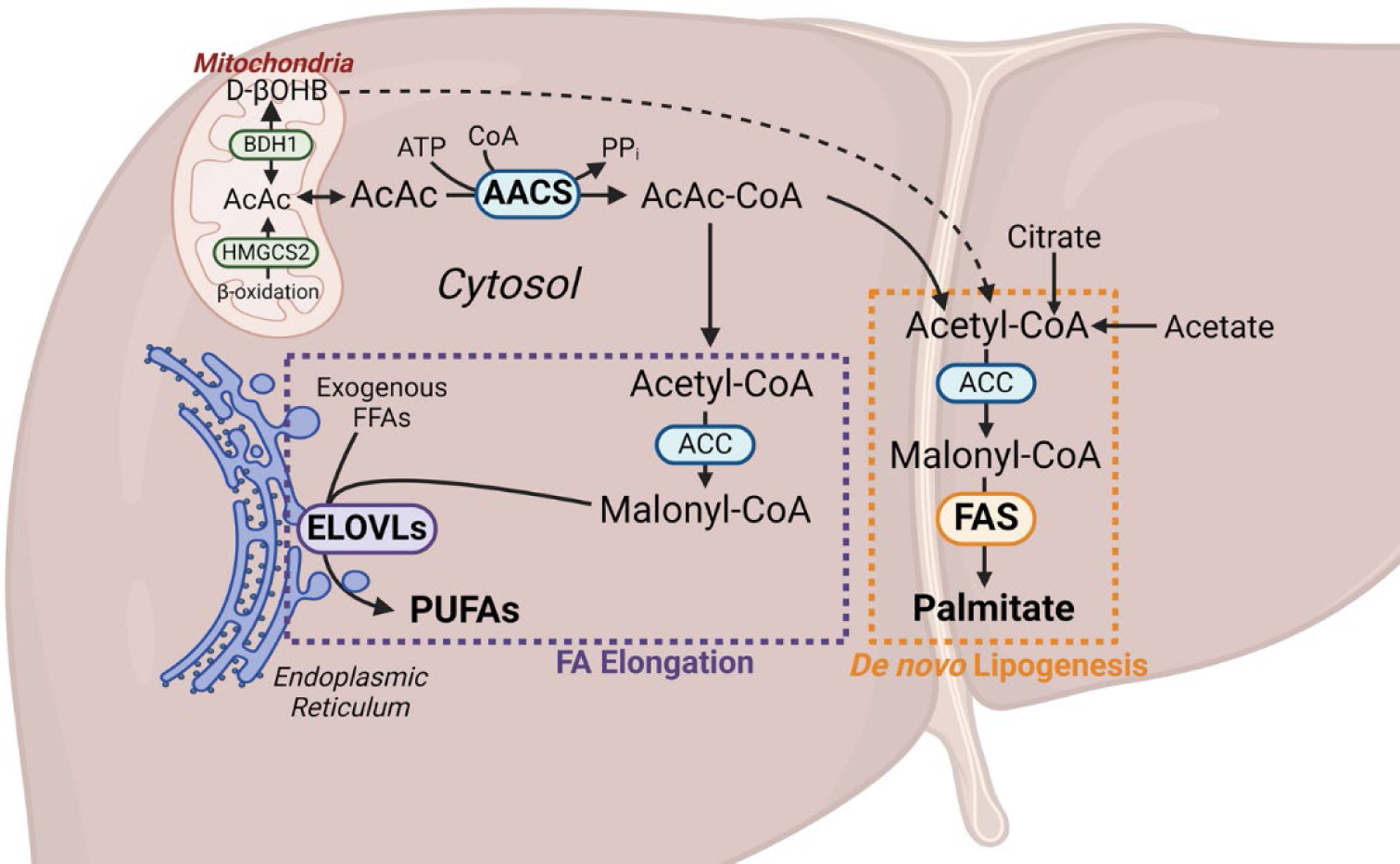
Model for ketogenesis-sourced DNL and FA elongation in hepatocytes. Ketogenesis synthesize the ketone bodies, AcAc and D-βOHB, which can both be used to support FA biosynthesis in hepatocytes, via contributing carbon to both DNL and the FA elongation pathways. The entry of ketone body-sourced carbon for DNL, appears to proceed through multiple redundant metabolic pathways, for which AACS only appears to be partially obligate. Sourcing of FA elongation from ketone bodies, however, appears to be fully dependent on AACS. Loss of AACS *in vitro*, or HMGCS2 *in vivo*, caused impairments in FA elongation, manifesting in a diminution of free and esterified PUFA pools, and dysregulation of the glycerophospholipidome, mirroring the lipidomics profile of human MASLD-MASH.

DNL occurs in the cytosol, catalyzed by FAS, a large multi-domain protein which polymerizes malonyl-CoA-derived acetyl units into palmitate^58,59^, leading to the successive incorporation of two-carbon units into a progressively lengthened, covalently attached acyl-chain. In contrast, FA elongation is not successive, occurs within the cytoplasmic leaflet of the endoplasmic reticulum (ER) membrane system, and begins with the condensation of a fatty acyl-CoA with malonyl-CoA via one of seven elongases (ELOVL1-7)^37,38^. Completion of a single elongation cycle requires engagement of multiple other separately encoded enzymes including (i) ketoacyl-CoA reductase, (ii) 3-hydroxyacyl-CoA dehydrogenase, and (iii) *trans*-2-enoyl-CoA reductase activities, which collectively lead to the incorporation of a single M+2 label into FAs, as opposed to the characteristic M+2, M+4, M+6… pattern which arises from DNL^37,38,60^. Independent desaturation cycles occur though the fatty acid desaturase (FADS) enzymes catalyzing delta-6 and -5 desaturation activities. The utilization of ketone bodies for both DNL and FA elongation pathways suggests a high-degree of subcellular compartmentalization, whereby ketogenesis in the mitochondria, DNL in the cytosol, and FA elongation in the ER membrane system, are coordinately regulated to support FA anabolic homeostasis in hepatocytes.

Analogous diminutions of PUFAs in hepatocytes derived from AACS KO mice and in livers derived from HMGCS2 KO supports a connection between endogenous ketogenesis and FA elongation. While changes in the PUFA-enriched PE pools were observed, this was not transmitted to the PUFA-enriched PC pool. One explanation for this PL class-specific effect could be that the relative provision of ketone-derived FAs into the Kennedy pathway, a major route for both PE and PC synthesis, is impaired, but is further coupled with a compensation in phosphatidylethanolamine N-methyltransferase (PEMT) activity, which converts PEs to PCs. Alternatively, dysregulation of the Land’s cycle, which increases de-acylation and conversion of PEs into LPEs could cause a PE-specific perturbation of PL pools.

Our data support a model in which hepatocytes sequester intra-compartmental acetyl-CoA pools within a hepatocyte, through colocalization of metabolic pathways/enzymes. It is also possible, and not mutually exclusive, that the roles of ketogenesis and ketone-derived FA elongation are distributed among cells in the zonal acinar axis. The liver’s microarchitectural periportal zone, with relatively higher oxygen tension, favors fatty acid oxidation, relative to the pericentral zone, which displays enhanced rates of lipogenesis^35,61^. HMGCS2 is uniformly expressed in hepatocytes across liver zones, but BDH1 expression is enriched in the pericentral zone, suggesting that ketogenesis could play a role in coordinating liver fat oxidation and synthesis across hepatic zones. Thus, intercellular exchange of ketone body-derived carbon among primary hepatocytes in culture cannot be excluded, and thus intracellular metabolic channeling of discrete metabolite pools versus distribution of these distinct metabolic functions among distinct hepatocyte populations will be of great interest in future experiments.

The lipidomics profiles of the ketogenesis insufficient (HMGCS2-deficient) mouse liver model, encompassed TAG steatosis, a depletion of PUFAs (free and esterified), and a diminution in the PC/PE ratio, which collectively mimicked the lipidomics signature observed in human MASLD^62–70^. These findings suggest that ketogenesis might be linked to the lipidomic remodeling of the liver that occurs in MASLD-MASH. Though numerous studies have linked ketogenesis to MASLD-MASH^19,71–74^, the underlying mechanisms and consequences remain unclear. Prior studies from our group and others demonstrated ketogenic insufficiency increases the susceptibility to MASLD-MASH by increasing hepatic oxidative fluxes and provoking a DNL metabolic and transcriptomic signature^24,75^. Given that PUFAs inhibit DNL by suppression of SREBP1c^76,77^, our findings are consistent with the notion that the diminution of PUFAs in HMGCS2 KO livers may attenuate the PUFA-mediated inhibition of SREBP1c, and thereby stimulate DNL. This novel mechanism is further supported by studies in human MASLD patients harboring the PNPLA3 I148M polymorphism^78,79^, whereby dysregulated PUFA partitioning into phospholipid pools disrupts hepatic TAG metabolism and VLDL secretion^78^. Moreover, the PNPLA3 I148M polymorphism is linked to increased ketogenesis, suggesting that ketogenesis may compensate for impaired PUFA metabolism.

Interestingly, PUFAs confer protection against numerous diseases including neurological^80–83^ and cardiovascular disease^84–86^, both of which are diseases for which ketone bodies have been heavily implicated^87–94^. Furthermore, hepatic PUFAs are inversely related to MASLD severity^62,63,65,69,70,95–97^. Our current findings therefore suggest that hepatic ketogenesis may influence hepatic and extrahepatic PUFA metabolism, representing a novel molecular mechanism through which ketone bodies could regulate systemic physiology and chronic human diseases, like MASLD-MASH.

Additionally, PUFAs possess anti-inflammatory properties, and are integral to immunological functions^98–100^. Ketone bodies also act as immunological regulators^27^. For example, D-βOHB inhibits the NLRP3 inflammasome, independent of its oxidation and known G-protein coupled receptors^101–103^. Thus, it is tempting to speculate that this known role of D-βOHB proceeds through a ketone-derived PUFA signaling intermediate. This hypothesis was however supported by the observation that [U-^13^C_4_]ketone bodies sourced arachidonic acid (20:4), thereby directly linking ketogenesis to the eicosanoid biosynthesis pathway^104,105^. Since ketone bodies are known to regulate inflammation and oxidative stress^26,27^, their ability to modulate immunological functions via reprogramming PUFA metabolism warrants further investigation.

## Limitations of Study

In this study, MS-based ^13^C-ITUM and multidimensional shotgun lipidomics enabled the detection and quantification of many diverse metabolites and lipid species in biological samples. The technologies used here, however, do not resolve positional isotopomers of chemical features, or determine the localization of double bonds within FAs or other PL species.

## Acknowledgements

The authors acknowledge support from the National Institutes of Health (grants DK091538, AG069781, DK007203, HL166142, and P30 AG013319). The authors are also grateful to Rumina Todorova and Justin Lengfeld for technical support.

## Author contributions

Conceptualization PP, EDQ, PAC

Methodology PP, EDQ, DBS, KF, XH, DAD, ABN, ZM, AH, JRG

Resources PAC, PP, HBR, XH

Writing-original draft EDQ, PP, PAC

Writing-review and editing All authors

Supervision PAC, PP, HBR

Funding acquisition PAC, HBR

## Materials and Methods

### Reagents and Chemicals

LCMS grade water (H_2_O) (Fisher, W6-4), LCMS grade methanol (MeOH) (Fisher, A456-4), LCMS grade acetonitrile (ACN) (Fisher, A955-4), LCMS grade isopropanol (Fisher, A461-4), LCMS grade chloroform (Fisher, C297-4), LCMS grade acetic acid (Sigma, 49199), Sodium D-[U-^13^C_4_]βOHB (CIL, CLM-3853-PK), naturally-occurring sodium D-βOHB (Sigma, 298360), naturally-occurring ethylacetoacetate (Sigma, W241512), [1,2,3,4-^13^C_4_]ethylacetoacetate (CIL, CLM-3297-PK), sodium [U-^13^C_2_]acetate (CIL, CLM-440-PK), naturally-occurring sodium acetate (Sigma, PHR2615), DMEM (high glucose) (Thermo, 11965092), DMEM (no glucose) (Thermo, A1443001), fetal bovine serum (Biotechne, S11150), Pen/Strep (Thermo, 15140122), L-glutamine (200 mM) (Thermo, 25030081), Phosphate Buffered Saline (PBS) (no CaCl_2_ or MgCl_2_) (Thermo, 14190144), Phosphate Buffered Saline (PBS) (Thermo, 14040133), Molecular Biology Grade Water (Corning, 46-000-CM), Collagen Type I (Corning, 354236), Collagenase from *Clostridium histolyticum* (Sigma, C5138), Surface treated polystyrene 6-well culture plates (Fisher, FB012927), 70µm sterile cell strainer (Corning, 431751), Trypan Blue Solution (Thermo, 15250061), Hanks’ Balanced Salt Solution (HBSS) (Ca^2+^, Mg^2+^, and phenol red-free) (Thermo, 14175095), Pierce BCA Protein Assay Kit (Thermo, 23225).

### Animals

All animal experiments were approved by the Institutional Animal Care and Use Committee at the University of Minnesota. All mice were bred on a C57BL/6NJ background, maintained on standard chow diet (2016 Teklad global 16% protein rodent diet), and received autoclaved water ad libitum. Mice were housed on corncob bedding in groups of 4 to 5 with lights off between 2000 and 0600 in a room maintained at 22°C.

Hepatocyte-specific β-hydroxybutyrate (βOHB) dehydrogenase 1 (BDH1) null mice were generated as previously described^39^, by crossing homozygous Bdh1^flox/flox^ mice^88^ to heterozygous Cre recombinase expressing mice, driven by the albumin promoter^106^. Littermate wild-type (WT) Cre-negative (Bdh1^flox/flox^, Cre^-/-^) animals were used as controls for Cre-positive (Bdh1^flox/flox^, Cre^+/-^) BDH1 knockout (KO) mice.

Whole-body acetoacetyl-CoA synthetase (AACS) null mice were generated by Jackson Laboratory (#042183) by injecting Cas9 nuclease and guide sequences designed to target and delete 256 bp in exon 2 of the Aacs gene, into C57BL/6NJ-derived fertilized eggs, followed by transference to pseudopregnant females, and subsequent breeding to C57BL/6NJ mice. Littermate homozygous WT (Aacs^+/+^) mice were used as controls for homozygous AACS KO (Aacs^-/-^) mice.

Hepatocyte-specific 3-hydroxymethylglutaryl-CoA synthase 2 (HMGCS2) null mice were generated by crossing homozygous Hmgcs2^flox/flox^ mice, with heterozygous Cre recombinase expressing mice, driven by the albumin promoter^106^. *Hmgcs2^flox/flox^* mice were generated in collaboration with the inGenious Targeting Laboratory (Ronkonkoma, NY). Briefly, BF1 clone C57BL/6N mouse embryonic stem (ES) cells harboring an expression construct for FLP recombinase were electroporated with a donor vector targeting exon 2 of the *Hmgcs2* locus (NM_008256), along with Cas9 and synthetic guide RNAs. Targeting exon 2 results in a frame shift mutation and resulting nonsense (premature stop) encoding *Hmgcs2* transcript. After selection for neomycin resistance using G418, and confirmation of successful targeting using both Southern blots and PCR, targeted ES cells were microinjected into Balb/c blastocysts, and chimeras with a high percentage black coat color were mated to C57BL/6N WT mice to generate Germline Neo Deleted mice (*Hmgcs2^flox/+^*). PCR and DNA sequencing were used to confirm (i) successful flanking of exon 2 with *loxP* sites, (ii) the loss of the neomycin resistance-encoding gene, and (iii) the absence of a FLP recombinase transgene in the progeny. These mice were crossed to B6.Cg-*Speer6-ps1^Tg^*(Alb–cre)*^21Mgn^*/J mice (the Jackson Laboratory). Littermate WT Alb-Cre-negative (*Hmgcs2^flox/flox^*, Alb-Cre^-/-^) animals were used as controls for Cre-positive (*Hmgcs2^flox/flox^*, Alb-Cre^+/-^) HMGCS2 KO mice, which are maintained on a C57BL6/NJ substrain hybrid background.

All mice were random-fed at the time of primary hepatocyte isolation or *ex vivo* liver perfusion. Both male and female mice were used for primary hepatocyte isolation experiments. For *in vivo* fasting experiments, food was removed at 1700 the evening prior to tissue collection and only male mice were studied. All mice were 10-20 weeks of age at time of experimentation.

### AcAc synthesis

Exogenous naturally-occurring [^12^C] and [U-^13^C_4_]acetoacetate was synthesized as previously described^107^. Briefly, naturally-occurring [^12^C]ethylacetoacetate or [1,2,3,4-^13^C_4_]ethylacetoacetate underwent a base-catalyzed hydrolysis using an equimolar amount of sodium hydroxide at 60°C, followed by neutralization with HCl, sterile filtration and storage at -80°C. Chemical purity was assessed via ^1^H-NMR and concentration was quantified via UHPLC-MS/MS^107^.

### Quantification of Total Ketone Bodies

Naturally-occurring [^12^C] and [^13^C]ketone bodies were formally quantified using UHPLC-MS/MS as described previously^39,107^. Briefly, for only naturally-occurring [^12^C]ketone measurements, [U-^13^C_4_]AcAc and [3,4,4,4- D_4_]βOHB internal standards were spiked into ice cold MeOH:ACN (1:1), then ketones were extracted, separated via reverse-phase UHPLC, and detected via parallel reaction monitoring (PRM) on a QExactive Plus hybrid quadrupole-orbitrap mass spectrometer. For naturally-occurring [^12^C] and exogenous [^13^C] ketone measurements, [3,4,4,4-D_4_]βOHB internal standard was spiked into samples, followed by reduction with NaBD_4_. ACN extracted ketone bodies were then bound to a cation-exchange resin, eluted into LCMS grade water, dried by SpeedVac, reconstituted into mobile phase (98% H_2_O, 2% MeOH, 0.0125% acetic acid), then separated and detected using the same UHPLC-MS/MS based approach as above, with modified PRM transitions.

### In vivo [^13^C] Stable-Isotope Delivery

Random-fed mice were injected with 10 µmol/gram body weight of sodium D-[U-^13^C_4_]βOHB or naturally-occurring D-[^12^C]βOHB intraperitoneally (IP), then animals were sacrificed via cervical dislocation. Serum was collected at various time points. Tissues were collected and rapidly freeze-clamped after 30 minutes post-IP injection and analyzed via ^13^C-ITUM.

### Ex vivo Liver Perfusion Protocol

Liver perfusions were performed as described previously^75^. Briefly, mice were anesthetized with IP sodium pentobarbital injection, externally sterilized with 70% ethanol, then mice were dissected to expose the hepatic portal vein, followed by cannulation with a 24-gauge catheter needle. The cannula was secured, then the abdominal aorta, inferior vena cava and cardiac right atrium were severed to isolate the liver from peripheral tissues and prevent recirculation. Livers were perfused with oxygenated Krebs-Henseleit buffer (118 mM NaCl, 4.7 mM KCl, 2.5 mM CaCl_2_, 1.2 mM KH_2_PO_4_, 1.22 mM MgSO_4_, 25 mM NaHCO_3_, pH 7.4), warmed to ∼45°C and supplemented with 1.5 mM sodium lactate, 0.15 mM sodium pyruvate, and 1 mM sodium D-[U-^13^C_4_]βOHB or sodium [U-^13^C_4_]AcAc. Buffer was oxygenated by continuous bubbling of a 95% O_2_, 5% CO_2_ gas mixture through the buffer reservoir using a fritted glass tube. Perfusion buffer was delivered using a peristaltic pump at a flow rate of ∼8 mL/min for 30 minutes. Buffer and effluent samples were collected every 10 minutes. At the completion of the perfusion, the liver was freeze-clamped in liquid nitrogen, and subsequently weighed to obtain total liver weight for flux normalizations. Perfusion buffer (input), effluent (output) and tissues were stored at -80°C until analysis. Extraction of exogenous [U-^13^C_4_]ketones was calculated by quantifying [^13^C]ketone bodies in the perfusion buffer and effluent via UHPLC-MS/MS, as described above, then performing a mass-balance analysis across the liver, as described below.

### Mass-balance Analysis in Ex Vivo Liver Perfusion

The net extraction of either [U-^13^C_4_]ketone by the perfused liver was calculated at steady-state by subtracting the moles of [U-^13^C_4_]ketones quantified in the effluent from the moles of [U-^13^C_4_]ketones in the perfusion buffer, at each time point (10, 20, 30 min). Average µmoles of [U-^13^C_4_]ketones extracted by the liver per minute was calculated and normalized to total wet liver weight. To assess the interconversion of exogenous [U-^13^C_4_]ketones via BDH1, a mass-balance for the net production of each [^13^C]redox partner was quantified. Average µmoles [U-^13^C_4_]ketone produced by the liver per minute was calculated, then normalized to total wet liver weight.

### In vitro Primary Mouse Hepatocyte Isolation Protocol

Primary mouse hepatocytes were isolated from ∼10-20 week old, random-fed mice, after a two-step collagenase liver perfusion. Mice were anaesthetized with IP injection of sodium pentobarbital, externally sterilized with 70% ethanol, then were dissected to expose the portal vein, which was cannulated with a 24-gauge catheter needle. The liver was perfused using a peristaltic pump at ∼4 mL/min for 8 minutes with Hank’s Balanced Salt Solution (without Mg^2+^, Ca^2+^, or phenol red) supplemented with 10 mM HEPES and 0.5 mM EGTA, at pH 7.4 and 45°C, subsequently followed by perfusion for 8 minutes with Hank’s Balanced Salt Solution (without Mg^2+^, Ca^2+^, or phenol red) supplemented with 10 mM HEPES, 5 mM CaCl_2_(2H_2_O) and 0.75 mg/mL collagenase type IV, at pH 7.4 and 45°C. Post-perfusion, the liver was dissected out of the body cavity, placed in a petri-dish with ∼10 mL of the warmed collagenase buffer, then was mechanically shredded using forceps and left to digest for another ∼5 minutes at room temperature. 10 mL of ice cold DMEM (high glucose) was added to the petri dish to quench the digestion, then cells were transferred to a 50 mL conical tube. The plate was rinsed with another 10 mL of cold DMEM, cells were pooled together, then were filtered through a sterile 70 μm plastic cell strainer into a new 50 mL conical tube. Cells were centrifuged at 50 x g for 1 min at 4°C to pellet hepatocytes. The supernatant was aspirated, then the hepatocyte pellet was washed by reconstitution into 25 mL of cold DMEM, followed by centrifugation at 50 x g for 1 min at 4°C. The pellet was washed another two times, for a total of three washes. Isolated primary hepatocytes were resuspended into 20 mL of warm (37°C) complete hepatocyte media, consisting of DMEM (high glucose) supplemented with 10% fetal bovine serum, 2 mM glutamine and 0.1% penicillin-streptomycin. Cells were counted and viability was determined using trypan blue method. Cells were diluted to ∼1 million cells per mL in complete media and seeded at a density of ∼1 million cells/well onto collagen type I coated 6-well plates. Hepatocytes were incubated at 37°C and 5% CO_2_ for ∼two hours to allow cells to attach, then adherent cells were washed with warm PBS, and cultured overnight in 1 mL of complete media. Isolated primary hepatocytes were used the next day for experiments.

### [U-^13^C_4_]ketone Stable-isotope Delivery in Primary Mouse Hepatocytes

Primary mouse hepatocytes were cultured as described above. The day after plating, primary hepatocytes were first serum starved for 1 hour in DMEM (no glucose), then cells were incubated in 1 mL of complete hepatocyte media, supplemented with 1 mM sodium [^12^C] or [U-^13^C_4_]ketones, for various times to allow ketone body-derived carbon to become incorporated throughout the metabolome of hepatocytes. After incubating with the ^12^C or ^13^C-labeled substrates, media were collected then cells were harvested for metabolomics and lipidomics analyses using previously described methods^25,34^ .

### Acetate-ketone Body Competition Experiment

Primary mouse hepatocytes were cultured as described above. The day after plating, hepatocytes were serum starved for 1 hour in DMEM (no glucose), then cells were incubated in 1 mL of complete hepatocyte media, supplemented with 1mM [U-^13^C_2_]acetate, in the absence (control), or presence of 1 mM naturally-occurring [^12^C] D-βOHB or AcAc. After 24 hours, cells were harvested, metabolites and lipids were extracted, then ^13^C-enrichment of FFAs was quantified as described for [^13^C]ketone tracing experiments.

### Metabolite Extraction from Cultured Cells and In Vivo Tissues

*For cells:* Conditioned media was collected and snap frozen, then adherent cells were washed twice with 1mL warm (37°C) PBS (-MgCl_2, -_CaCl_2_), twice with warm (37°C) cell-culture grade H_2_O, then the entire plate was submerged into liquid nitrogen to snap freeze cells, which rapidly quenches metabolism. To permanently preserve the metabolome, cells were harvested into 500 µL of cold (-20°C) LCMS grade MeOH per well of cells, then the MeOH was evaporated off using a SpeedVac. Dried cell pellets were stored at -80°C until analysis. Metabolites/lipids were extracted as described below. *For tissues:* tissues were rapidly freeze-clamped and stored at -80°C. Tissues were lyophilized and metabolites/lipids extracted as follows. *For both cells and tissues:* to lyophilized tissue (∼2 mg) or dried cell pellet (∼2 million cells), metabolites were extracted into 1 mL of ice cold (-20°C) LCMS grade MeOH:ACN:H_2_O (2:2:1) via three rounds of vortexing, freeze-thawing and water bath sonication. Samples were incubated at -20°C for ∼1 hour, centrifuged for 10 minutes at 15,000 x g, then an equal volume of supernatant was transferred to fresh tubes, and the solvent was evaporated completely in a SpeedVac at room temperature. The dried metabolite pellet was reconstituted into 40 µL (cells) or 100 µL (tissue) of LCMS grade ACN:H_2_O (1:1) using vortexing and sonication, followed by incubation at 4°C for ∼1hr, and centrifugation at 15,000 x g for 10 minutes. The supernatant was transferred to a glass LC-MS vial, and analyzed via ^13^C-ITUM as described below.

### ^13^C Stable Isotope <u>T</u>racing <u>U</u>ntargeted <u>M</u>etabolomics (ITUM) pipeline^25,34^

#### Sample Preparation and Raw Data Acquisition

Metabolites were first extracted into MeOH:ACN:H_2_O (2:2:1), reconstituted into ACN:H_2_O (1:1), then were separated using liquid chromatography (LC) on a Thermo Vanquish UHPLC system, followed by detection using full-scan high-resolution mass spectrometry (HR-MS) on a Thermo QExactive plus hybrid quadrupole-orbitrap mass-spectrometer, fitted with a heated electrospray ionization source operated in either negative or positive polarity mode. To expand the chemical landscape surveyed, both hydrophilic interaction chromatography (HILIC) and reverse phase (RP) liquid chromatography were employed, using a total of three unique UHPLC methods: **[1]** SeQuant ZIC-pHILIC column (2.1 x 150 mm, 5 µm) (Millipore Sigma, 1.50460). Mobile phase A (MPA) was 95% H_2_O, 5% ACN, 10 mM ammonium acetate, and 10 mM ammonium hydroxide. Mobile phase B (MPB) was 100% ACN. The total run time was 50 minutes, flow rate was 2 mL/min, and column chamber was set to 45°C. Mobile phase gradient was as follows: 0-0.5 min, 90% MPB; 0.5-30 min, 90→30% MPB; 30-31 min, 30% MPB; 31-32 min, 30→0% MPB; 32-33 min, 0→90% MPB; 33-50 min, 90% MPB. **[2]** Acquity UPLC BEH Amide (2.1 x 150 mm, 1.7 µm) (Waters, 186004802). MPA was 50% H_2_O, 50% ACN, 10 mM ammonium acetate. MPB was 5% H_2_O, 95% ACN, 10 mM ammonium acetate. The total run time was 10 minutes, flow rate was 0.6 mL/min, and column chamber was 45°C. Mobile phase gradient was as follows: 0-2 min, 99→80% MPB; 2-5 min, 80→20% MPB; 5-8 min, 20→99% MPB; 8-10 min, 99% MPB. **[3]** Hypersil GOLD aQ C18 column (1 x 150 mm, 3 µm) (Thermo, 25303-151030). MPA was 60% ACN, 40% H_2_O, 0.1% formic acid, and 10 mM ammonium formate. MPB was 10% ACN, 90% IPA, 0.1% formic acid, and 10 mM ammonium formate. The total run time was 30 minutes, flow rate was 0.2 mL/min, and column chamber was 45°C. Mobile phase gradient was as follows: 0-3 min, 5% MPB; 3-5 min, 5→15% MPB; 5-10 min, 15→48% MPB; 10-15 min, 48→82% MPB; 15-18 min, 82→95% MPB; 18-23 min, 95→15% MPB; 23-30 min, 15% MPB.

For all ^13^C-ITUM pipelines, both blanks and pooled quality control (QC) samples are injected periodically throughout the run. Blanks were ACN:H_2_O (1:1) and the QC sample was a pooled sample including all naturally-occurring [^12^C] and [^13^C]-labeled samples. To aid in chemical feature identification, two additional samples were also injected. First, was a standard mix consisting of authentic standards, for all expected analytes. Second, was a pooled sample consisting only of the naturally-occurring samples, analyzed via data-dependent analysis (DDA) tandem mass spectrometry (MS/MS) using IE omics script^108^ and R-Studio. Data processing and initial analysis was performed using Thermo Compound Discoverer 3.3. After raw mass spectra were uploaded, background ions were removed, retention time (RT) for detected signals were aligned across samples, chemical formulas were predicted, then grouped chemical features were profiled to determine compound identity, based on (1) the m/z predicted from the chemical formula, (2) the RT compared to an authentic external standard, and (3) the MS/MS fragmentation pattern, compared to in-house standards or online databases. Putatively identified metabolites and lipids were then carried forward and [^13^C] stable-isotope enrichment of each metabolic pool was determined after correction for natural abundance as described below.

### ^13^C-stable isotope tracing analysis

To carry out [^13^C] stable isotope tracing, first all [^13^C] mass isotopologues within the isotopic envelope of each identified metabolic pool were identified based on the diagnostic shift in m/z (Δm/z = 1.0033 Da, natural abundance, 1.11% of all carbon) induced by the presence of [^13^C] in chemical compounds. Raw ion counts for each isotopologue were extracted, summed, and expressed as a percentage of the total pool. After natural abundance correction, the fractional intensities were then graphed as a function of [^13^C] content, generating mass isotopologue distributions (MIDs) for each detected metabolite or lipid. [^13^C] enrichment of analytes was then reported as either fractional [^13^C] enrichment or [^13^C] atom percent enrichment (APE). *Fractional [^13^C] enrichment*, which reports on the percentage of molecules in a given metabolic pool that are enriched in [^13^C], was calculated by summing together the relative intensities (% of the total pool) for all [^13^C] mass isotopologues detected. *[^13^C] APE,* which reports on the percentage of the carbon atoms within a given metabolic pool that are enriched in [^13^C], was calculated by first weighting isotopologues based on [^13^C] carbon content, then summing together the weighted relative intensities for all [^13^C] mass isotopologues, and is given by the equation: 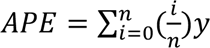 where *i* is the number of ^13^C carbons in an isotopologue, n is the total number carbons possible in the pool, and y is the fractional ^13^C enrichment of each isotopologue obtained from the ^13^C-MIDs. When relevant, the ^13^C-MID of specific metabolites/lipids are also presented, and inferences about metabolic flux are made based on the ^13^C-enrichment patterns.

### Tracer-dilution modeling of endogenous ketogenesis

Primary hepatocytes were incubated with either 2 mM D-[U-^13^C_4_]βOHB or 1 mM [U-^13^C_4_]AcAc for 24 hours, then both naturally-occurring endogenous [^12^C] ketone bodies and [^13^C] exogenous ketone bodies were quantified in conditioned media. The magnitude of tracer dilution was used as a surrogate of formal flux and was calculated using the following equation: Ra_ketogenesis_ 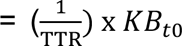, in which TTR is the tracer:tracee ratio, calculated as total ^13^C ketone &&’ bodies divided by total naturally-occurring, unlabeled [^12^C]ketone bodies, in conditioned media after 24 hours, and KB_t0_ is the initial concentration (mM) of [U-^13^C_4_]ketone body.

### Quantification of Nucleotides

Energy nucleotides (ATP, ADP and AMP), and redox nucleotides (NAD^+^ and NADH) were measured as previously described^39,109^, with modifications. Briefly, metabolites were extracted from cells for ITUM in MeOH:ACN:H_2_O (2:2:1), then extracts were separated and detected using ion-pairing reverse-phase UHPLC-MS/MS on a C_18_ column (Waters Xbridge, 150 x 2.1mm, 3µm). Nucleotides were detected as adducts of dibutylamine acetate on a Thermo QExactive Plus mass spectrometer, operated in positive ionization mode, using parallel reaction monitoring transitions as previously described^39,109^.

### Isotopic Enrichment of Acetyl-CoA and Malonyl-CoA

Acetyl-CoA and malonyl-CoA were extracted from cells for ITUM in MeOH:ACN:H_2_O (2:2:1), then were separated and detected using the same UHPLC-MS/MS method as for the quantification of nucleotides, adapted from previous publications^39,109^. The transitions were modified in order to detect the [^13^C] mass isotopologues for both acetyl-CoA and malonyl-CoA.

### Multidimensional Mass Spectrometry-based Shotgun Lipidomics

*For cells:* conditioned media was aspirated, then adherent cultured cells were washed twice with 0.1x PBS (-MgCl_2,_ -CaCl_2_), followed by trypsinization in 300 µL of 0.025% trypsin 0.01% EDTA PBS solution. Cells were collected into eppendorf tubes, then each well was washed with cold DMEM and cells were pooled together, followed by centrifugation to pellet cells. Cells were washed by reconstitution into 1 mL of 0.1x PBS (-MgCl_2,_ -CaCl_2_), followed by centrifugation. Cells were washed a total of two times, then cell pellets were snap frozen in liquid nitrogen and stored at -80°C. ∼1 million cells were homogenized in 300 µL ice-cold 0.1 x PBS (-MgCl_2,_ -CaCl_2_) using multiple cycles of freeze-thawing, vortexing and water bath sonication. *For tissues*: ∼20 mg freeze clamped liver was homogenized in 1 mL of ice-cold 0.1x PBS (-MgCl_2,_ -CaCl_2_) using zirconium oxide beads and tissue homogenizer (Omni International, Bead Ruptor 12, 19-050A). *For both cells and tissues:* total protein was first quantified in sample homogenates using bicinchoninic acid (BCA) protein assay, then an appropriate amount of internal standard mix (2.909 nmol [D_4_]16:0 FFA; 14.83 nmol D16:1 PE; 14.772 nmol D14:1 PC; 15.013 nmol T17:1 TAG) were added per mg of protein, followed by bulk lipid extraction using a modified Bligh-Dyer method as described previously^75,110,111^. Samples were directly infused into a TSQ Vantage triple quadrupole mass spectrometer (Thermo Fisher Scientific TSQ Vantage, San Jose, CA) equipped with an automated nanospray device (Triversa Nanomate, Advion Biosciences, Ithaca, NY) and multidimensional mass spectrometry-based shotgun lipidomics was performed^112^. FFA^113^ and PE^114^ lipids were derivatized with 4-(aminomethyl)-1-methylpyridin-1-ium (AMPP) or fluorenylmethoxylcarbonyl (FMOC), respectively. Identification and quantification of lipids was performed using in-house software.

### Immunoblotting

Cultured primary mouse hepatocytes were washed with warm (37°C) 1x PBS then trypsinized into 300 µL of a 0.025% trypsin 0.01% EDTA PBS solution. Cells were collected into eppendorf tubes, then wells were washed with 300 µL of DMEM to quench trypsin, followed by centrifugation to pellet cells. Cells were washed by suspending in 1 mL of 1x PBS, followed by centrifugation. The supernatant was discarded and then cell pellets were frozen in liquid nitrogen and stored at -80°C. For protein analysis, cells were thawed on ice then lysed in ∼300 µL RIPA buffer supplemented with protease and phosphatase inhibitor cocktails. Total protein was quantified by BCA assay, then 10 µg protein was separated on a 4-12% Bis-Tris polyacrylamide gel using the ThermoFisher NuPAGE system. Proteins were transferred to PVDF membranes (0.45µm, Thermo, 88518), equal loading was assessed using BlotFastStain (G-Biosciences, 786-34), then membranes were blocked in 5% NFDM in TBST overnight at 4°C. Primary antibodies against BDH1 (Atlas Antibodies, HPA030947), AACS (Proteintech, 13815-1-AP), HMGCS2 (Santa Cruz, Sc-33828) or SCOT (Proteintech, 12175-1-AP) were diluted ∼1:2000-5000 in TBST with 5% NFDM and were incubated overnight. Following washing in TBST, membranes were incubated with horseradish peroxidase (HRP)-conjugated anti-rabbit IgG secondary antibody in TBST with 5% NFDM for 1-2 hours. Blots were developed using SuperSignal West Pico PLUS Chemiluminescent substrate (Thermo, 34577) and imaged on a Bio-Rad ChemiDoc MP imaging system.

### Immunohistochemistry

For immunohistochemical detection of HMGCS2 protein in liver tissue, goat antihuman/mouse HMGCS2 antibody (Santa Cruz, Sc-33828), was used in combination with Alexa Fluor 488-conjugated donkey anti-goat IgG (Invitrogen). Antibodies were diluted 1:100, incubated, then washed with phosphate-buffered saline, containing 1% bovine serum albumin (BSA) and 0.1% Triton X-100 before imaging.

### Gene Expression

For mRNA expression analysis, cells or tissue were harvested and homogenized in Qiazol, then total RNA was isolated using Qiagen’s RNeasy protocol. First-strand cDNA was synthesized using iScript (BioRad, 170-8891), then quantitative real-time PCR was performed on a BioRad CFX384 Real-Time thermocycler using SsoAdvanced Universal SYBR Green supermix (BioRad, 170-5274). mRNA expression was calculated using the 2^-ΔΔCt^ method, where ΔCt = housekeeping gene Ct – gene of interest CT, then expressed as fold-change relative to the intact liver. mRNA levels were normalized to the intact liver, which was set to 1. Housekeeping gene was L32 for all targets. Primer sequences are given in **Supplementary Table 10.**

### Statistics

All analyses were performed using GraphPad Prism version 9. Unpaired 2-tailed Student’s *t*-tests or 1-way ANOVA, was used to determine statistically significant differences. P-value < 0.05 was accepted as significant for all tests. Multiple comparison corrections using the Holm-Sidak method were applied as appropriate. When appropriate, 2-way ANOVA was used to determine significant main effects and interactions amount variables, with Tukey’s post-hoc t-test applied as appropriate. Information about statistical analysis is included in each figure legend. Data are presented as mean ± SD, unless otherwise indicated.

## Acronyms

MALSD: metabolic dysfunction-associated steatotic liver disease
MASH: metabolic dysfunction-associated steatohepatitis
AcAc: acetoacetate
D-βOHB: D-β-hydroxybutyrate
FA: fatty acid
DNL: *de novo* lipogenesis
PUFAs: polyunsaturated fatty acids<colcnt=2>
BDH1: D-βOHB dehydrogenase 1
AACS: acetoacetyl-CoA synthetase
PE: phosphatidylethanolamine
PC: phosphatidylcholine
HMGCS2: 3-hydroxymethylglutaryl-CoA synthase 2
TAG: triacylglycerol
HMGCL: HMG-CoA lyase
SCOT: Succinyl-CoA:3-ketoacid-CoA Transferase
ITUM: [^13^C] stable-isotope tracing untargeted metabolomics
IP: intraperitoneally
UHPLC: ultra-high performance liquid chromatography
MS/MS: tandem mass spectrometry
MID: mass isotopologue distribution
PL: phospholipid
PG: phosphatidylglycerol
PI: phosphatidylinositol
PS: phosphatidylserine
FFA: free fatty acid
SFA: saturated fatty acid
MUFA: monounsaturated fatty acid
FAS: fatty acid synthase
ER: endoplasmic reticulum
ELOVL: elongases (elongation-of-very-long-chain-fatty acids)

