## Supplemental figures and tables for "Ketogenesis supports hepatic polyunsaturated fatty acid homeostasis via fatty acid elongation"

**Supplemental Figure 1**

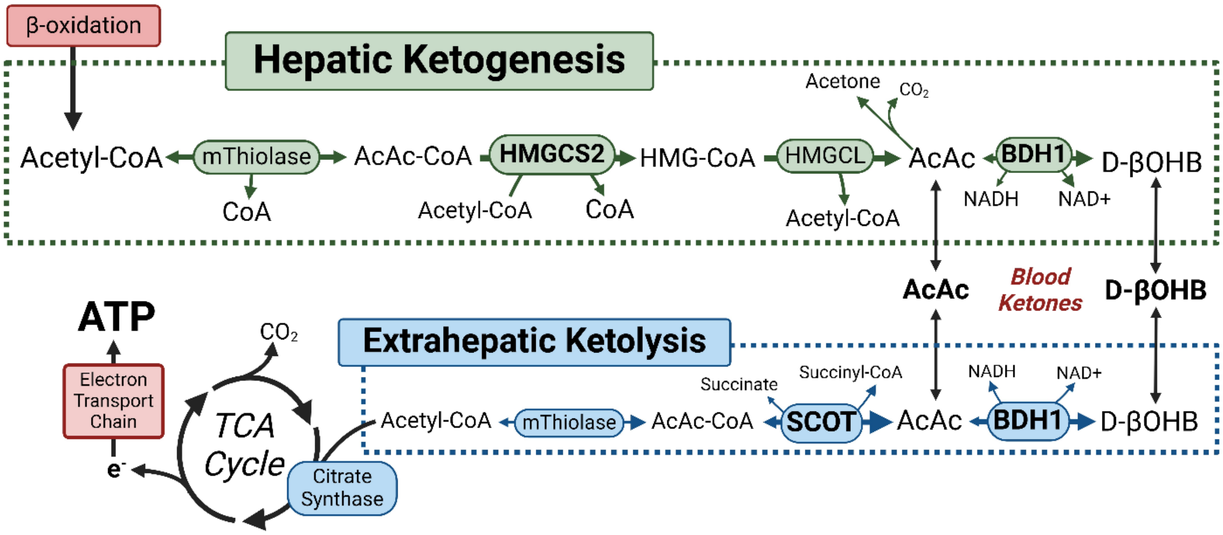

**Supplementary Figure 1: Overview of Canonical Ketone Body Metabolism**

Robust ketogenesis occurs almost exclusively in the hepatocyte mitochondria, and synthesizes acetoacetate (AcAc) and D-beta-hydroxybutyrate (D-βOHB) from excess β-oxidation derived acetyl-CoA. Other tissues (e.g., kidney, gut) can perform ketogenesis, however, the contribution of extrahepatic ketogenesis to circulating ketone bodies is neglible. Fasting decreases circulating insulin, triggering the mobilization of adipose-derived free fatty acids (FFAs), which stimulates hepatic fat oxidation and ketogenesis. Ketogenesis begins with the condensation of two molecules of acetyl-CoA via mitochondrial thiolase, generating acetoacetyl-CoA (AcAc-CoA), which then condenses with another molecule of acetyl-CoA to form 3-hydroxymethylglutaryl-CoA (HMG-CoA) via the rate limiting enzyme of the ketogenesis pathway, HMG-CoA synthase 2 (HMGCS2). AcAc is synthesized by cleavage from HMG-CoA by HMG-CoA Lyase (HMGCL). AcAc then forms a redox partner with D-βOHB, via the actions of mitochondrial NAD-dependent D-βOHB Dehydrogenase 1 (BDH1). The spontaneous decarboxylation of AcAc also results in the minor non-enzymatic production of acetone. **(Bottom)** AcAc and D-βOHB passively diffuse down concentration gradients, into circulation, into extrahepatic tissues. To utilize D-βOHB, tissues must express the enzyme BDH1, which facilitates the oxidation of D-βOHB back into AcAc. The mitochondrial rate limiting enzyme of the ketolysis pathway, Succinyl-CoA:3-ketoacid-CoA Transferase (SCOT), activates AcAc through transesterification with succinyl-CoA generating AcAc-CoA, which is in equilibrium with acetyl-CoA via mitochondrial thiolase. Acetyl-CoA, through the irreversible action of citrate synthase, enters the TCA cycle for terminal oxidation, which stimulates flux through the electron transport chain and oxidative phosphorylation, thereby providing fuel for extrahepatic tissues. Ketone body derived carbon is also sourced for anabolic reactions (not shown here).

**Supplemental Figure 2**

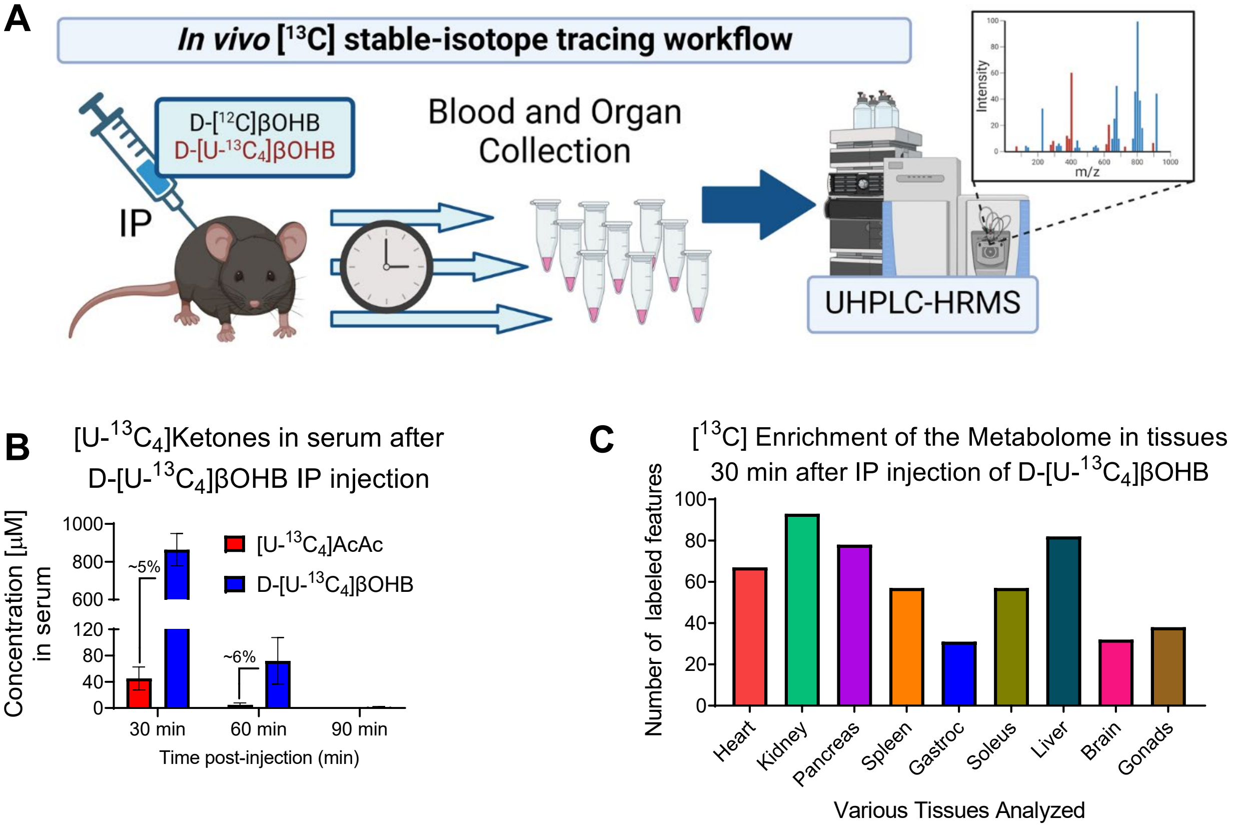

**Supplementary Figure 2. D-[U-$^{13}C_4$]βOHB rapidly integrates into the metabolome of numerous tissues *in vivo*, including the liver.**

**(A)** Random-fed mice, maintained on a standard chow diet, were injected intraperitoneally (IP) with naturally-occurring (‘[$^{12}C$]’)sodium D-βOHB or D-[U-$^{13}C_4$]βOHB (10 µmol/gram body weight; n=4 mice). Blood was collected at various times. Tissues were collected and freeze-clamped at 30 minutes post-injection. All tissues were lyophilized and analyzed using $^{13}C$-stable isotope tracing untargeted metabolomics (ITUM). **(B)** The concentration (µM) of [U-$^{13}C_4$]ketones in serum at 30, 60 and 90 minutes after injection of D-[U-$^{13}C_4$]βOHB measured via UHPLC-MS/MS (n=2/group). Data shown as average ± SD. Percentages show degree of systemic interconversion of injected D-[U-$^{13}C_4$]βOHB into [U-$^{13}C_4$]AcAc. **(C)** The total number of $^{13}C$-enriched chemical features (*i.e.,* unique retention time and m/z pair) in each tissue analyzed at 30 minutes post-injection using a pooled sample from each tissue. For confident feature identification, signals had to be detected across all biological replicates, and therefore the total number of $^{13}C$-enriched features is the same across all tissue replicates, and displays no variance. Supplementary Table 1 provides the retention time and molecular weight for all $^{13}C$-enriched chemical features in each tissue (n=4/group).

### Supplemental Figure 3

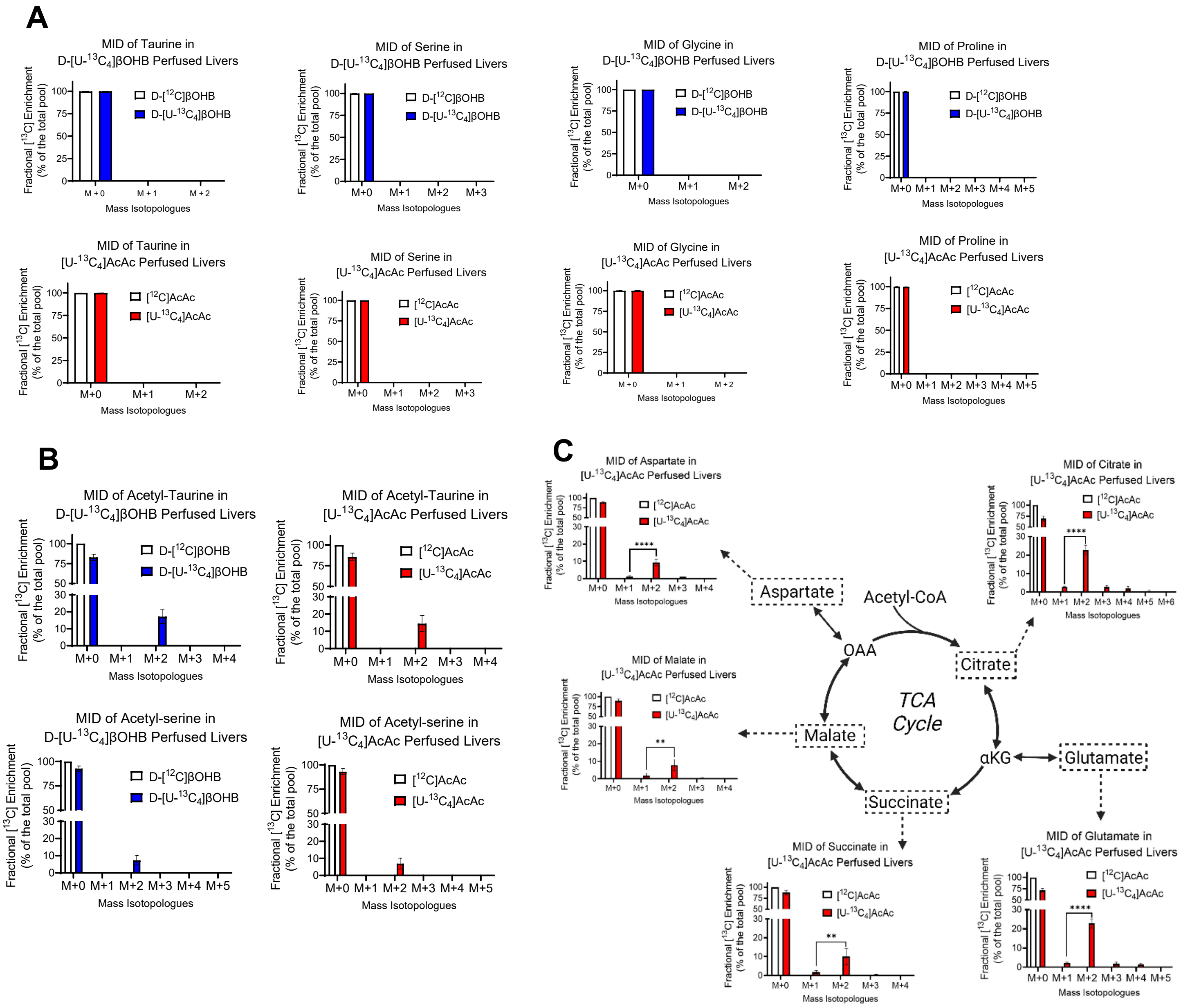

**Supplementary Figure 3. [U-$^{13}C_4$]ketone bodies integrate into distinct metabolic pools in the perfused liver.**

MID of **(A)** amino acids, including taurine, serine, glycine and proline, **(B)** acetyl-taurine and acetyl-serine and **(C)** TCA cycle intermediates from 1 mM sodium D-[U-$^{13}C_4$]βOHB (blue) or 1 mM sodium [U-$^{13}C_4$]AcAc (red) in freeze-clamped and lyophilized livers post 30 minute perfusion analyzed via [$^{13}C$] ITUM (n=4-5/group).

Data are expressed a mean ± SD. Statistically significant differences were determined by Student's t test, and accepted as $p<0.05$. *$p < 0.05$, **$p < 0.01$, ***$p < 0.001$, ****$p < 0.0001$ as indicated. ns = not significant.

### Supplemental Figure 4

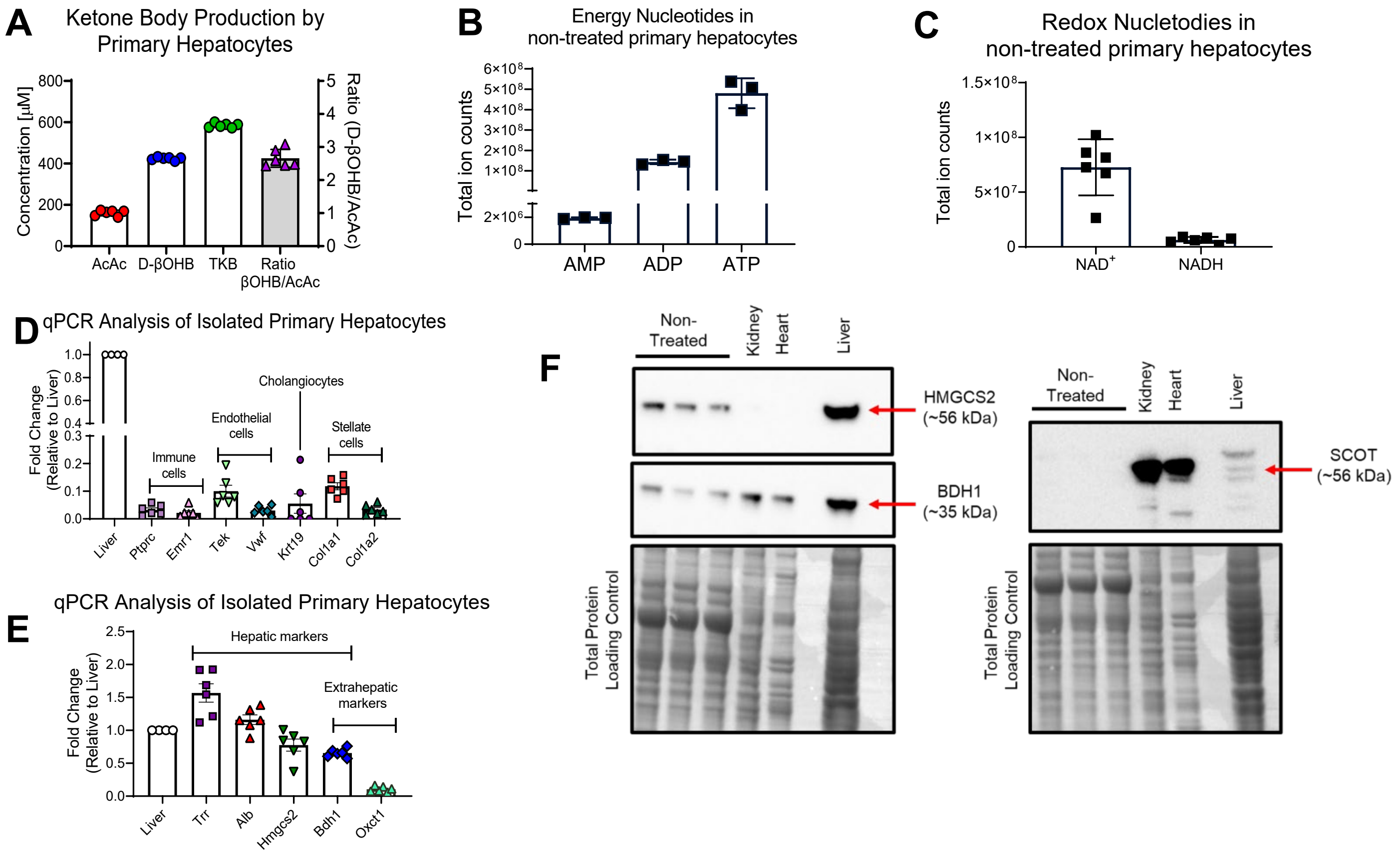

**Supplementary Figure 4. Validation of Primary Mouse Hepatocyte Model**

**(A)** Concentration (μM) of ketone bodies in conditioned media from cultured primary hepatocytes. **(B)** Total ion count of ATP, ADP, AMP, and **(C)** $NAD^+$ and NADH in primary hepatocytes (n=3-6/group). **(D)** Transcript abundances of non-hepatocyte cell type markers (immune cells (*Ptptc, Emr1*), endothelial cells (*Tek, Vwf*), cholangiocytes (*Kirt19)* and stellate cells (*Col1a1, Col1a2*)), and **(E)** hepatic (*Trr, Alb, Hmgcs2, Bdh1*) and extrahepatic markers (*Bdh1, Oxct1*) measured via qRT-PCR in isolated primary hepatocytes and whole liver lysate (n=6/group). House-keeping gene was L32 in all. **(F)** Immunoblot for HMGCS2, BDH1 and SCOT proteins migrating at ~56 kDa, ~35 kDa and ~56 kDa, respectively, in isolated non-treated primary hepatocytes and from whole tissue kidney, heart and liver lysates. Kidney and heart protein lysates were used as positive controls for BDH1 and SCOT, and negative controls for HMGCS2. Liver lysates were used as positive control for HMGCS2 and BDH1, and negative control for SCOT. Total protein assessed via FastStain.

Data are expressed a mean ± SD. Total protein for western blots was quantified via FastStain and shown above blots.

### Supplemental Figure 5

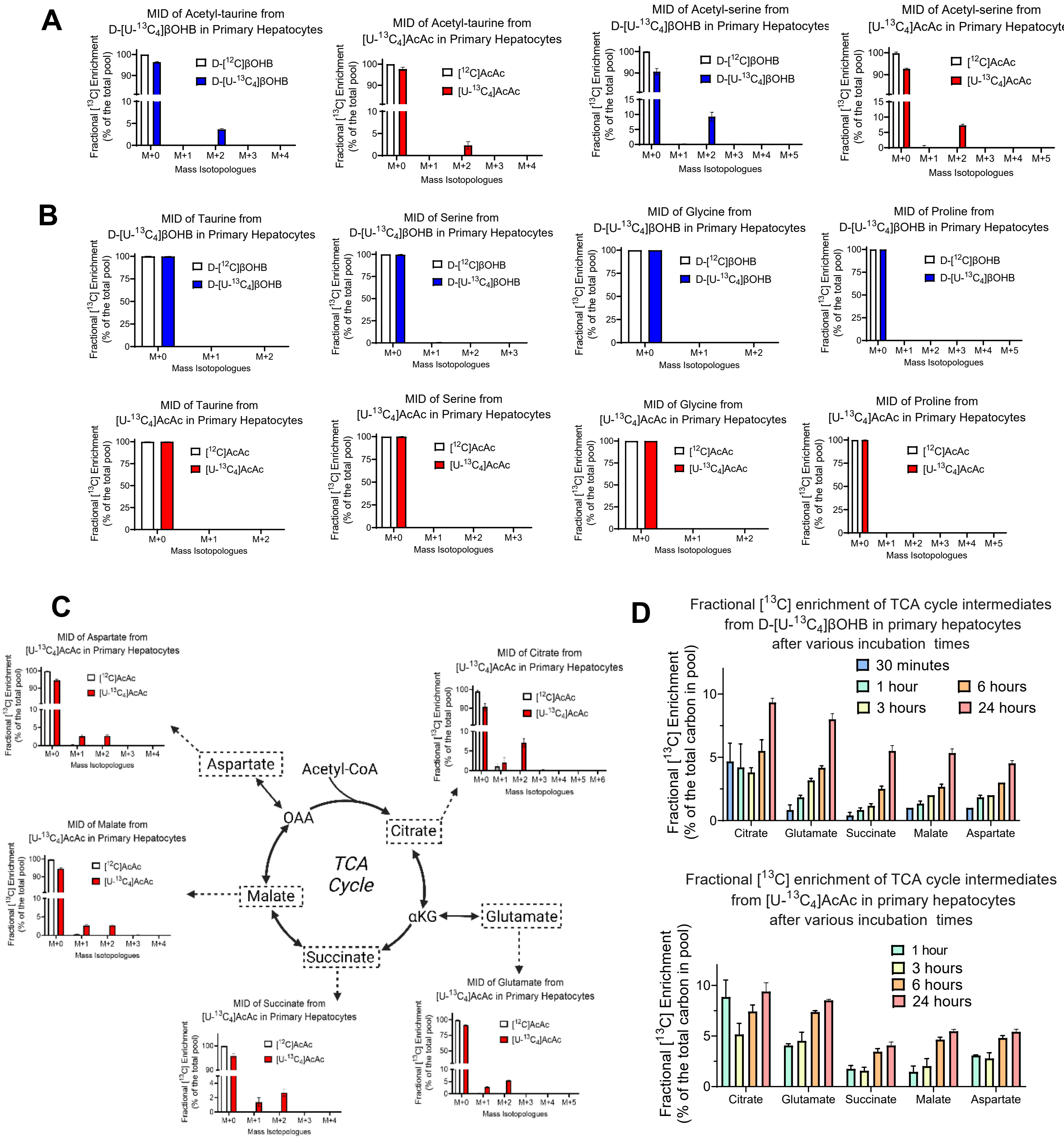

**Supplementary Figure 5. [U-$^{13}C_4$]ketone bodies integrate into distinct metabolic pools in primary hepatocytes.**

MID of **(A)** acetyl-taurine and acetyl-serine, **(B)** amino acids, including taurine, serine, glycine and proline, and **(C)** TCA cycle intermediates from 1 mM sodium D-[U-$^{13}C_4$]βOHB (blue) or 1 mM sodium [U-$^{13}C_4$]AcAc (red) in primary hepatocytes after 24 hours analyzed via [$^{13}C$] ITUM (n=6/group). **(D)** Fractional [$^{13}C$] enrichment of TCA cycle intermediates from 1 mM sodium D-[U-$^{13}C_4$]βOHB (top) or 1 mM sodium [U-$^{13}C_4$]AcAc (bottom) in primary hepatocytes after various incubation lengths (n=6/group).

Data are expressed a mean ± SD. Statistical differences were determined by Student's t-test, and accepted as significant if $p<0.05$. *$p < 0.05$, **$p < 0.01$, ***$p < 0.001$, ****$p < 0.0001$, as indicated.

### Supplemental Figure 6

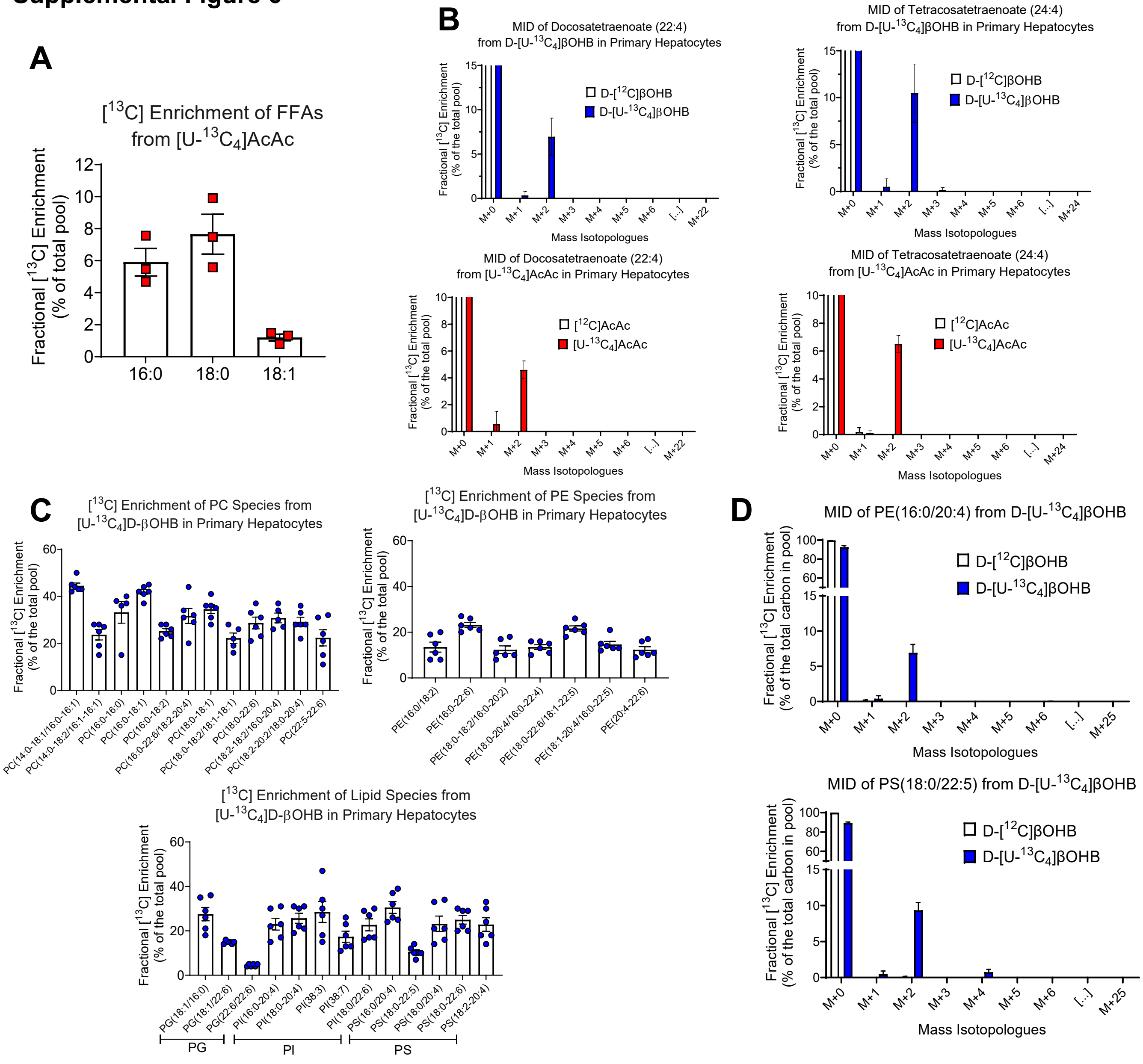

**Supplementary Figure 6. Both AcAc and D-βOHB are incorporated extensively throughout the lipidome.**

**(A)** Fractional [$^{13}C$] enrichment of the FFAs palmitate (16:0), stearate (18:0) and oleate (18:1) from 1mM sodium [U-$^{13}C_4$] AcAc in primary hepatocytes after 24 hours (n=3/group). **(B)** MIDs of long chain PUFAs from 2 mM sodium D-[U-$^{13}C_4$]βOHB (blue) or 1 mM sodium [U-$^{13}C_4$]AcAc (red) in primary hepatocytes after 24 hours (n=3/group). **(C)** Fractional [$^{13}C$] enrichment of individual phosphatidylcholine (PC), phosphatidylethanolamine (PE), phosphatidylglycerol (PG), phosphatidylinositol (PI) and phosphatidylserine (PS) lipid species from 1mM D-[U-$^{13}C_4$]βOHB in primary hepatocytes (n=6/group). **(D)** MID of PE(16:0/20:4) and PS(18:0/22:5) from 1mM D-[U-$^{13}C_4$]βOHB in primary hepatocytes (n=3/group).

Data are expressed a mean ± SD.

**Supplemental Figure 7**

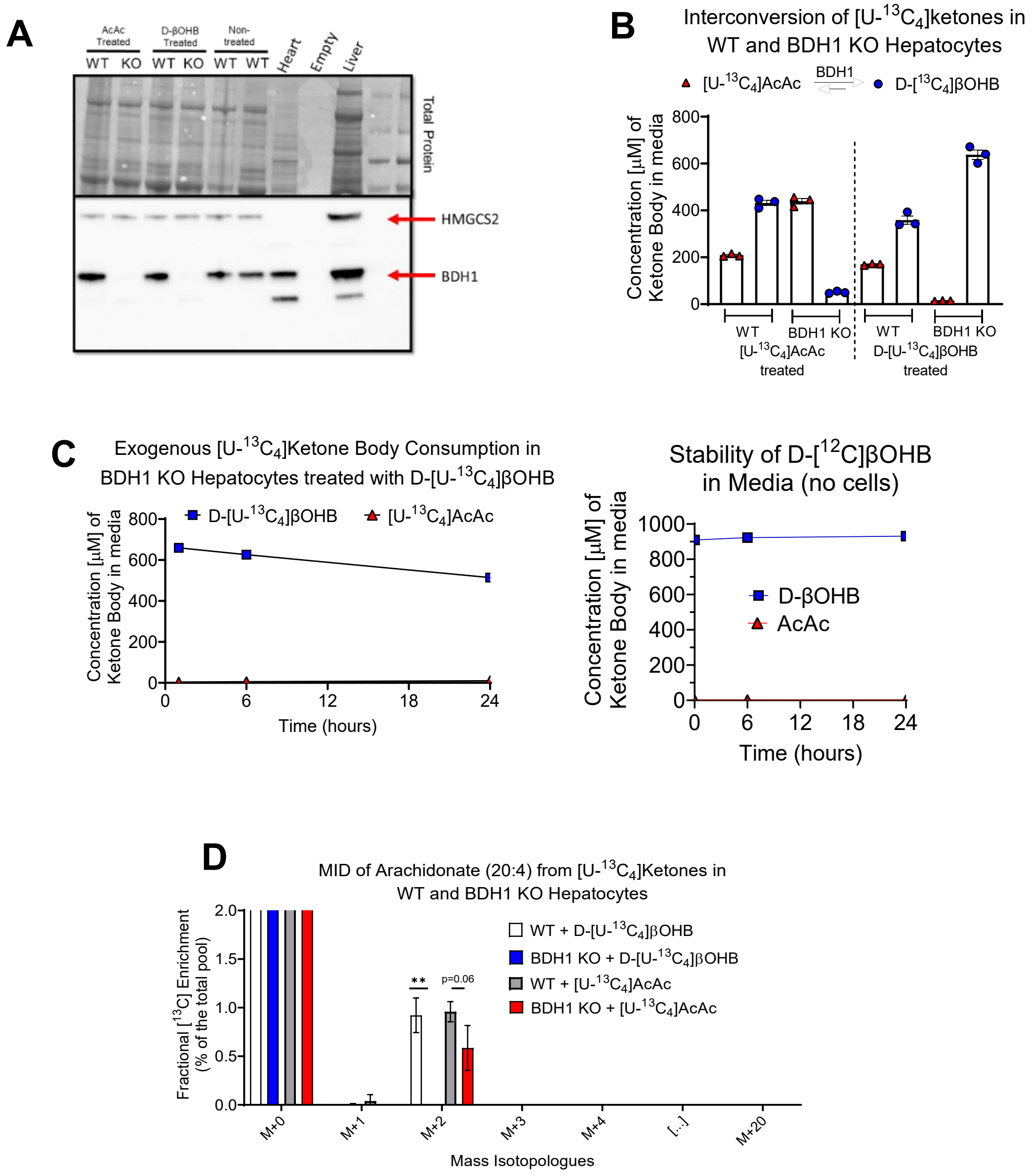

**Supplementary Figure 7. Validation of BDH1 KO primary hepatocytes model**

**(A)** Immunoblot for BDH1 protein migrating at ~35 kDa in WT and BDH1 KO primary hepatocytes. Heart and liver used as positive control for BDH1. Total protein assessed via FastStain. **(B)** Concentration (μM) of [U-$^{13}C_4$]ketones in conditioned media after 24 hour treatment or **(C)** in the media over 24 hours in WT and BDH1 KO hepatocytes treated with 1mM [U-$^{13}C_4$]AcAc or D-[U-$^{13}C_4$]βOHB (n=3/group). BDH1 KO hepatocytes still consume ketones, via BDH1-independent pathways, despite D-βOHB is chemically stable (right) (n=3/group). **(D)** MID of PUFA species from 1 mM sodium [U-$^{13}C_4$]ketones in WT and BDH1 KO hepatocytes (n=3/group) after 24 hours.

Data are expressed a mean ± SD. Statistical differences between genotypes were determined by Student's t test, and accepted as significant if $p<0.05$. *$p < 0.05$; **$p < 0.01$; ***$p < 0.001$; ****$p < 0.0001$; as indicated.

### Supplemental Figure 8

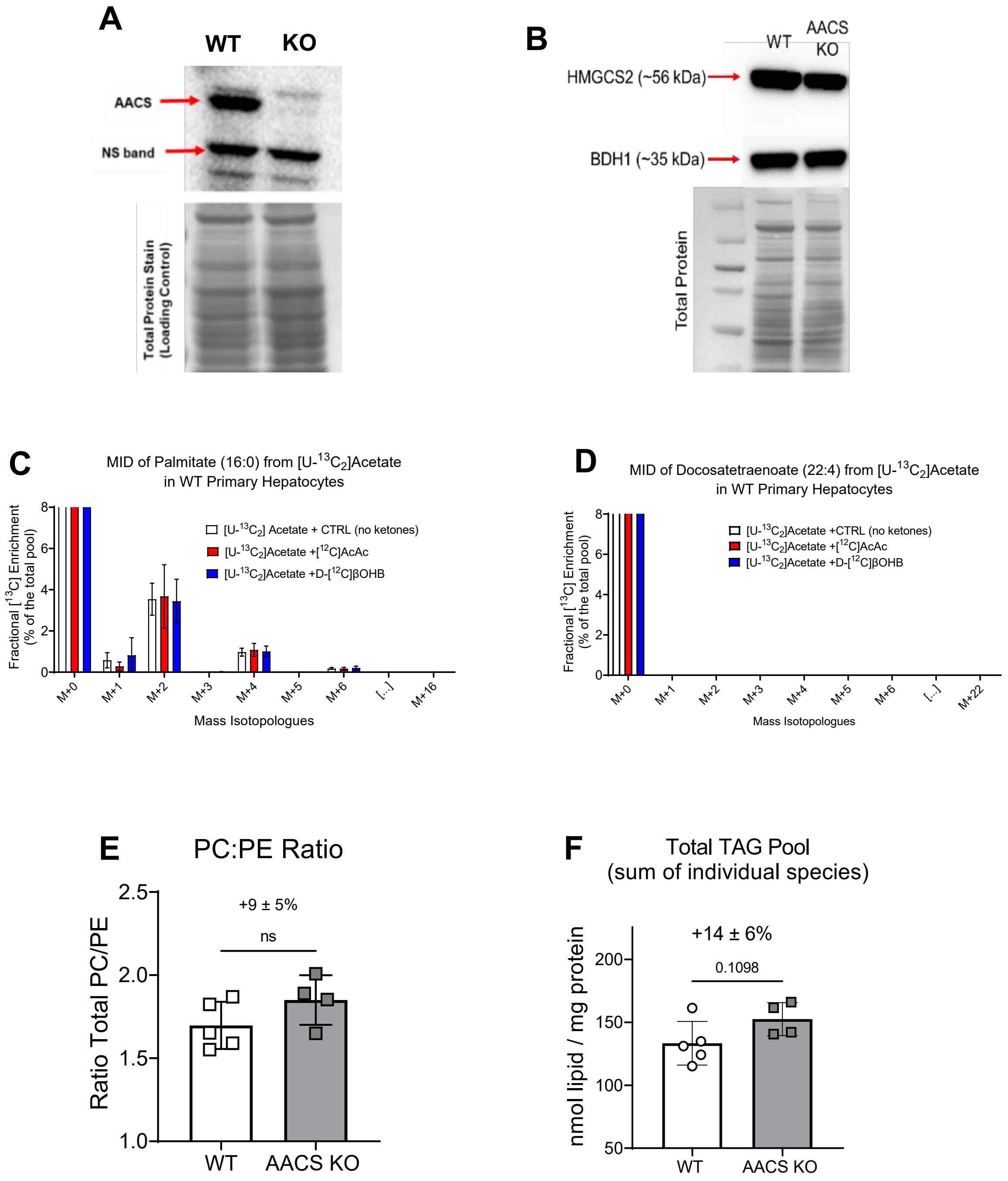

**Supplementary Figure 8. Loss of AACS reduces AcAc entry into DNL and FA elongation.**

**(A)** Immunoblot for AACS protein migrating at ~76 kDa in WT and AACS KO primary mouse hepatocytes NS = non-specific band. Total protein was quantified via FastStain. **(B)** Immunoblot for BDH1 and HMGCS2 proteins migrating at 35 kDa and 56 kDa, respectively, in WT and AACS KO primary hepatocytes. Total protein was quantified via FastStain. **(C)** Fractional [$^{13}$C] enrichment of palmitate and **(D)** MID of docosatetraenoate (22:4) from 1 mM [U-$^{13}C_2$]acetate in the absence (control), or presence of naturally-occurring 1 mM AcAc or D-βOHB (n=4/group). **(E)** The PC:PE ratio and **(F)** Total triacylglycerol (TAG) pool size in WT and AACS KO (n=4-5/group).

Data are expressed a mean ± SD. Statistical differences between genotypes were determined by Student's t test, and accepted as significant if $p<0.05$. *p < 0.05; **p < 0.01; ***p < 0.001; ****p < 0.0001; as indicated. Percent change and error propagation are given as X±Y% on graphs when relevant.

**Supplemental Figure 9**

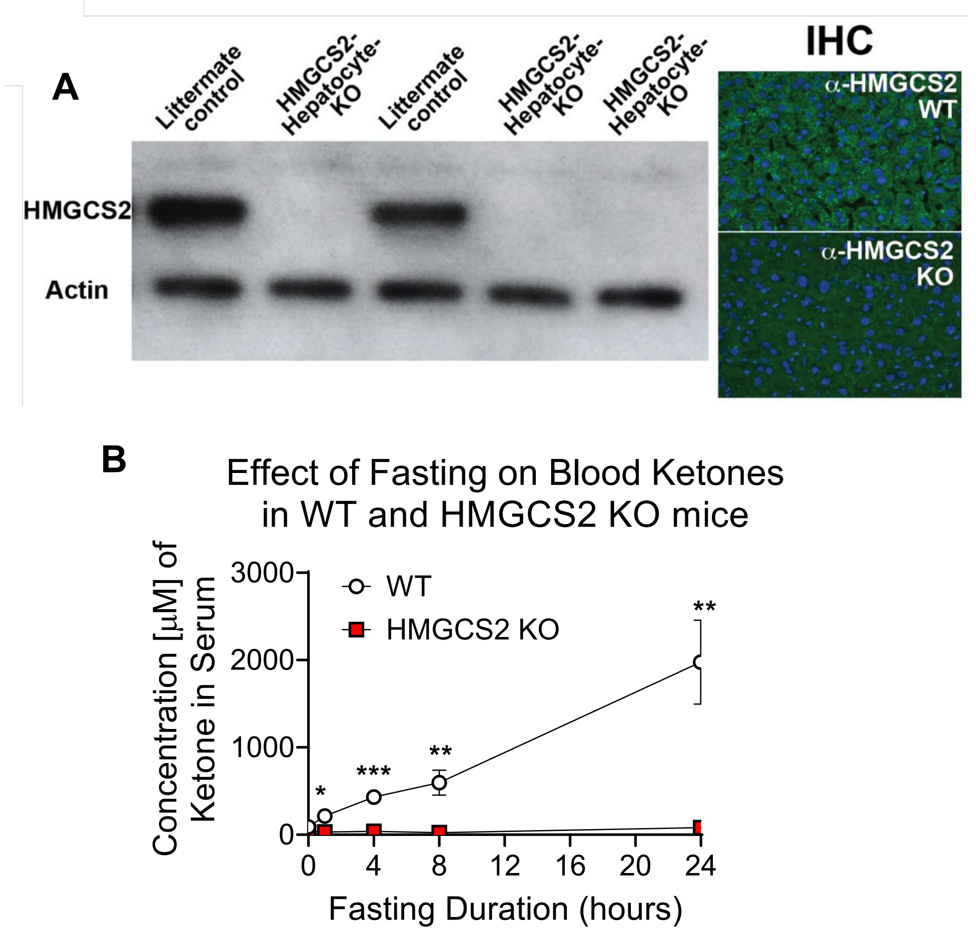

### Supplementary Figure 9. Validation of HMGCS2 KO Model

**(A)** Immunoblot of liver lysate, and immunohistochemistry (IHC) of liver tissue, confirming successful deletion of HMGCS2 protein. Actin protein used as loading control. **(B)** Concentration (μM) of total ketone bodies in serum during a 24 hour fast, in WT and HMGCS2 KO mice (n=2-3/group).

Data are expressed a mean ± SD. Statistical differences between genotypes were determined by Student's t test, and accepted as significant if $p<0.05$. *$p < 0.05$; **$p < 0.01$; ***$p < 0.001$; ****$p < 0.0001$; as indicated.

**Table 1. List of $^{13}$C-enriched chemical features from D-[U-$^{13}C_4$]βOHB detected across tissues *in vivo***

List of unique chemical features (retention time (RT) and mass/charge (m/z) pairs) detected in each tissue analyzed from mice injected with D-[U-13C4]βOHB (10 µmol/gram body weight) intraperitoneally after 30 minutes.

| Tissues | Heart | | Kidney | | Pancreas | | Spleen | | Gastroc | | Soleus | | Liver | | Brain | | Gonads | |
|---|---|---|---|---|---|---|---|---|---|---|---|---|---|---|---|---|---|---|
| **Chemical Features (RT + m/z pair)** | **Molecular Weight** | **RT [min]** | **Molecular Weight** | **RT [min]** | **Molecular Weight** | **RT [min]** | **Molecular Weight** | **RT [min]** | **Molecular Weight** | **RT [min]** | **Molecular Weight** | **RT [min]** | **Molecular Weight** | **RT [min]** | **Molecular Weight** | **RT [min]** | **Molecular Weight** | **RT [min]** |
| **1** | 74.037 | 33.1 | 72.021 | 34.0 | 74.000 | 38.0 | 89.048 | 16.2 | 100.016 | 35.9 | 74.037 | 35.9 | 89.048 | 24.5 | 90.032 | 14.681 | 90.032 | 14.818 |
| **2** | 89.048 | 24.8 | 74.037 | 33.3 | 89.048 | 16.2 | 89.048 | 26.4 | 103.063 | 26.4 | 89.048 | 26.9 | 100.016 | 33.5 | 103.063 | 25.358 | 103.063 | 25.217 |
| **3** | 90.032 | 15.3 | 86.037 | 46.9 | 89.048 | 27.0 | 90.032 | 15.1 | 104.047 | 14.9 | 100.016 | 35.9 | 102.079 | 17.2 | 103.063 | 18.329 | 104.047 | 14.915 |
| **4** | 103.063 | 24.9 | 89.048 | 16.1 | 90.032 | 15.2 | 100.016 | 37.7 | 104.047 | 13.0 | 102.032 | 35.7 | 103.063 | 24.6 | 104.047 | 12.951 | 118.027 | 34.87 |
| **5** | 104.011 | 33.8 | 89.048 | 24.7 | 100.016 | 37.8 | 103.063 | 18.2 | 112.016 | 48.6 | 103.063 | 26.6 | 104.047 | 14.9 | 128.059 | 17.294 | 129.043 | 17.645 |
| **6** | 104.047 | 15.7 | 90.032 | 14.7 | 103.063 | 26.9 | 103.063 | 26.4 | 116.011 | 36.9 | 104.047 | 15.1 | 104.047 | 13.5 | 129.043 | 17.365 | 133.038 | 25.291 |
| **7** | 104.047 | 14.3 | 103.063 | 24.7 | 104.047 | 37.5 | 116.011 | 37.9 | 118.027 | 35.8 | 104.047 | 35.7 | 116.011 | 37.9 | 129.043 | 25.322 | 134.022 | 34.875 |
| **8** | 116.011 | 33.3 | 104.011 | 34.0 | 104.047 | 15.9 | 116.011 | 26.4 | 128.059 | 17.4 | 112.016 | 48.7 | 118.027 | 33.1 | 132.053 | 17.66 | 136.037 | 34.616 |
| **9** | 117.043 | 18.2 | 104.047 | 15.0 | 105.043 | 18.1 | 116.989 | 5.9 | 129.043 | 18.3 | 116.011 | 36.9 | 128.059 | 17.3 | 133.038 | 25.682 | 146.069 | 17.326 |
| **10** | 118.027 | 33.1 | 104.047 | 13.6 | 112.016 | 50.6 | 118.027 | 37.7 | 130.027 | 35.6 | 116.011 | 36.2 | 129.043 | 17.5 | 134.022 | 35.843 | 147.053 | 25.298 |
| **11** | 129.043 | 18.4 | 106.027 | 19.0 | 115.027 | 17.7 | 129.043 | 18.3 | 130.027 | 48.7 | 118.027 | 35.9 | 129.043 | 24.6 | 146.069 | 17.277 | 148.037 | 34.652 |
| **12** | 133.038 | 24.8 | 112.016 | 46.3 | 115.063 | 14.3 | 129.043 | 17.2 | 133.038 | 26.9 | 128.059 | 17.6 | 130.027 | 32.9 | 147.053 | 25.365 | 162.053 | 34.314 |
| **13** | 134.022 | 33.2 | 116.011 | 33.5 | 116.011 | 38.0 | 133.038 | 26.4 | 134.022 | 36.1 | 130.027 | 35.7 | 133.038 | 24.5 | 147.066 | 17.153 | 167.025 | 12.151 |
| **14** | 146.022 | 34.8 | 117.043 | 17.5 | 118.027 | 37.8 | 134.022 | 37.9 | 146.069 | 17.4 | 130.027 | 49.0 | 134.022 | 33.4 | 174.064 | 19.617 | 173.105 | 13.022 |
| **15** | 146.069 | 17.6 | 118.027 | 33.3 | 129.043 | 12.9 | 146.069 | 17.2 | 147.053 | 26.4 | 133.038 | 27.0 | 145.074 | 14.0 | 188.009 | 14.402 | 174.064 | 19.563 |
| **16** | 147.053 | 24.9 | 128.059 | 17.2 | 129.043 | 18.4 | 147.053 | 26.4 | 148.037 | 35.6 | 134.022 | 36.2 | 146.069 | 17.2 | 188.080 | 19.185 | 175.048 | 34.834 |
| **17** | 148.037 | 32.8 | 129.043 | 17.7 | 129.043 | 17.4 | 148.037 | 37.3 | 174.016 | 48.7 | 146.069 | 17.6 | 147.053 | 24.5 | 192.027 | 48.923 | 188.009 | 14.158 |
| **18** | 162.053 | 32.3 | 129.043 | 13.5 | 130.027 | 50.6 | 162.053 | 36.7 | 192.027 | 48.6 | 147.053 | 26.5 | 147.053 | 18.1 | 199.051 | 18.341 | 188.080 | 19.087 |
| **19** | 167.025 | 12.5 | 132.054 | 17.6 | 132.054 | 17.7 | 167.025 | 12.6 | 196.040 | 48.5 | 148.037 | 35.8 | 148.037 | 32.9 | 202.045 | 14.601 | 191.062 | 13.107 |
| **20** | 169.035 | 24.9 | 133.038 | 24.7 | 132.090 | 18.3 | 169.035 | 26.3 | 226.157 | 1.1 | 162.053 | 35.3 | 161.069 | 17.1 | 209.065 | 17.291 | 192.027 | 50.546 |
| **21** | 171.051 | 16.5 | 134.022 | 33.5 | 133.038 | 27.0 | 172.035 | 15.1 | 253.062 | 33.1 | 167.025 | 13.0 | 162.053 | 32.3 | 292.138 | 17.3 | 202.045 | 14.32 |
| **22** | 173.105 | 14.0 | 136.038 | 19.2 | 134.022 | 38.0 | 172.049 | 36.2 | 290.123 | 29.2 | 174.016 | 48.9 | 167.025 | 11.8 | 294.106 | 25.347 | 207.090 | 12.938 |
| **23** | 174.016 | 46.5 | 145.074 | 14.0 | 146.069 | 17.4 | 175.096 | 18.2 | 291.992 | 35.8 | 175.048 | 35.4 | 174.064 | 19.3 | 302.072 | 25.664 | 218.019 | 14.721 |
| **24** | 181.041 | 11.1 | 146.022 | 35.1 | 147.053 | 26.9 | 180.064 | 7.5 | 292.138 | 17.4 | 186.064 | 34.2 | 175.096 | 18.1 | 314.120 | 17.282 | 231.047 | 17.35 |
| **25** | 182.058 | 16.1 | 146.069 | 17.3 | 148.037 | 37.5 | 186.064 | 35.6 | 294.106 | 26.4 | 192.027 | 48.7 | 185.080 | 16.0 | 320.034 | 17.322 | 275.069 | 12.987 |
| **26** | 185.080 | 16.6 | 147.053 | 24.7 | 161.069 | 17.2 | 188.080 | 19.9 | 314.120 | 17.4 | 196.040 | 49.0 | 186.064 | 26.0 | 326.132 | 17.138 | 276.132 | 27.011 |
| **27** | 186.064 | 31.5 | 148.037 | 33.0 | 163.030 | 13.7 | 202.045 | 15.1 | 316.088 | 26.4 | 201.064 | 35.4 | 188.080 | 19.0 | 330.094 | 17.278 | 292.138 | 17.32 |
| **28** | 187.025 | 16.4 | 159.090 | 13.6 | 167.025 | 12.7 | 218.019 | 15.1 | 320.034 | 17.4 | 209.065 | 17.6 | 189.064 | 32.2 | 343.000 | 25.048 | 294.106 | 25.123 |
| **29** | 195.057 | 10.4 | 161.069 | 13.5 | 174.016 | 51.1 | 218.090 | 19.5 | 321.018 | 26.3 | 211.051 | 13.2 | 192.027 | 47.6 | 394.109 | 1.015 | 302.072 | 25.155 |
| **30** | 202.025 | 15.6 | 172.014 | 35.3 | 174.064 | 20.9 | 219.075 | 36.4 | 321.018 | 26.5 | 218.015 | 26.9 | 196.058 | 21.0 | 446.376 | 1.097 | 316.088 | 25.19 |
| **31** | 202.045 | 15.3 | 173.105 | 13.3 | 175.096 | 18.3 | 232.035 | 15.0 | 330.094 | 17.4 | 232.131 | 1.3 | 197.080 | 19.8 | 482.171 | 17.307 | 320.034 | 17.338 |
| **32** | 216.051 | 16.4 | 174.016 | 46.9 | 188.009 | 15.1 | 250.011 | 14.0 | 332.062 | 26.4 | 238.193 | 11.0 | 201.137 | 13.0 | 498.145 | 17.303 | 323.014 | 25.749 |
| **33** | 223.034 | 14.0 | 175.096 | 18.1 | 188.080 | 20.1 | 253.073 | 18.0 | 338.070 | 26.4 | 275.083 | 10.3 | 204.075 | 26.0 | | | 330.094 | 17.341 |
| **34** | 225.031 | 31.3 | 184.147 | 11.8 | 188.116 | 16.1 | 254.057 | 33.8 | 385.964 | 36.1 | 276.029 | 36.3 | 217.106 | 19.9 | | | 403.172 | 17.669 |
| **35** | 228.072 | 17.5 | 186.064 | 26.2 | 189.100 | 14.8 | 268.081 | 11.8 | | | 284.118 | 9.7 | 218.090 | 25.4 | | | 420.016 | 21.723 |
| **36** | 230.077 | 15.6 | 187.121 | 13.1 | 191.062 | 13.4 | 276.096 | 39.9 | | | 287.068 | 19.7 | 218.090 | 18.7 | | | 445.931 | 35.622 |
| **37** | 232.035 | 15.5 | 188.080 | 19.3 | 191.080 | 21.0 | 288.057 | 26.5 | | | 288.057 | 27.0 | 223.034 | 13.2 | | | 525.283 | 1.238 |
| **38** | 237.995 | 33.3 | 189.064 | 32.3 | 192.027 | 50.6 | 290.123 | 29.6 | | | 289.976 | 37.1 | 225.031 | 31.2 | | | 865.582 | 13.226 |
| **39** | 238.193 | 11.7 | 190.084 | 13.6 | 194.079 | 15.1 | 291.992 | 37.6 | | | 290.123 | 29.2 | 236.999 | 33.4 | | | | |
| **40** | 253.062 | 30.6 | 192.027 | 46.3 | 197.080 | 21.2 | 294.106 | 26.4 | | | 291.992 | 36.0 | 237.995 | 33.4 | | | | |
| **41** | 254.057 | 30.5 | 196.058 | 21.2 | 201.137 | 12.8 | 301.057 | 39.5 | | | 292.138 | 17.6 | 244.046 | 17.1 | | | | |
| **42** | 254.188 | 11.1 | 197.046 | 13.5 | 202.045 | 15.2 | 304.031 | 26.5 | | | 294.106 | 26.9 | 248.101 | 25.7 | | | | |
| **43** | 261.121 | 24.0 | 200.066 | 13.9 | 205.095 | 19.8 | 304.240 | 1.1 | | | 304.031 | 26.9 | 251.075 | 19.2 | | | | |
| **44** | 273.073 | 15.8 | 202.025 | 15.0 | 205.095 | 19.0 | 307.002 | 26.6 | | | 306.035 | 37.4 | 253.073 | 17.6 | | | | |
| **45** | 280.090 | 24.8 | 202.045 | 14.6 | 207.090 | 13.1 | 309.106 | 30.6 | | | 307.002 | 27.1 | 254.188 | 10.2 | | | | |
| **46** | 287.068 | 19.9 | 204.075 | 26.2 | 216.051 | 16.3 | 312.107 | 1.0 | | | 309.999 | 36.3 | 259.010 | 13.2 | | | | |
| **47** | 287.068 | 21.7 | 218.019 | 14.5 | 217.106 | 21.3 | 318.047 | 26.4 | | | 314.120 | 17.6 | 275.112 | 25.7 | | | | |
| **48** | 291.992 | 33.0 | 218.090 | 25.6 | 218.019 | 15.2 | 320.034 | 17.2 | | | 316.088 | 26.6 | 276.132 | 26.7 | | | | |
| **49** | 294.106 | 24.9 | 219.075 | 32.3 | 242.019 | 23.1 | 328.094 | 22.0 | | | 320.034 | 17.7 | 280.090 | 24.5 | | | | |
| **50** | 304.058 | 17.5 | 223.034 | 13.3 | 249.030 | 49.7 | 332.062 | 26.3 | | | 321.018 | 26.4 | 288.057 | 24.4 | | | | |
| **51** | 308.122 | 22.4 | 225.031 | 31.4 | 250.095 | 13.8 | 358.141 | 17.2 | | | 322.002 | 35.9 | 289.127 | 25.1 | | | | |
| **52** | 309.106 | 28.8 | 227.152 | 12.8 | 275.112 | 28.1 | 359.125 | 26.3 | | | 330.094 | 17.6 | 289.976 | 33.8 | | | | |
| **53** | 309.999 | 33.2 | 234.086 | 26.5 | 276.096 | 39.9 | 383.108 | 39.4 | | | 332.062 | 26.6 | 291.992 | 33.1 | | | | |
| **54** | 314.120 | 17.5 | 238.193 | 11.1 | 276.132 | 28.5 | 384.980 | 26.6 | | | 369.969 | 36.0 | 292.138 | 17.2 | | | | |
| **55** | 316.119 | 16.0 | 248.101 | 25.8 | 283.073 | 24.5 | 398.996 | 26.2 | | | 387.976 | 36.4 | 294.106 | 24.5 | | | | |
| **56** | 320.034 | 17.6 | 250.063 | 25.7 | 287.068 | 19.6 | 525.307 | 15.3 | | | 463.074 | 55.3 | 295.091 | 28.0 | | | | |
| **57** | 321.018 | 24.7 | 253.002 | 14.5 | 288.057 | 27.1 | 591.892 | 13.7 | | | 796.146 | 27.1 | 302.072 | 24.5 | | | | |
| **58** | 321.018 | 25.1 | 254.057 | 24.9 | 288.061 | 13.2 | | | | | | | 308.122 | 21.7 | | | | |
| **59** | 322.002 | 32.8 | 254.188 | 10.7 | 291.992 | 37.8 | | | | | | | 309.106 | 28.2 | | | | |
| **60** | 330.094 | 17.5 | 270.095 | 14.6 | 302.072 | 27.0 | | | | | | | 309.999 | 33.4 | | | | |
| **61** | 332.062 | 25.0 | 275.112 | 26.4 | 304.058 | 17.4 | | | | | | | 314.120 | 17.2 | | | | |
| **62** | 385.964 | 33.4 | 276.096 | 35.7 | 307.002 | 27.1 | | | | | | | 316.088 | 24.5 | | | | |
| **63** | 385.964 | 33.0 | 276.132 | 26.8 | 309.106 | 30.8 | | | | | | | 320.034 | 17.2 | | | | |
| **64** | 390.106 | 17.5 | 283.073 | 22.6 | 309.106 | 24.2 | | | | | | | 322.002 | 32.9 | | | | |
| **65** | 445.931 | 33.3 | 287.068 | 19.3 | 320.034 | 17.4 | | | | | | | 326.132 | 17.4 | | | | |
| **66** | 476.154 | 17.6 | 289.976 | 34.0 | 321.018 | 27.1 | | | | | | | 330.094 | 17.2 | | | | |
| **67** | 477.957 | 32.8 | 291.992 | 33.2 | 321.018 | 26.8 | | | | | | | 331.138 | 30.8 | | | | |
| **68** | | | 292.138 | 17.3 | 330.094 | 17.3 | | | | | | | 345.197 | 2.7 | | | | |
| **69** | | | 294.106 | 24.7 | 358.141 | 17.4 | | | | | | | 349.095 | 32.1 | | | | |
| **70** | | | 303.008 | 18.0 | 369.969 | 37.8 | | | | | | | 350.098 | 32.1 | | | | |
| **71** | | | 307.084 | 26.9 | 389.156 | 17.7 | | | | | | | 357.110 | 19.4 | | | | |
| **72** | | | 309.106 | 22.3 | 398.996 | 26.8 | | | | | | | 363.029 | 32.2 | | | | |
| **73** | | | 309.107 | 28.4 | 463.074 | 56.9 | | | | | | | 372.215 | 13.4 | | | | |
| **74** | | | 309.999 | 33.5 | 566.055 | 38.5 | | | | | | | 390.298 | 1.2 | | | | |
| **75** | | | 310.095 | 28.1 | 572.368 | 17.4 | | | | | | | 399.980 | 32.9 | | | | |
| **76** | | | 321.018 | 24.5 | 605.077 | 42.1 | | | | | | | 403.135 | 19.2 | | | | |
| **77** | | | 322.002 | 33.0 | 607.082 | 37.6 | | | | | | | 403.172 | 17.5 | | | | |
| **78** | | | 344.028 | 35.3 | 861.473 | 13.8 | | | | | | | 408.191 | 13.4 | | | | |
| **79** | | | 346.069 | 22.6 | | | | | | | | | 409.098 | 29.7 | | | | |
| **80** | | | 369.969 | 33.1 | | | | | | | | | 422.207 | 13.3 | | | | |
| **81** | | | 384.112 | 35.2 | | | | | | | | | 496.126 | 31.1 | | | | |
| **82** | | | 385.964 | 33.4 | | | | | | | | | 514.119 | 17.3 | | | | |
| **83** | | | 386.231 | 13.3 | | | | | | | | | | | | | | |
| **84** | | | 403.172 | 17.4 | | | | | | | | | | | | | | |
| **85** | | | 423.957 | 35.2 | | | | | | | | | | | | | | |
| **86** | | | 426.088 | 34.0 | | | | | | | | | | | | | | |
| **87** | | | 451.102 | 22.6 | | | | | | | | | | | | | | |
| **88** | | | 457.038 | 22.6 | | | | | | | | | | | | | | |
| **89** | | | 555.132 | 38.5 | | | | | | | | | | | | | | |
| **90** | | | 567.060 | 34.1 | | | | | | | | | | | | | | |
| **91** | | | 580.035 | 45.6 | | | | | | | | | | | | | | |
| **92** | | | 607.082 | 33.4 | | | | | | | | | | | | | | |
| **93** | | | 700.531 | 1.0 | | | | | | | | | | | | | | |

**<u>Table 2. Absolute Pool Size of FFAs in WT and AACS KO Primary Hepatocytes</u>**

Average and SD for each individual FFA species detected via shotgun lipidomics. Fold change is the average of the KO divided by the average of the WT. Significant differences determined by unpaired two-way Student's t test between WT and AACS KO with Holm-Sidak multi-comparisions correction.

| Shotgun Lipidomics Free Fatty Acid (FFA) Pool Sizes (nmol lipid / mg protein) | | | | | | | |
|---|---|---|---|---|---|---|---|
| **FFA Species** | **WT** | | **AACS KO** | | **Fold Change** | **Raw P Value** | **Adjusted P-Value** |
| | Average | ±SD | Average | ±SD | | | |
| **14:0** | 6.9 | 1.2 | 7.6 | 1.1 | 1.10 | 0.4760 | 0.8933 |
| **16:1** | 38.7 | 4.0 | 40.1 | 8.9 | 1.04 | 0.7903 | 0.8968 |
| **16:0** | 363.7 | 42.6 | 336.3 | 43.4 | 0.92 | 0.4285 | 0.8933 |
| **18:3** | 14.1 | 1.8 | 7.3 | 2.2 | 0.52 | ***0.0032*** | ***0.0437*** |
| **18:2** | 269.8 | 24.6 | 140.6 | 37.7 | 0.52 | ***0.0009*** | ***0.0169*** |
| **18:1** | 336.0 | 41.7 | 290.3 | 46.8 | 0.86 | 0.2141 | 0.8149 |
| **18:0** | 173.4 | 38.0 | 183.9 | 21.7 | 1.06 | 0.6788 | 0.8968 |
| **20:5** | 4.5 | 0.4 | 3.4 | 0.7 | 0.75 | ***0.0353*** | 0.2766 |
| **20:4** | 116.3 | 9.9 | 78.7 | 14.7 | 0.68 | ***0.0050*** | *0.0634* |
| **20:3** | 19.0 | 1.6 | 10.0 | 3.0 | 0.53 | ***0.0014*** | ***0.0217*** |
| **20:2** | 6.7 | 0.4 | 3.2 | 0.8 | 0.48 | ***0.0001*** | ***0.0028*** |
| **20:1** | 7.2 | 0.5 | 5.6 | 1.2 | 0.79 | *0.0550* | 0.3638 |
| **20:0** | 3.6 | 0.4 | 2.6 | 0.3 | 0.73 | ***0.0083*** | *0.0957* |
| **22:6** | 97.6 | 9.3 | 67.7 | 17.1 | 0.69 | ***0.0215*** | 0.2128 |
| **22:5** | 12.5 | 1.3 | 8.4 | 2.5 | 0.67 | ***0.0234*** | 0.2128 |
| **22:4** | 8.8 | 0.9 | 4.1 | 1.5 | 0.46 | ***0.0013*** | ***0.0212*** |
| **24:6** | 1.4 | 0.3 | 1.0 | 0.5 | 0.74 | 0.2484 | 0.8197 |
| **24:5** | 2.5 | 0.4 | 1.1 | 0.4 | 0.42 | ***0.0020*** | ***0.0295*** |
| **24:4** | 2.0 | 0.4 | 0.2 | 0.1 | 0.11 | ***0.0001*** | ***0.0018*** |

**Table 3. Absolute Pool Size of PEs WT and AACS KO Primary Hepatocytes**

Average and SD for each individual PE species detected via shotgun lipidomics. Fold change is the average of the KO divided by the average of the WT. Significant differences determined by unpaired two-way Student's t test between WT and AACS KO with Holm-Sidak multi-comparisions correction.

| Shotgun Lipidomics Phosphatidylethanolamine (PE) Pool Sizes (nmol lipid / mg protein) | | | | | | | |
|---|---|---|---|---|---|---|---|
| **PE Species** | **WT** | | **AACS KO** | | **Fold Change** | **Raw P Value** | **Adjusted P-Value** |
| | Average | ±SD | Average | ±SD | | | |
| **D16:0-16:1/D14:0-18:1** | 0.4 | 0.1 | 0.3 | 0.1 | 0.86 | 0.5850 | 0.9426 |
| **D16:0-18:2/D16:1-18:1** | 3.1 | 0.2 | 1.9 | 0.2 | 0.62 | ***0.0002*** | ***0.0033*** |
| **D16:0-18:1** | 0.6 | 0.1 | 0.6 | 0.2 | 1.14 | 0.4521 | 0.9426 |
| **D16:1-20:4** | 0.6 | 0.1 | 0.6 | 0.1 | 0.94 | 0.6685 | 0.9426 |
| **D16:0-20:4/D18:2-18:2** | 10.2 | 0.7 | 9.1 | 0.5 | 0.89 | *0.0531* | 0.4804 |
| **D18:1-18:2/D16:0-20:3** | 1.8 | 0.2 | 1.3 | 0.1 | 0.72 | ***0.0054*** | *0.0925* |
| **D18:0-18:2/D18:1-18:1/D16:0-20:2** | 4.7 | 0.4 | 2.8 | 0.3 | 0.60 | ***0.0001*** | ***0.0033*** |
| **D18:0-18:1/D16:0-20:1** | 0.5 | 0.1 | 0.4 | 0.0 | 0.68 | ***0.0028*** | *0.0511* |
| **P16:0-22:6/D18:0-18:0/P18:2-20:4** | 0.4 | 0.1 | 0.3 | 0.0 | 0.77 | 0.1134 | 0.7000 |
| **P18:0-20:3** | 0.5 | 0.0 | 0.5 | 0.1 | 0.92 | 0.4353 | 0.9426 |
| **D16:1-22:6** | 0.3 | 0.0 | 0.3 | 0.1 | 0.96 | 0.7652 | 0.9426 |
| **D16:0-22:6** | 13.5 | 1.2 | 9.4 | 0.2 | 0.69 | ***0.0004*** | ***0.0083*** |
| **D18:1-20:4D/16:0-22:5** | 8.2 | 0.3 | 6.2 | 0.1 | 0.76 | ***0.0000*** | ***0.0002*** |
| **D18:0-20:4/D16:0-22:4** | 42.4 | 2.0 | 37.8 | 1.9 | 0.89 | ***0.0188*** | 0.2339 |
| **D18:0-20:3/D18:1-20:2/D16:0-22:3** | 1.7 | 0.2 | 1.3 | 0.3 | 0.73 | ***0.0489*** | 0.4787 |
| **P18:2-22:6/D18:1-20:1** | 0.3 | 0.1 | 0.2 | 0.0 | 0.69 | 0.1961 | 0.7831 |
| **A20:0-20:4/P18:0-22:3** | 0.7 | 0.0 | 0.6 | 0.1 | 0.87 | 0.1175 | 0.7000 |
| **D18:2-22:6** | 0.4 | 0.1 | 0.2 | 0.1 | 0.51 | ***0.0127*** | 0.1847 |
| **D18:1-22:6** | 3.3 | 0.3 | 2.5 | 0.3 | 0.77 | ***0.0141*** | 0.1924 |
| **D18:0-22:6/D18:1-22:5** | 10.4 | 0.5 | 9.0 | 0.5 | 0.86 | ***0.0067*** | 0.1085 |
| **D18:0-22:5/D18:1-22:4** | 1.0 | 0.2 | 0.9 | 0.1 | 0.87 | 0.3272 | 0.9072 |
| **D20:0-20:4/D18:0-22:4** | 0.9 | 0.1 | 0.8 | 0.2 | 0.83 | 0.1494 | 0.7261 |
| **P22:6-22:6** | 0.2 | 0.0 | 0.1 | 0.0 | 0.70 | *0.0890* | 0.6412 |

**Table 4. Absolute Pool Size of PCs in WT and AACS KO Primary Hepatocytes**

Average and SD for each individual PC species detected via shotgun lipidomics. Fold change is the average of the KO divided by the average of the WT. Significant differences determined by unpaired two-way Student's t test between WT and AACS KO with Holm-Sidak multi-comparisions correction.

| Shotgun Lipidomics Phosphatidylcholine (PC) Pool Sizes (nmol lipid / mg protein) | | | | | | | |
|---|---|---|---|---|---|---|---|
| **PC Species** | **WT** | | **AACS KO** | | **Fold Change** | **Raw P Value** | **Adjusted P-Value** |
| | Average | ±SD | Average | ±SD | | | |
| **D16:0-18:2** | 38.9 | 4.9 | 29.4 | 1.4 | 0.76 | ***0.0125*** | 0.1401 |
| **D16:0-18:1** | 15.7 | 1.6 | 21.5 | 1.0 | 1.37 | ***0.0008*** | ***0.0120*** |
| **D16:0-18:0** | 7.8 | 1.2 | 10.1 | 0.3 | 1.30 | ***0.0268*** | 0.2585 |
| **D18:2-18:2/D16:0-20:4** | 26.9 | 3.2 | 27.9 | 2.3 | 1.04 | 0.6505 | 0.9664 |
| **D18:1-18:2/D16:0-20:3** | 8.8 | 0.7 | 9.3 | 1.3 | 1.05 | 0.6161 | 0.9664 |
| **D18:0-18:2/D18:1-18:1** | 17.0 | 2.3 | 11.6 | 1.2 | 0.68 | ***0.0072*** | *0.0963* |
| **D18:0-18:1** | 9.0 | 1.1 | 5.3 | 2.3 | 0.59 | ***0.0293*** | 0.2585 |
| **D18:2-20:5** | 0.8 | 0.2 | 0.4 | 0.1 | 0.50 | ***0.0110*** | 0.1334 |
| **D16:0-22:6/D18:2-20:4** | 11.6 | 1.3 | 11.5 | 0.8 | 0.99 | 0.9332 | 0.9664 |
| **D18:1-20:4/D16:0-22:5** | 3.9 | 0.9 | 5.1 | 1.8 | 1.32 | 0.2980 | 0.8295 |
| **D18:2-20:2/D18:0-20:4** | 14.1 | 1.9 | 12.3 | 1.1 | 0.87 | 0.1608 | 0.7069 |
| **D18:0-20:3** | 17.8 | 2.1 | 12.3 | 6.7 | 0.69 | 0.1677 | 0.7069 |
| **D18:0-20:2/P18:2-22:6** | 1.5 | 0.4 | 0.9 | 0.2 | 0.62 | ***0.0430*** | 0.2965 |
| **D18:1-22:6/D18:2-22:5** | 0.6 | 0.0 | 0.6 | 0.3 | 1.13 | 0.5718 | 0.9664 |
| **D18:0-22:5** | 5.9 | 0.7 | 4.7 | 0.3 | 0.81 | ***0.0382*** | 0.2956 |

**Table 5. Absolute Pool Size of TAGs in WT and AACS KO Primary Hepatocytes**

Average and SD for each individual TAG species detected via shotgun lipidomics. Fold change is the average of the KO divided by the average of the WT. Significant differences determined by unpaired two-way Student's t test between WT and AACS KO with Holm-Sidak multi-comparisions correction.

| Shotgun Lipidomics Triacylglycerol (TAG) Pool Sizes (nmol lipid / mg protein) | | | | | | | |
|---|---|---|---|---|---|---|---|
| **TAG Species** | **WT** | | **AACS KO** | | **Fold Change** | **Raw P Value** | **Adjusted P-Value** |
| | Average | ±SD | Average | ±SD | | | |
| **C50:3/C51:10** | 2.9 | 0.5 | 4.7 | 0.4 | 1.7 | ***0.0011*** | ***0.0293*** |
| **C50:2/C51:9** | 6.4 | 1.2 | 9.7 | 0.8 | 1.5 | ***0.0051*** | 0.1234 |
| **C50:1/C51:8** | 3.9 | 0.8 | 6.5 | 0.6 | 1.7 | ***0.0013*** | ***0.0345*** |
| **C52:5** | 2.9 | 0.9 | 3.9 | 0.8 | 1.3 | 0.1688 | 0.9481 |
| **C52:4/C53:11** | 9.7 | 1.3 | 11.0 | 0.7 | 1.1 | 0.1508 | 0.9379 |
| **C52:3/C53:10** | 16.1 | 2.3 | 19.3 | 1.4 | 1.2 | *0.0684* | 0.7577 |
| **C52:2/C53:9** | 12.3 | 2.4 | 15.5 | 1.0 | 1.3 | *0.0577* | 0.7294 |
| **C52:1/C53:8** | 2.6 | 0.4 | 2.8 | 0.3 | 1.1 | 0.5883 | 0.9999 |
| **C53:0/C54:7** | 2.2 | 0.6 | 3.0 | 0.7 | 1.4 | 0.1348 | 0.9331 |
| **C54:6** | 6.1 | 1.1 | 6.6 | 0.8 | 1.1 | 0.5576 | 0.9999 |
| **C54:5/C55:12** | 7.3 | 0.5 | 7.3 | 1.0 | 1.0 | 0.9579 | 0.9999 |
| **C54:4/C55:11** | 8.4 | 0.9 | 8.6 | 0.5 | 1.0 | 0.8298 | 0.9999 |
| **C54:3/C55:10** | 7.7 | 1.0 | 9.4 | 0.6 | 1.2 | ***0.0332*** | 0.5406 |
| **C54:2/C55:9** | 3.7 | 0.3 | 3.3 | 0.2 | 0.9 | 0.1327 | 0.9331 |
| **C54:1/C55:8** | 1.7 | 0.2 | 1.5 | 0.2 | 0.9 | 0.1983 | 0.9637 |
| **C54:0/C55:7** | 2.5 | 1.1 | 2.9 | 0.8 | 1.2 | 0.5900 | 0.9999 |
| **C55:2/C56:9** | 1.6 | 0.4 | 1.9 | 0.5 | 1.1 | 0.5499 | 0.9999 |
| **C55:1/C56:8** | 4.7 | 0.4 | 5.0 | 0.7 | 1.0 | 0.5941 | 0.9999 |
| **C55:0/C56:7** | 5.5 | 0.4 | 6.5 | 0.5 | 1.2 | ***0.0260*** | 0.4825 |
| **C56:6** | 5.1 | 0.2 | 4.4 | 0.4 | 0.9 | ***0.0319*** | 0.5406 |
| **C56:5/C57:12** | 3.5 | 0.5 | 3.1 | 0.6 | 0.9 | 0.3896 | 0.9990 |
| **C56:4/C57:11** | 2.5 | 0.5 | 2.3 | 0.6 | 0.9 | 0.7046 | 0.9999 |
| **C56:3/C57:10** | 2.5 | 0.4 | 2.3 | 0.2 | 0.9 | 0.4683 | 0.9997 |
| **C56:2/C57:9** | 1.8 | 0.3 | 1.4 | 0.1 | 0.8 | 0.0582 | 0.7294 |
| **C57:3/C58:10** | 1.6 | 0.2 | 1.6 | 0.2 | 1.0 | 0.6683 | 0.9999 |
| **C57:2/C58:9** | 2.4 | 0.4 | 2.4 | 0.1 | 1.0 | 0.7865 | 0.9999 |
| **C57:1/C58:8** | 3.2 | 0.3 | 3.3 | 0.1 | 1.0 | 0.7787 | 0.9999 |
| **C57:0/C58:7/C59:14** | 2.6 | 0.5 | 2.5 | 0.3 | 1.0 | 0.7688 | 0.9999 |

**Table 6. Absolute Pool Size of FFAs in WT and HMGCS2 KO Primary Hepatocytes**

Average and SD for each individual FFA species detected via shotgun lipidomics. Fold change is the average of the KO divided by the average of the WT. Significant differences determined by unpaired two-way Student's t test between WT and HMGCS2 KO with Holm-Sidak multi-comparisions correction.

| Shotgun Lipidomics Free Fatty Acid (FFA) Pool Sizes (nmol lipid / mg protein) | | | | | | | |
|---|---|---|---|---|---|---|---|
| **FFA Species** | **WT** | | **HMGCS2 KO** | | **Fold Change** | **Raw P Value** | **Adjusted P-Value** |
| | Average | ±SD | Average | ±SD | | | |
| **14:0** | 0.7 | 0.2 | 1.7 | 0.4 | 2.6 | ***0.0004*** | ***0.0078*** |
| **16:2** | 0.2 | 0.1 | 1.3 | 0.4 | 5.4 | ***0.0002*** | ***0.0042*** |
| **16:1** | 3.3 | 1.0 | 13.4 | 5.2 | 4.1 | ***0.0013*** | ***0.0271*** |
| **16:0** | 26.1 | 3.3 | 38.9 | 10.1 | 1.5 | ***0.0189*** | 0.2910 |
| **18:3** | 3.2 | 0.6 | 7.3 | 2.9 | 2.3 | ***0.0093*** | 0.1706 |
| **18:2** | 34.8 | 6.2 | 55.3 | 20.4 | 1.6 | *0.0516* | 0.5716 |
| **18:1** | 17.6 | 3.4 | 36.9 | 14.0 | 2.1 | ***0.0117*** | 0.1996 |
| **18:0** | 6.1 | 0.9 | 7.0 | 1.0 | 1.2 | 0.1237 | 0.8643 |
| **20:5** | 0.5 | 0.1 | 0.5 | 0.2 | 1.0 | 0.8403 | 0.9998 |
| **20:4** | 2.7 | 0.6 | 3.2 | 0.6 | 1.2 | 0.1853 | 0.9422 |
| **20:3** | 0.6 | 0.3 | 0.5 | 0.2 | 0.9 | 0.7213 | 0.9998 |
| **20:2** | 0.3 | 0.1 | 0.4 | 0.2 | 1.2 | 0.5763 | 0.9989 |
| **20:1** | 0.7 | 0.3 | 0.7 | 0.3 | 1.1 | 0.7492 | 0.9998 |
| **20:0** | 0.4 | 0.1 | 0.7 | 0.3 | 1.8 | ***0.0236*** | 0.3247 |
| **22:6** | 2.1 | 0.7 | 2.1 | 0.4 | 1.0 | 0.9946 | 0.9998 |
| **22:5** | 0.8 | 0.2 | 0.7 | 0.2 | 0.8 | 0.2867 | 0.9788 |
| **22:4** | 0.4 | 0.2 | 0.3 | 0.2 | 0.8 | 0.3630 | 0.9796 |
| **22:3** | 1.7 | 0.5 | 1.8 | 1.7 | 1.1 | 0.8922 | 0.9998 |
| **22:0** | 0.2 | 0.1 | 0.3 | 0.2 | 1.5 | 0.2387 | 0.9712 |
| **24:6** | 0.0 | 0.0 | 0.1 | 0.1 | 2.0 | 0.3295 | 0.9788 |
| **24:5** | 0.1 | 0.0 | 0.2 | 0.2 | 1.9 | 0.2743 | 0.9772 |
| **24:4** | 0.8 | 0.3 | 0.7 | 0.5 | 0.9 | 0.7729 | 0.9998 |
| **24:0** | 1.8 | 0.5 | 1.8 | 1.7 | 1.0 | 0.9705 | 0.9998 |

**Table 7. Absolute Pool Size of PEs in WT and HMGCS2 KO Primary Hepatocytes**

Average and SD for each individual FFA species detected via shotgun lipidomics. Fold change is the average of the KO divided by the average of the WT. Significant differences determined by unpaired two-way Student's t test between WT and HMGCS2 KO with Holm-Sidak multi-comparisions correction.

| Shotgun Lipidomics Phosphatidylethanolamine (PE) Pool Sizes (nmol lipid / mg protein) | | | | | | | |
|---|---|---|---|---|---|---|---|
| PE Species | WT | | HMGCS2 KO | | Fold Change | Raw P Value | Adjusted P-Value |
| | Average | ±SD | Average | ±SD | | | |
| D16:0-16:1/D14:0-18:1 | 0.4 | 0.1 | 0.4 | 0.2 | 1.02 | 0.9299 | 1.0000 |
| P18:1-16:0/P16:0-18:1 | 0.1 | 0.0 | 0.2 | 0.1 | 1.43 | 0.4161 | 1.0000 |
| P18:0-16:0/P16:0-18:0 | 0.1 | 0.0 | 0.1 | 0.0 | 1.11 | 0.6509 | 1.0000 |
| D16:1-18:2 | 0.3 | 0.2 | 0.8 | 0.2 | 2.46 | ***0.0019*** | *0.0721* |
| D16:0-18:2/D16:1-18:1 | 5.2 | 1.7 | 6.1 | 1.8 | 1.18 | 0.3821 | 0.9999 |
| D16:0-18:1 | 0.5 | 0.1 | 0.7 | 0.3 | 1.34 | 0.2084 | 0.9976 |
| P16:0-20:4 | 0.5 | 0.1 | 0.7 | 0.6 | 1.50 | 0.3412 | 0.9999 |
| P16:0-20:3/P18:1-18:2 | 0.1 | 0.0 | 0.1 | 0.0 | 1.01 | 0.9838 | 1.0000 |
| P18:1-18:1/P18:0-18:2/P16:0-20:2 | 0.1 | 0.0 | 0.2 | 0.1 | 2.20 | 0.1400 | 0.9924 |
| P18:0-18:1/P16:0-20:1 | 0.2 | 0.0 | 0.2 | 0.0 | 1.22 | 0.1100 | 0.9806 |
| D16:1-20:4 | 0.4 | 0.1 | 0.6 | 0.1 | 1.70 | ***0.0035*** | 0.1172 |
| D16:0-20:4/D18:2-18:2 | 6.3 | 1.0 | 7.5 | 1.8 | 1.20 | 0.1941 | 0.9976 |
| D18:1-18:2/D16:0-20:3 | 2.3 | 0.5 | 4.5 | 0.9 | 1.95 | ***0.0003*** | ***0.0138*** |
| D18:0-18:2/D18:1-18:1/D16:0-20:2 | 3.4 | 0.9 | 5.6 | 1.4 | 1.66 | ***0.0084*** | 0.2666 |
| D18:0-18:1/D16:0-20:1 | 0.2 | 0.1 | 0.3 | 0.2 | 1.62 | 0.1989 | 0.9976 |
| P16:0-22:6/D18:0-18:0/P18:2-20:4 | 0.2 | 0.1 | 0.5 | 0.4 | 1.91 | 0.3012 | 0.9997 |
| P18:1-20:4/P16:0-22:5 | 0.3 | 0.2 | 0.4 | 0.2 | 1.14 | 0.7253 | 1.0000 |
| P18:0-20:4/P16:0-22:4/P18:1-20:3 | 0.5 | 0.1 | 0.6 | 0.4 | 1.31 | 0.4558 | 1.0000 |
| P18:0-20:3 | 0.4 | 0.0 | 0.4 | 0.2 | 1.17 | 0.4700 | 1.0000 |
| P20:1-18:1/P18:1-20:1 | 0.1 | 0.1 | 0.2 | 0.1 | 1.69 | 0.1916 | 0.9976 |
| D16:1-22:6 | 0.3 | 0.1 | 0.4 | 0.1 | 1.21 | 0.3276 | 0.9999 |
| D16:0-22:6 | 14.0 | 1.4 | 8.7 | 2.8 | 0.62 | ***0.0025*** | 0.0917 |
| D18:1-20:4D/16:0-22:5 | 4.2 | 0.9 | 4.7 | 0.9 | 1.11 | 0.4412 | 1.0000 |
| D18:0-20:4/D16:0-22:4 | 16.7 | 1.9 | 18.0 | 3.1 | 1.07 | 0.4588 | 1.0000 |
| D18:0-20:3/D18:1-20:2/D16:0-22:3 | 0.6 | 0.1 | 0.6 | 0.4 | 0.92 | 0.7542 | 1.0000 |
| P18:0-22:5/P18:1-22:4 | 0.4 | 0.1 | 0.3 | 0.1 | 0.80 | 0.1484 | 0.9924 |
| P18:0-22:4/P20:0-20:4/P18:1-22:3 | 0.3 | 0.1 | 0.3 | 0.1 | 0.87 | 0.5517 | 1.0000 |
| A20:0-20:4/P18:0-22:3 | 0.4 | 0.0 | 0.4 | 0.1 | 0.96 | 0.7004 | 1.0000 |
| P18:0-22:2 | 0.1 | 0.0 | 0.2 | 0.2 | 1.60 | 0.4219 | 1.0000 |
| D18:2-22:6 | 0.2 | 0.1 | 0.2 | 0.1 | 0.72 | 0.1521 | 0.9951 |
| D18:1-22:6 | 2.4 | 0.6 | 1.8 | 0.3 | 0.74 | ***0.0412*** | 0.7867 |
| D18:0-22:6/D18:1-22:5 | 4.7 | 0.6 | 3.9 | 1.1 | 0.83 | 0.1733 | 0.9961 |
| D18:0-22:5/D18:1-22:4 | 0.6 | 0.2 | 0.6 | 0.2 | 0.92 | 0.6650 | 1.0000 |
| D20:0-20:4/D18:0-22:4 | 0.3 | 0.1 | 0.3 | 0.1 | 1.07 | 0.7454 | 1.0000 |
| D20:0-20:0/D18:0-22:0 | 0.1 | 0.0 | 0.1 | 0.0 | 1.20 | 0.5077 | 1.0000 |
| D20:6-22:7 | 0.1 | 0.0 | 0.1 | 0.0 | 0.72 | ***0.0373*** | 0.8440 |
| D20:4-22:6 | 0.1 | 0.0 | 0.1 | 0.0 | 0.92 | 0.4818 | 1.0000 |
| D20:2-22:6/D20:4-22:4 | 0.1 | 0.0 | 0.1 | 0.0 | 1.53 | 0.2418 | 0.9976 |
| P22:6-22:7 | 0.1 | 0.0 | 0.1 | 0.1 | 0.89 | 0.7784 | 1.0000 |
| D22:6-22:4 | 0.4 | 0.1 | 0.3 | 0.2 | 0.69 | 0.1398 | 0.9924 |
| D22:6-24:6 | 0.2 | 0.0 | 0.2 | 0.1 | 0.79 | 0.4265 | 1.0000 |

**Table 8. Absolute Pool Size of PCs in WT and HMGCS2 KO Primary Hepatocytes**

Average and SD for each individual PC species detected via shotgun lipidomics. Fold change is the average of the KO divided by the average of the WT. Significant differences determined by unpaired two-way Student's t test between WT and HMGCS2 KO with Holm-Sidak multi-comparisions correction.

| Shotgun Lipidomics Phosphatidylcholine (PC) Pool Sizes (nmol lipid / mg protein) | | | | | | | |
|---|---|---|---|---|---|---|---|
| **PC Species** | **WT** | | **HMGCS2 KO** | | **Fold Change** | **Raw P Value** | **Adjusted P-Value** |
| | Average | ±SD | Average | ±SD | | | |
| **D16:1-18:2** | 1.3 | 0.3 | 2.0 | 0.4 | 1.55 | ***0.0095*** | *0.0794* |
| **D16:0-18:2** | 31.0 | 5.9 | 35.4 | 11.2 | 1.14 | 0.4377 | 0.9683 |
| **D18:2-18:3/D16:1-20:4** | 0.7 | 0.2 | 0.7 | 0.2 | 0.94 | 0.7103 | 0.9683 |
| **D18:2-18:2/D16:0-20:4** | 14.3 | 1.5 | 7.7 | 2.7 | 0.54 | ***0.0004*** | ***0.0051*** |
| **D18:1-18:2/D16:0-20:3** | 2.4 | 0.2 | 3.3 | 0.4 | 1.34 | ***0.0011*** | ***0.0121*** |
| **D18:2-20:5** | 0.4 | 0.1 | 0.4 | 0.1 | 0.95 | 0.6959 | 0.9683 |
| **D16:0-22:6/D18:2-20:4** | 10.2 | 1.3 | 9.0 | 4.8 | 0.88 | 0.5803 | 0.9683 |
| **D18:1-20:4/D16:0-22:5** | 2.9 | 0.5 | 2.7 | 0.4 | 0.93 | 0.4684 | 0.9683 |
| **D18:2-20:2/D18:0-20:4** | 6.2 | 0.4 | 5.3 | 0.5 | 0.85 | ***0.0098*** | *0.0794* |
| **D18:0-20:3** | 0.5 | 0.1 | 0.4 | 0.1 | 0.84 | 0.2042 | 0.7813 |
| **D18:0-20:2/P18:2-22:6** | 0.4 | 0.1 | 0.3 | 0.1 | 0.91 | 0.5943 | 0.9683 |
| **D18:1-22:6/D18:2-22:5** | 1.3 | 0.3 | 0.8 | 0.2 | 0.63 | ***0.0065*** | *0.0624* |
| **D18:0-22:6** | 2.5 | 0.1 | 1.8 | 0.2 | 0.74 | ***0.0000*** | ***0.0001*** |

**Table 9. Absolute Pool Size of TAGs in WT and HMGCS2 KO Primary Hepatocytes**

Average and SD for each individual TAG species detected via shotgun lipidomics. Fold change is the average of the KO divided by the average of the WT. Significant differences determined by unpaired two-way Student's t test between WT and HMGCS2 KO with Holm-Sidak multi-comparisions correction.

| Shotgun Lipidomics Triacylglycerol (TAG) Pool Sizes (nmol lipid / mg protein) | | | | | | | |
|---|---|---|---|---|---|---|---|
| **TAG Species** | **WT** | | **HMGCS2 KO** | | **Fold Change** | **Raw P Value** | **Adjusted P-Value** |
| | Average | ±SD | Average | ±SD | | | |
| **C50:4** | 0.9 | 0.3 | 2.8 | 0.4 | 2.9 | ***0.0000*** | ***0.0001*** |
| **C50:3/C51:10** | 14.8 | 11.6 | 108.7 | 35.1 | 7.3 | ***0.0001*** | ***0.0021*** |
| **C50:2/C51:9** | 18.4 | 12.6 | 100.8 | 27.8 | 5.5 | ***0.0001*** | ***0.0013*** |
| **C52:6** | 0.8 | 0.2 | 1.7 | 0.2 | 2.1 | ***0.0000*** | ***0.0006*** |
| **C52:5** | 23.9 | 16.4 | 105.2 | 22.9 | 4.4 | ***0.0000*** | ***0.0007*** |
| **C52:4/C53:11** | 88.3 | 54.1 | 247.2 | 45.3 | 2.8 | ***0.0002*** | ***0.0037*** |
| **C52:3/C53:10** | 85.2 | 53.3 | 243.6 | 48.2 | 2.9 | ***0.0003*** | ***0.0043*** |
| **C52:2/C53:9** | 22.8 | 13.9 | 75.1 | 17.5 | 3.3 | ***0.0002*** | ***0.0034*** |
| **C52:1/C53:8** | 1.6 | 0.9 | 4.4 | 1.1 | 2.7 | ***0.0008*** | ***0.0105*** |
| **C53:0/C54:7** | 8.9 | 5.7 | 32.0 | 6.4 | 3.6 | ***0.0001*** | ***0.0011*** |
| **C54:6** | 28.7 | 15.0 | 92.3 | 15.3 | 3.2 | ***0.0000*** | ***0.0005*** |
| **C54:5/C55:12** | 38.8 | 20.3 | 146.3 | 19.6 | 3.8 | ***0.0000*** | ***0.0001*** |
| **C54:4/C55:11** | 26.6 | 13.7 | 121.3 | 15.2 | 4.6 | ***0.0000*** | ***0.0000*** |
| **C54:3/C55:10** | 9.3 | 4.5 | 43.4 | 7.7 | 4.7 | ***0.0000*** | ***0.0001*** |
| **C54:2/C55:9** | 4.0 | 2.0 | 8.8 | 2.7 | 2.2 | ***0.0070*** | ***0.0426*** |
| **C55:2/C56:9** | 2.2 | 1.0 | 4.2 | 0.8 | 2.0 | ***0.0031*** | ***0.0298*** |
| **C55:1/C56:8** | 8.6 | 4.1 | 14.1 | 2.5 | 1.6 | ***0.0205*** | *0.0946* |
| **C55:0/C56:7** | 8.8 | 4.4 | 17.0 | 3.4 | 1.9 | ***0.0052*** | ***0.0409*** |
| **C56:6** | 5.3 | 2.8 | 10.2 | 1.6 | 1.9 | ***0.0037*** | ***0.0322*** |
| **C56:5/C57:12** | 3.5 | 1.8 | 7.3 | 1.0 | 2.1 | ***0.0012*** | ***0.0130*** |
| **C56:4/C57:11** | 3.6 | 2.0 | 8.6 | 1.9 | 2.4 | ***0.0014*** | ***0.0147*** |
| **C56:3/C57:10** | 2.8 | 1.8 | 6.6 | 1.9 | 2.3 | ***0.0061*** | ***0.0409*** |
| **C57:3/C58:10** | 2.1 | 0.8 | 2.4 | 0.4 | 1.1 | 0.4538 | 0.4848 |
| **C57:2/C58:9** | 2.8 | 1.1 | 3.6 | 0.7 | 1.3 | 0.1432 | 0.2508 |
| **C57:1/C58:8** | 2.1 | 0.9 | 3.2 | 0.6 | 1.5 | ***0.0399*** | 0.1231 |
| **C57:0/C58:7/C59:14** | 1.6 | 0.9 | 2.6 | 0.4 | 1.6 | ***0.0325*** | 0.1231 |

**Table 10. qRT-PCR Primer Sequences**

Sequences of primers for qRT-PCR analysis of mRNA transcript abundance, given in the 5` to 3` direction.

| qRT-PCR Primer Sequences | | |
|---|---|---|
| **Gene** | **Forward (5`-->3`)** | **Reverse (5`-->3`)** |
| **Ptprc** | GACAGAGTTAGTGAATGGAGACC | AAAAGTTCGGAGAGTGTAGGC |
| **Emr1** | CTTTGGCTATGGGCTTCCAGTC | GCAAGGAGGACAGAGTTTATCGTG |
| **Tek** | GCAGATTTTGGATTGTCACGAG | GAGCAATACACCATAGGACCAG |
| **Vwf** | TGCGGACATTTTCTCAGACC | TTTTGCTACGGTGAGACAGG |
| **Krt19** | GTTCTCAGACCTGCGTCC | TGACCCAATGCGTACTGAAC |
| **Col1a1** | TGCTTCGTGTAAACTCCCTC | TTGTTCGTCTGTTTCCAGGG |
| **Col1a2** | ATTCGCACCACTTGTGGCTTCT | TGGGATGGTCTACACTCACTGCAA |
| **Trr** | AGGCATCCTTTCCATCTTGG | CTGCTTTCTGACCTATCTTCCTC |
| **Alb** | GACACCTGCTTCTCGACTG | CACCAACAGAAAAGATGAGTCC |
| **Hmgcs2** | TGGTTCAAGACAGGGACACAGAAC | AGAGGAATACCAGGGCCCAACAAT |
| **Bdh1** | TGCAACAGTGAAGAGGTGGAGAAG | CAAACGTTGAGATGCCTGCGTTGT |
| **Oxct1** | TGGCCAACTGGATGATACCTGG | TCCATGGTGACCACCACTTTGG |
| **L32** | CCTCTGGTGAAGCCCAAGATC | TCTGGGTTTCCGCCAGTTT |
